# A common neural signature between genetic and environmental risk

**DOI:** 10.1101/2024.06.13.598859

**Authors:** Maria Vedechkina, Joni Holmes, Varun Warrier, Duncan Astle

## Abstract

Not everyone is equally likely to experience mental illness. What is the contribution of an individual’s genetic background, or experiences of childhood adversity, to that likelihood? And how do these dimensions of risk interact at the level of the brain? We investigated the relationship between genetic liability for mental illness, childhood adversity, and cortico-limbic connectivity in a large developmental sample drawn from the ABCD cohort. First, we used Canonical Correlation Analysis to uncover two genetic dimensions of mental health using polygenic risk scores for ADHD, Anxiety, Depression, and Psychosis. The first dimension represented liability for broad psychopathology which positively correlated with adversity, while the second represented neurodevelopmental-specific risk which negatively interacted with adversity. Next, we investigated the cortico-limbic signature of adversity and genetic liability using Partial Least Squares. We found that the neural correlates of adversity broadly mirrored those of genetic liability, with adversity capturing most of the shared variance. These novel findings suggest that genetic and environmental risk *overlap* in the neural connections that underlie behavioural symptomatology.

## INTRODUCTION

The prevalence of mental illness represents a significant global health challenge, with a myriad of factors contributing to its onset and progression.^1^ Different combinations of interacting genetic, biological and environmental constraints are involved in shaping the neural substrates of mental health.^2^ To better understand the etiology of common mental health conditions, it is therefore necessary to consider both genetic and environmental influences *in combination* and discern how they converge to produce risk-relevant neural and behavioural phenotypes.^3–5^

The human genome interacts with the environmental factors to shape both physical and behavioural traits.^6–9^ The molecular embedding of adversity in stress-regulatory systems suggests that genetic and environmental risk may converge at the level of neurobiology, resulting in comparable changes in neural structure and function.^10–12^ Indeed, emerging evidence suggests that correlations between childhood maltreatment and mental health are partly driven by genetic factors.^13^ Genetic liability for mental illness is believed to manifest by altering the development of neural systems that play a role in symptomology, such as those involved in affective processing, fear learning, and social cognition.^11,12,14,15^ One popular account is that genetic and environmental risk factors may converge at the level of the brain by altering stress-susceptible neural systems in similar ways.^16–18^ Indeed, adversity exposed individuals often exhibit similar changes in brain structure and function to those typically observed in psychiatric patients, notably in the cortico-limbic connections.^19–23^ These neural differences may serve as partially heritable intermediate processes that mediate the link from genetic and environmental risk to more complex behavioural outcomes.^24–26^

What is the contribution of an individual’s genetic background or experience of early life adversity to mental illness? How do genetic and environmental influences shape risk-relevant neural and behavioural phenotypes? And, crucially, to what extent do these dimensions of risk overlap at the level of the brain? This study investigates relationships between genetic liability for mental illness, childhood adversity and functional connectivity in a large sample of US children aged 9-10 drawn from the Adolescent Brain and Cognitive Development study (ABCD).

First, we estimate polygenic risk scores (PRS) for four mental health phenotypes: ADHD, Anxiety, Depression, and Psychosis. While the classification of psychiatric disorders has historically segmented mental health conditions into distinct categories, recent genetic studies have challenged the assumptions of disorder-specific risk by identifying a large number of *common* genetic variants with small effects, many of which predispose to *multiple* mental disorders ^27–29^ and by demonstrating a high degree of genetic correlation across different conditions.^30,31^ We, therefore, use Canonical Correlation Analysis (CCA) to assess the degree of shared genetic liability across the four mental health conditions and uncover overlapping dimensions of genetic risk that transcend traditional phenotypic boundaries. Next, we examine how these genetic liability dimensions relate to history of adversity. Finally, we use Partial Least Squares (PLS) to investigate the cortico-limbic signature of genetic and environment risk and assess whether these neural differences mediate mental health.

## RESULTS

### Polygenic risk scores

GWAS summary statistics from four mental health disorders (ADHD; Anxiety; Depression; Psychosis) were used to calculate PRS scores for four mental health phenotypes in the ABCD cohort for 6,535 participants who passed all filters and QC steps (Figures S1-S6; Table S3). Hierarchical linear regressions assessing how much variance in each mental health phenotype is explained by each of the PRS scores, with age, sex, and the first 6 population substructure PCs as covariates are reported in Tables S4-S7. All four PRS scores were significantly correlated with their respective mental health phenotype and with adversity (Figure 1).

**Fig 1.**
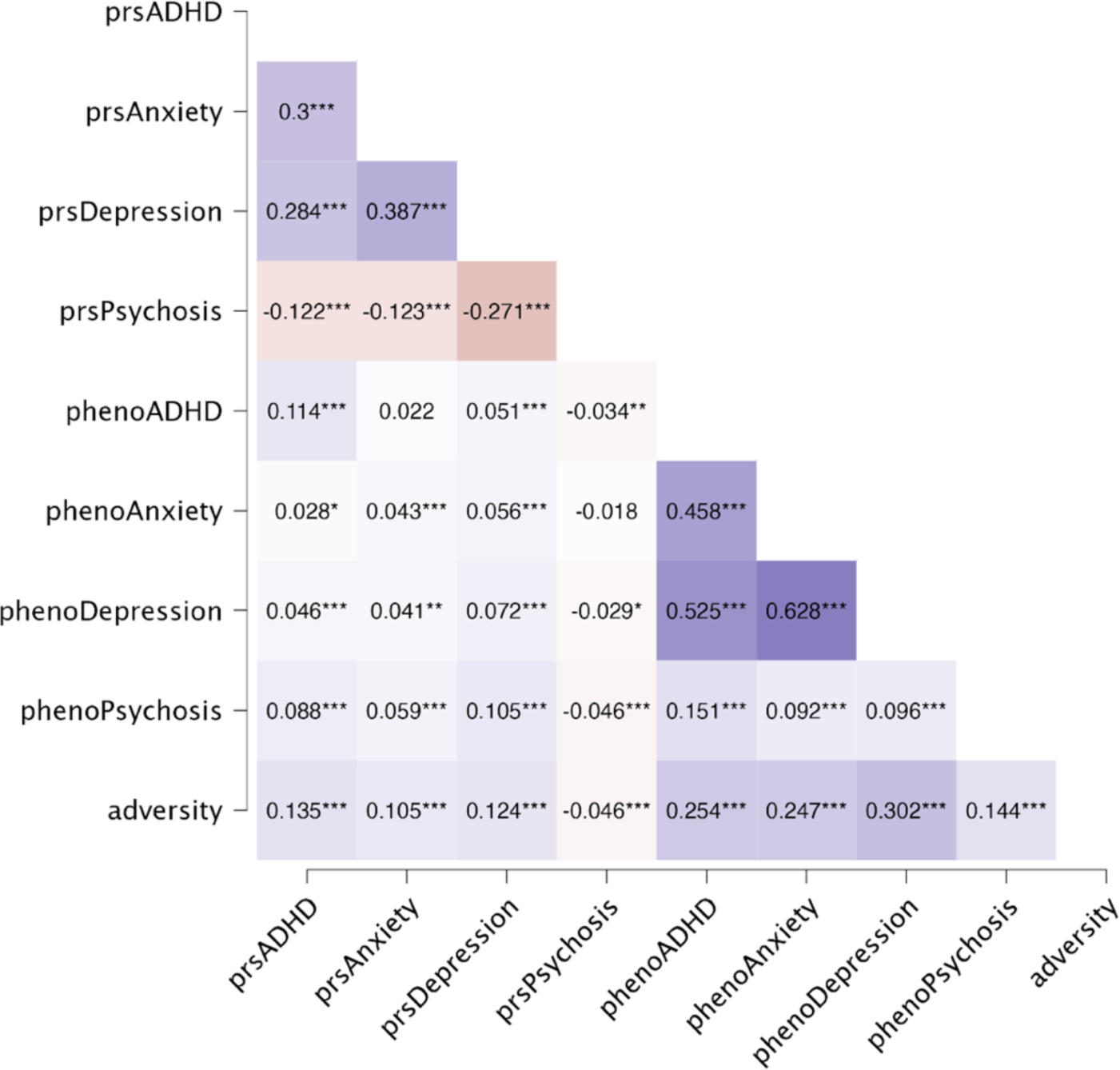
Correlations between genotype scores, their respective phenotypes, and adversity. * *p* < .017, ** *p* < .01, *** *p* < .001.

### Shared dimensions of genetic risk

Canonical Correlation Analysis (CCA) was used to explore the underlying relationships between the four PRS scores (Set A) and their respective phenotypes (Set B) and identify possible genetic and phenotypic overlap across the four mental health conditions. A CCA model was fitted to identify pairs of canonical variates across two sets of variables: Set A (prsADHD, prsAnxiety, prsDepression, prsPsychosis) and Set B (phenoADHD, phenoAnxiety, phenoDepression, phenoPsychosis). The CCA yielded two significant canonical variates with the following canonical correlation coefficients: Canonical Variate 1 (*r*= 0.15; Wilks’ λ = .97, F (16, 19950.12) = 13.23, *p* < .001); Canonical Variate 2 (*r*= 0.09; Wilks’ λ = .99, F (9, 15894.89) = 6.43, *p* < .001; Figure S7). The canonical loadings showing the corresponding strength and significance of correlations between disorder-specific mental health variables and each of the obtained genetic and phenotypic component scores are shown in Figure 2 and Table S8. These loadings suggest that the first canonical variate can be thought of as representing general risk for psychopathology across multiple conditions (henceforth referred to as *CVgeneral*), whereas the second canonical variate represents unique neurodevelopmental-specific variance, characterised by positive loadings only from prsADHD (henceforth referred to as *CVneurodev*).

**Fig 2.**
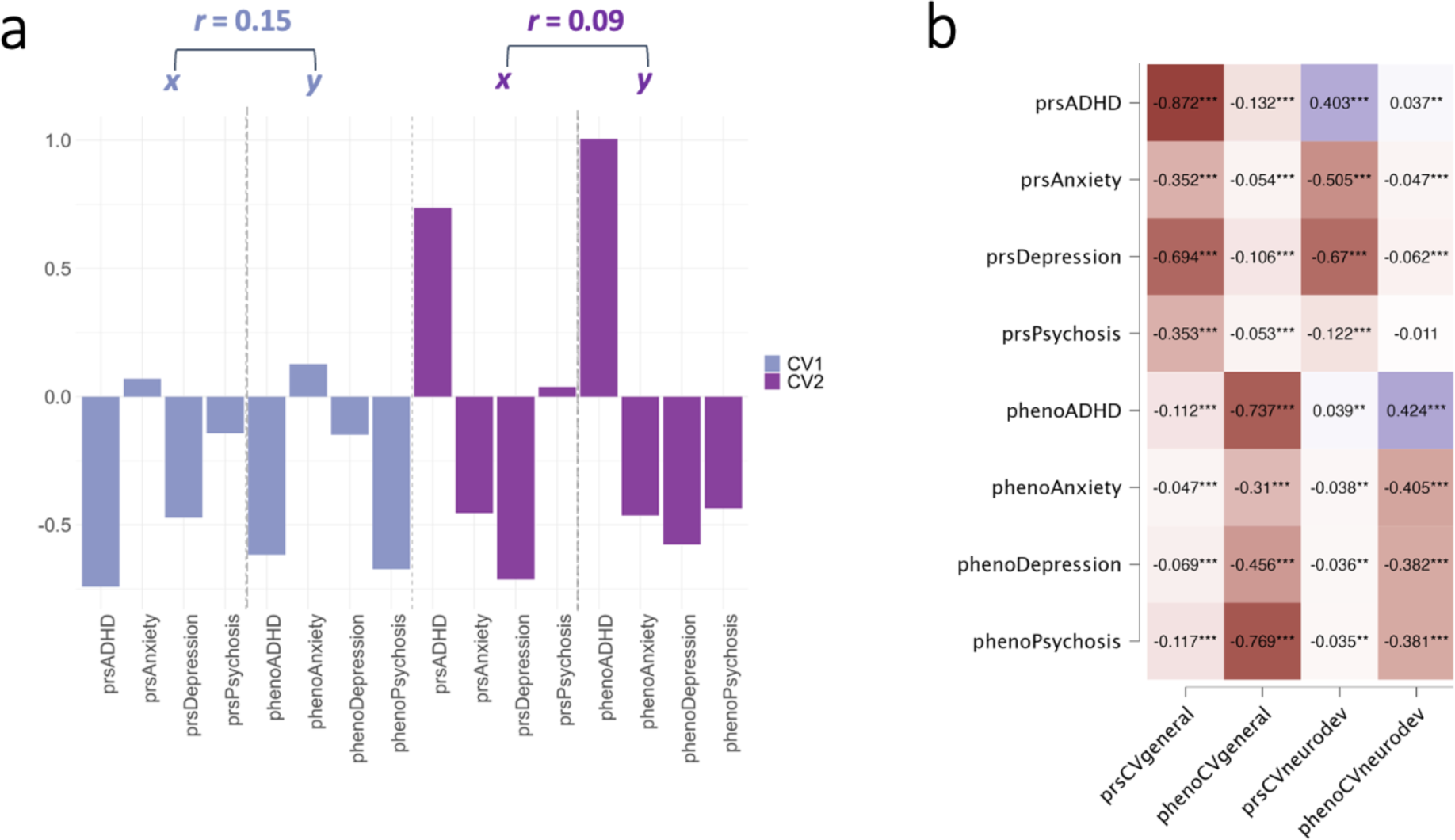
Canonical coefficients and canonical loadings for PRS scores and mental health phenotypes. (A) Scaled canonical correlation coefficients for each canonical variate. r= canonical correlation between each set of canonical components. (B) Canonical loadings showing the strength and significance of correlation of all variables with each canonical component. prsCVgeneral= PRS component scores from the first canonical variate representing general psychopathology. phenoCVgeneral= Phenotype component scores from the first canonical variate representing general psychopathology. prsCVneurodev= PRS component scores from the second canonical variate representing neurodevelopmental-specific variance. phenoCVneurodev= Phenotype component scores from the second canonical variate representing neurodevelopmental-specific variance. * *p* < .05, ** *p* < .01, *** *p* < .001.

Multiple linear regressions demonstrating the amount variance in each mental health phenotype explained by the two PRS components (prsCVgeneral & prsCVneurodev), controlling for age, sex, and the first 6 population substructure PCs and comparing these results to the predictive value of the disorder-specific PRS scores are reported in in Tables S9-S12. The results did not qualitatively change when including population substructure PCs as covariates for the entire sample (Tables S34-S36) and for individuals with European ancestry only (Tables S37-S39). Overall, this suggests that genetic liability for different mental health conditions can be reduced to two independent transdiagnostic dimensions to more powerfully predict variance in the respective behavioural phenotypes.

### Relationship between genetic liability for mental-ill health and adversity

To investigate whether genetic liability for mental illness is related to adversity, multiple linear regressions were conducted with prsCVgeneral and prsCVneurodev as predictors controlling for the respective CVpheno scores, age, sex, and the 6 PCs. prsCVgeneral (*b*=.089*, p*<.001), but not prsCVneurodev (*b*=.000, *p*=0.99), was positively related to adversity (Tables S13-S14). prsCVgeneral (*b*=.116*, p*<.001) and adversity (*b*=.243*, p*<.001) both independently predicted phenoCVgeneral, but there was no interaction effect (Figure 3A & C; full results in Table S19). In contrast, prsCVneurodev (*b*=.084*, p*<.001) and adversity (*b*=-.098*, p*<.001) both independently predicted phenoCVneurodev— with similar strength but with opposing directions of effect— and there was also a significant interaction between prsCVneurodev and adversity (*b*=.043*, R*² *change*=.001, *p*=.006), indicating that increases in the effect of either variable attenuate the influence of the other (Figure 3B & D; Table S20). In other words, the effect of environmental adversity is mitigated with increased neurodevelopmental genetic liability.

**Fig 3.**
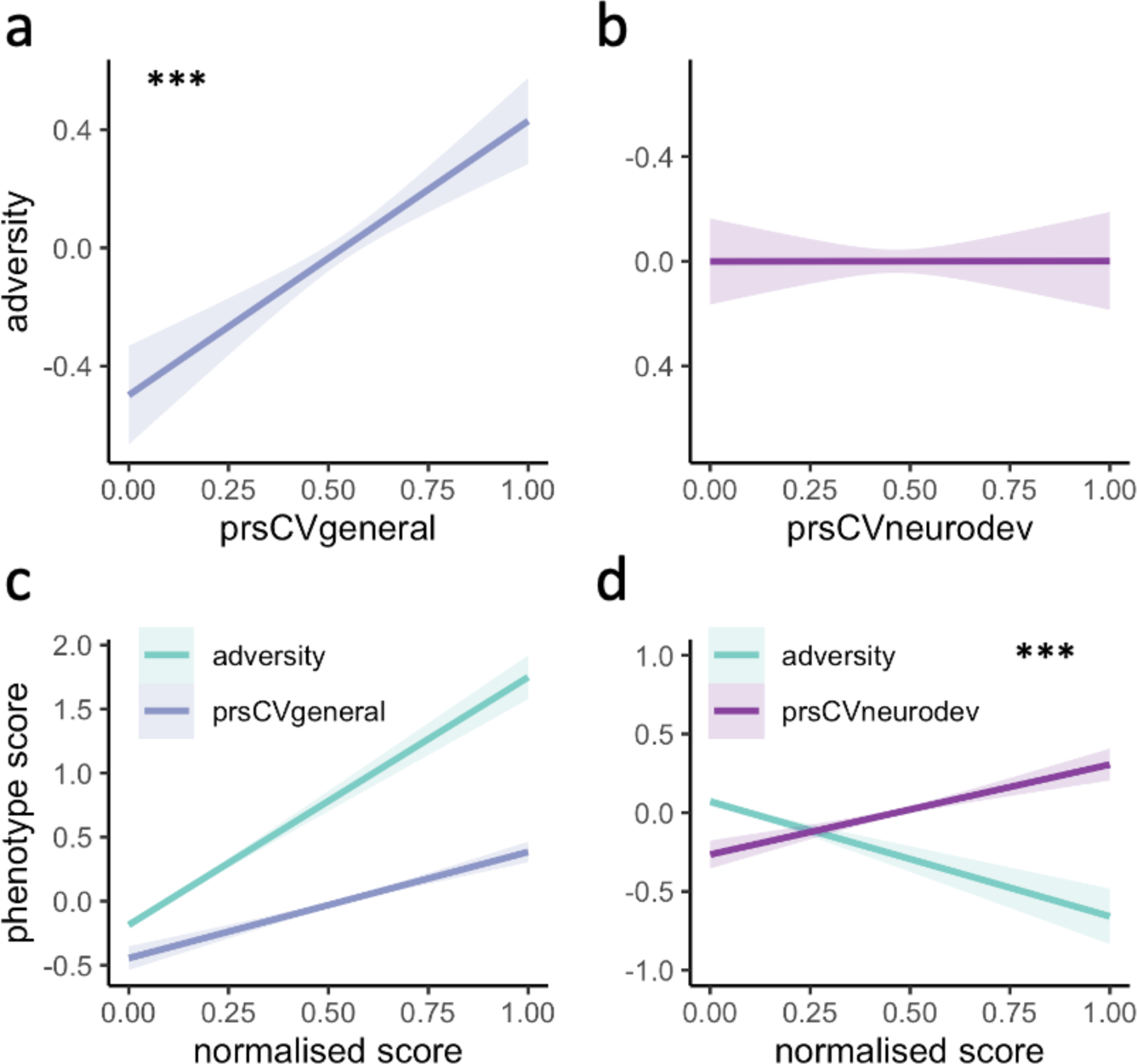
Correlation and interaction between genetic liability dimensions and adversity. (A-B) Association between genetic liability components and adversity. prsCVgeneral= PRS component scores from the first canonical variate representing general psychopathology. prsCVneurodev= PRS component scores from the first canonical variate representing neurodevelopmental-specific variance. Regression models include age, sex, the first 6 population components (PCs), and the respective phenotype measures as covariates. (C-D) Prediction of phenotype components by PRS components, adversity, and their interaction. Regressions model the predictive value for each phenotype component separately, including age, sex, and the first 6 population components (PCs) as covariates. *** *p* < .001.

### Cortico-limbic signature of genetic and environmental risk

In our final analyses, we leveraged Partial Least Squares (PLS) to uncover the neural signature of genetic and environmental risk and assess the extent of overlap between the two dimensions of risk in the connections between the limbic and cortical networks.

#### Cortico-limbic global connectivity

First, PLS was used to model the covariance between genetic/environmental risk and connectivity between the entire limbic network (n=1) and cortical networks (n=12). When using the four PRSs and adversity as joint predictors (joint model), a single significant component emerged (*p*perm< .001) and explained 33% of the variance in the predictors and 43% of the variance in the cortico-limbic network connections (Table S25). The correlation between the latent components was *b*= 0.03, SE= 0.009, *p*<.001 when controlling for age, sex and the 6PCs, and remained significant when mental health phenotype scores were added as covariates (*b*= 0.03, SE= 0.009, *p*< .001).

When using only the four PRSs as predictor variables (PRS-only model), a single significant component emerged (*p*perm<.001) and explained 42% of the variance in the PRSs and 43% of the variance in the cortico-limbic network connections (Figure 4; Table S26). The correlation between the latent components was *b*= 0.07, SE= .034, *p*=0.04 when controlling for age, sex and the 6PCs, and remained significant when mental health phenotype scores were added as covariates (*b*= 0.07, SE= 0.034, *p*=0.05). However, the CCA model became non-significant when adversity was added as a covariate (*b*= 0.05, SE= 0.034, *p*=0.11).

**Fig 4.**
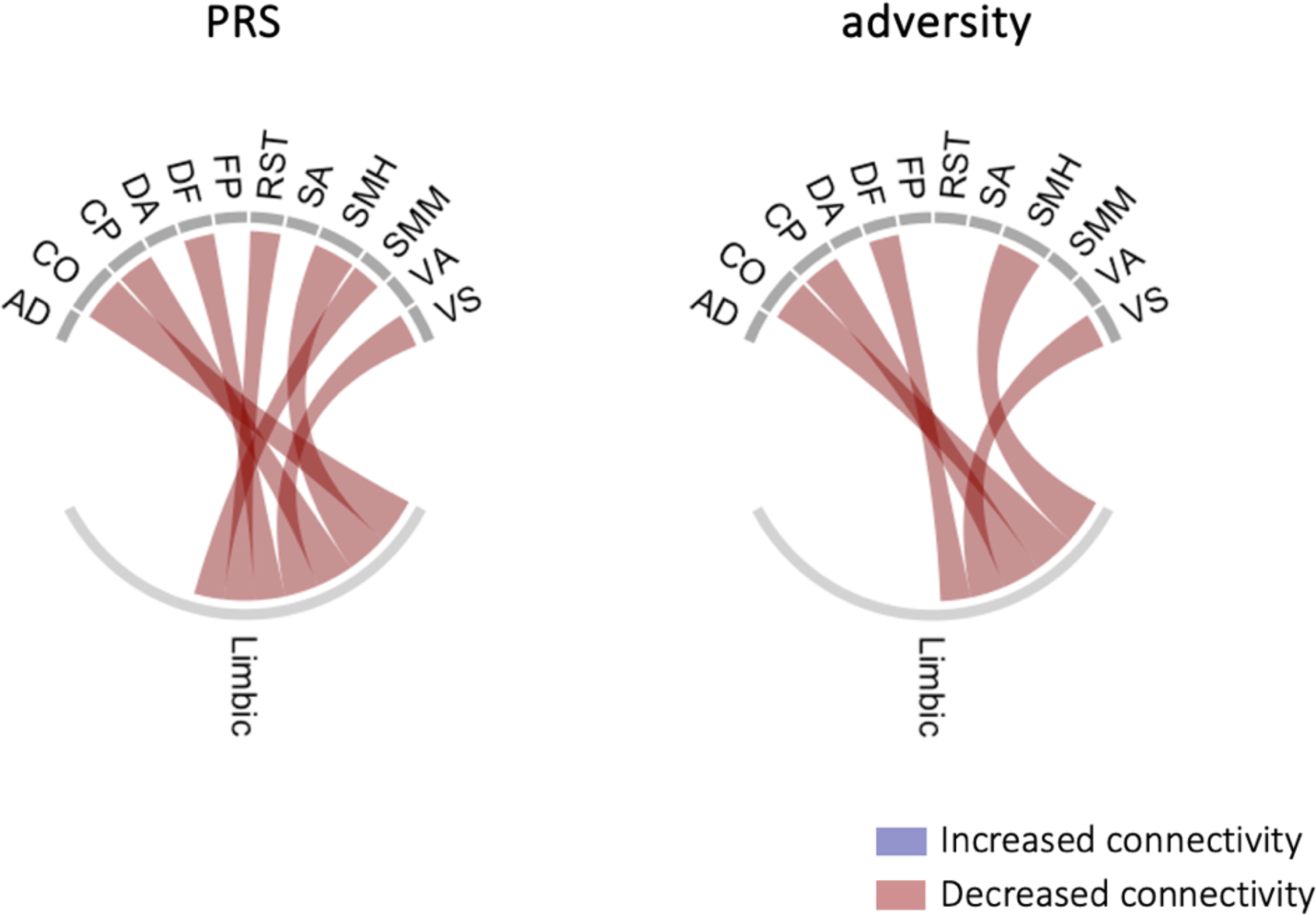
PLS loadings of genetic risk and adversity on cortico-limbic global connectivity. PLS loadings of cortico-limbic network connectivity on the PRS-only model and adversity-only model. Only significant loadings with a VIP score above 1 shown. AD= auditory; CO= cingulo-opercular; CP= cingulo-parietal; DA= dorsal attention; DF= default; FP= frontoparietal; RST= retrosplenial temporal; SA= salience; SMH= sensorimotor hand; SMM= sensorimotor mouth; VA= ventral attention; VS= visual.

When using only cumulative adversity as the predictor (adversity-only model), a single significant component emerged (*p*perm< .001) and explained 100% of the variance in adversity and 42% of the variance in the cortico-limbic network connections (Figure 4; Table S27). The correlation between the latent components was *b*= 0.03, SE= 0.008, *p*<.001 when controlling for age, sex and the 6PCs, and remained significant when mental health phenotypes scores (*b*= 0.03, SE= 0.007, *p*<.001) and the PRS scores (*b*= 0.03, SE= 0.007, *p*<.001) were added as covariates. The cortico-limbic network component significantly mediated the association between the adversity component and symptoms of depression and psychosis (*p*’s= .032-.033; Table S28).

Together, these analyses suggest that while adversity and the four genetic liability factors map onto common reductions in cortico-limbic connectivity at the global level, adversity captures much of this shared variance. Moreover, the mediation analysis underscores the role of cortico-limbic circuitry as a potential neural pathway through which these risk factors may influence clinical symptomatology.

#### Cortico-limbic regional connectivity

A second PLS was used to model the covariance between genetic/environmental risk and connectivity between individual limbic regions (n=19) and cortical networks (n=12). When using the PRSs and adversity as joint predictors (joint model), a single significant component emerged (*p*perm< .001), explaining 33% of the variance in the predictors and 10% of the variance in the cortico-limbic regional connections. There were 70 significant regional connection loadings out of a total of 228 (Table S29). The correlation between the latent components was *b*= 0.022, SE= 0.04, *p*< .001 when controlling for age, sex and the 6PCs, and remained significant when mental health phenotype scores were added as covariates (*b*= 0.021, SE= 0.004, *p*< .001).

When using only the PRSs as predictor variables (PRS-only model), a single significant component emerged (*p*perm< .001), explaning 42% of the variance in the PRSs and 10% of the variance in the cortico-limbic regional connections. There were 73 significant connections out of a total of 228 (Figure 5; Table S30). The correlation between the latent components was *b*= 0.015, SE= 0.005, *p*<.001 when controlling for age, sex and the 6PCs, and remained significant when mental health phenotype scores (*b*= 0.014, SE= 0.005 *p*=.001) and adversity (*b*= 0.013, SE= 0.005, *p*=.005) were added as covariates. The cortico-limbic regional component significantly mediated the association between the PRS component and symptoms of psychosis (*b*= −0.021, *p*= .038; Table S31).

**Fig 5.**
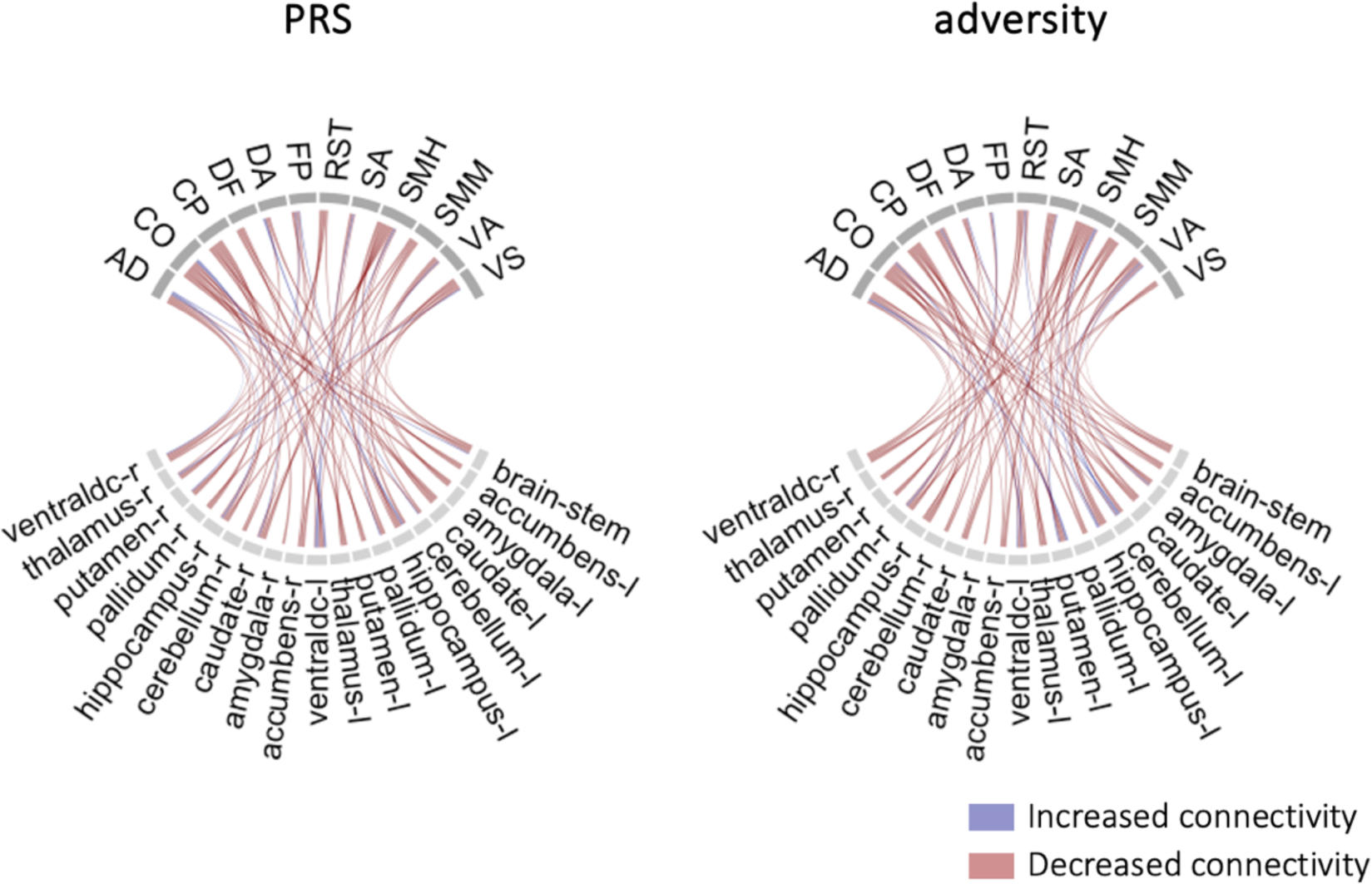
PLS loadings of genetic risk and adversity on cortico-limbic regional connectivity. PLS loadings of cortico-limbic regional connectivity on the PRS-only model and adversity-only model. Only significant loadings with a VIP score above 1 shown. AD= auditory; CO= cingulo-opercular; CP= cingulo-parietal; DA= dorsal attention; DF= default; FP= frontoparietal; RST= retrosplenial temporal; SA= salience; SMH= sensorimotor hand; SMM= sensorimotor mouth; VA= ventral attention; VS= visual. The PRS and adversity models differ on 41 out of 147 significant loadings.

When using only cumulative adversity as the predictor (adversity-only model), a single significant component emerged (*p*perm< .001). There were 74 significant regional connections out of a total of 228 (Figure 5; Table S32). The adversity-only model differed from the PRS-only model on 41 out of 147 significant loadings. Notably, the PRS-only model had greater loadings of limbic connectivity with the frontoparietal and visual networks, while the adversity-only model had greater loadings of limbic connectivity with the salience network. In contrast, there were few differences between the two models in terms of limbic connectivity with the cingulo-opercular, cingulo-parietal, and sensorimotor mouth networks. The correlation between the latent components was *b*= 0.023, SE= 0.004, *p*<.001 when controlling for age, sex and the 6PCs, and remained significant when mental health phenotypes scores (*b*= 0.023, SE= 0.004, *p*<.001) and the PRS scores (*b*= 0.021, SE=0.004, *p*<.001) were added as covariates. The cortico-limbic regional component significantly mediated the association between the adversity component and symptoms of psychosis (*b*= −0.042, *p*= .006; Table S33). Together, these analyses suggest that while adversity and the four genetic liability factors broadly correspond to similar regional differences in cortico-limbic connectivity, distinct patterns of connectivity associated with each dimension of risk become evident at the regional level of analysis. Moreover, the mediation analysis underscores the role of cortico-limbic circuitry as a potential neural pathway through which both genetic and environmental risk factors may independently influence clinical symptomatology.

## DISCUSSION

We investigated relationships between polygenic risk for mental health, childhood adversity, and cortico-limbic connectivity in a large developmental sample drawn from the ABCD study. This study uncovers two novel findings. First, that genetic liability for ADHD, anxiety, depression and psychosis can be effectively reduced to two pleotropic dimensions of risk, each of which uniquely interacts with exposure to adversity. Second, that genetic and environmental risk factors *overlap* in their neural correlates. Indeed, we identify a common neural signature between adversity and the four genetic risk scores, indicative of gene-environment correlations at the level of the brain. Together, our findings suggest that environmental adversity and genetic predispositions can have a compounding and interactive effect on neural connections that underlie behavioural symptom variance.

The first finding from this study highlights that current diagnostic classifications, which are largely based on clinical observation, do not reflect the underlying pathophysiology of mental health. These findings add to a growing body of evidence highlighting the lack of direct mapping of traditional psychiatric disorder categories with underlying neurobiological processes,^28,35,36^ and demonstrating links between genetic liability for psychopathology and a variety of environmental and socio-demographic exposures.^13,37,38^ It is likely that the same genetic variants are expressed differently across different tissues and different stages of development, leading to slight alterations in the behavioural presentation of similar genetic factors.^32^

Relatedly, environmental influences and modifier genes may moderate how genetic variants are expressed.^33^ Indeed, we show that genetic liability for general psychopathology and cumulative adversity are correlated and independently predict mental health symptoms, suggestive of an additive effect,^34^ while neurodevelopmental-specific liability negatively interacts with environmental risk, indicating that the influence of adversity on behavioural symptoms is attenuated with increasing neurodevelopmental-specific genetic risk. This points to the possibility that neurodevelopmental symptoms may arise from unique combinations of genetic and environmental factors that differ from other symptom domains. Given the pervasive co-occurrence between disorder categories and growing evidence of common genetic origins,^29,30,39^ disorder-specific GWAS studies may be in need of re-evaluation. A shift towards transdiagnostic models that better reflect the underlying pathophysiology of different symptom dimensions would likely provide valuable insights into the etiology of psychiatric conditions and improve clinical practice. Second, this study demonstrates that both adversity and genetic liability are associated with reduced functional connectivity between the limbic and several cortical networks, including the the cingulo-opercular, cingulo-parietal, visual, default, and sensorimotor networks. This findings aligns with the dysconnectivity hypothesis, which posits that disruptions in functional connectivity are a core feature of many mental health conditions.^40–42^ Our results suggest that *widespread* differences in cortico-limbic connectivity likely contribute mental health symptomatology, lending support to the idea that genetic and environmental influences are unlikely to be confined to specific neural circuits.^43^ A particularly novel finding of this study is the substantial overlap in the neural signatures of genetic and environmental risk, indicating a gene-environment correlation at the level of the brain. Notably, our results reveal that adversity accounts for the entirety of the genetic variance at the global level. This suggests that adversity may play a prepotent role in shaping the functional connections between limbic and cortical networks, beyond the influence of genetic factors alone.

While our global analyses revealed an overlapping pattern of reduced connectivity between the limbic and cortical networks, our regional analyses highlight some degree of heterogeneity *within* the limbic network in relation to genetic and environmental risk. The regional model identified 41 out of 147 cortico-limbic connections that uniquely associated with genetic risk. The most pronounced differences between genetic and environmental risk were in the connections with the auditory, fronto-parietal, ventral attention, sensorimotor hand, and visual networks. Furthermore, regional connectivity with the fronto-parietal and visual networks showed the strongest relationship to genetic liability, suggesting that these circuits may be particularly sensitive to genetic influences.^45,46^ This lends support to the idea that pleotropic risk genes increase susceptibility to a variety of clinically-distinct psychiatric conditions through *non-specific* changes in cortico-limbic connections.^47^

Together, this study demonstrates that the neural correlates of adversity largely mirror those of genetic liability for mental health, with some notable regional differences. In other words, comparable neural differences can emerge from various endogenous (genetic) and exogenous (environmental) processes. This suggests that many of the previously identified neural markers of psychopathology may actually be capturing adversity-related variance, or vice-vesa. In other words, the neural features stemming from early adversity are likely conflated with those that predispose individuals to mental illness in the existing literature.^20^ Indeed, most adversity studies do not control for co-occuring psychiatric symptoms. Likewise, most psychiatric studies fail to consider history of adversity. This complicates the task of discerning whether the observed neural differences are driven by exposure to early adversity, mental health symptoms, or a combination of both. Efforts to identify genetically driven neural endophenotypes of mental health^48^ would do well to recognise this as a serious limitation. Future research would benefit from stratifying mental health conditions based not only on clinical symptoms and neurobiological features, but also on genetic factors *and* adversity. This may help pave the way for the identification of clinical subtypes to inform the development of new targeted treatments^49–51^ that are more or less effective for certain groups.^52–54^

The strengths of this study lie in its application of diverse methods to capture multicollinear relationships between genetic liability, environmental risk, and functional brain connectivity. However, some limitations must also be acknowledged. First, the study sample is representative of the US context and was restricted to children aged 9-10, limiting the generalisability of the findings to other demographic groups and age ranges. The role of genetic liability factors is known to evolve over the course of an individual’s life.^55^ Additionally, the ongoing maturation of the neural connections^56,57^ suggests that new neural signatures of mental health may emerge in later life. Future longitudinal research should endeavour to replicate these findings within diverse age groups across development. Second, while the behavioural expression of mental health difficulties may vary at different developmental time points,^58,59^ this study only considered four mental health conditions due to data availability constraints. Third, while the use of dimensional symptom scales circumvents some shortcomings inherent to clinical diagnostic categories,^60^ it is worth noting that only a small fraction of the sample met the threshold for a clinical diagnosis. Indeed, the stronger association of adversity with functional connectivity in our study may reflect the relatively low behavioural symptom variance explained by the PRS scores. Relatedly, it will be necessary to validate these findings using genetic methods better suited to capturing shared genetic signals across multiple conditions.^61^ Finally, additional research is needed to determine whether different forms of adversity have unique associations with genetic risk at the neural level.

## METHODS

### Participants

The Adolescent Brain Cognitive Development (ABCD) study is a longitudinal cohort that involves 21 data acquisition sites across the US and follows over 11,000 children aged 9-10 for 10 years into early adulthood.^62^ The study was designed to approximate the socio-demographic distribution of US children in this age group (Table S1). Participants were required to be aged 9-10 years at baseline for inclusion in the study. Those who lacked English language proficiency; suffered from severe sensory, intellectual, medical or neurological issues; or were unable to participate in MRI scanning were excluded. Parents of participants were required to have either English or Spanish proficiency. Recruitment details and data-collection procedures are described by Garavan et al., 2018. The current study uses data from participants with available genotype (n=6,535) and phenotype data as described below.

### Mental health measures

The parent-reported Child Behaviour Checklist (CBCL) is made up of 113 items rated on a three-point scale (not true; somewhat or sometimes true; very often or always true; Achenbach, 2011). These items are then summed into several subscales. This study uses the three DSM-oriented scales from the CBCL that align with clinical disorder definitions: Attention-Deficit Hyperactivity Disorder (ADHD), Depressive Disorder, and Anxiety Disorder. The scales have good inter-interviewer and test-retest reliability.^64^ Raw scores from the baseline assessment (T1), which capture the number of total questions endorsed, were used (range: 0-20). Additionally, the Prodromal Psychosis Scale-Brief Child Version (PPS) was used to measure psychotic symptoms.^65^ The original screening questionnaire, developed for adolescents and adults, was modified for use with children.^66^ The raw score from T1, based on the number of total questions endorsed, was used (range: 0-117).

### Early life adversity

A total of 24 questions were used to assess whether a child had been exposed to an adverse experience before the age of 10 (Table S2). These were taken from the Demographics Survey; Family History Assessment; Neighbourhood Safety/Crime Survey; PTSD Module; and the Family Environment Scale. Questions were designed to capture experiences from birth up to the baseline assessment, except for the material deprivation questions that ask about experiences in the past 12 months (e.g., ‘In the past 12 months, has there been a time when…’). Caregiver reports were used due to the age of the participants. A total of 21 questions provided binary scores such that an adverse experience was coded as present or not (1 or 0 respectively). The remaining three questions question were Likert-type scales (range: 0-5), which were coded as 1 (adversity present) if the parent ‘strongly agreed/strongly disagreed’ depending on the valence of the statement. The 24 questions were summed to create a cumulative adversity score that represents the total number of adverse experiences endorsed ^67^.

Participants missing more than 15% of data on the adversity measures were removed from the analysis (n=314) and the remaining missing answers were coded as ‘0’ (i.e., adversity not endorsed). These responses were coded as 0 because sensitivity analyses (reported in Supplement 1.2) revealed that either coding the missing responses as 1 (endorsing adversity) or using imputation resulted in estimates of adversity that were significantly higher than population prevalence estimates,^68–70^ indicating both approaches were heavily biased. Moreover, imputation was not appropriate for the adversity data as it was both non-binary and not missing at random (see Supplement for details).

### MRI acquisition and preprocessing

ABCD standard imaging protocols for resting-state functional MRI (rsfMRI) including acquisition, processing, and quality assurance procedures have been described in detail elsewhere.^71,72^ Briefly, the preprocessing pipeline includes within- and between-scan head-motion correction, distortion corrections, removal of initial frames, normalisation, demeaning, regression, and temporal filtering.^73,74^ Average time courses for each region of interest (ROI) were calculated using FreeSurfer’s automated brain segmentation (aseg) and resampled to align with voxels from the fMRI data. Motion time courses are adjusted to account for signals linked to respiration (Hagler et al., 2019). This study uses imaging data from a subset of participants with 10 minutes of valid rsfMRI data below a framewise displacement threshold of 0.2mm collected at T1 (n= 5,995).

Using the processed rsfMRI data, parcellated time series were computed using a seed-based correlational approach.^75^ Regions of interest (ROIs) were defined using the functional Gordon atlas template which comprises 352 ROIs (333 cortical) belonging to one of 13 networks (12 cortical and 1 subcortical).^76^ The functional connectivity between any two ROIs was estimated by calculating the lag-zero Pearson correlation coefficient of parcellated time series. This produced an ROI x ROI correlation matrix for each participant, which underwent an additional variance stabilization procedure using a Fischer z-transform.^77^

### Polygenic risk score calculation

Saliva and blood samples were collected from participants as part of the ABCD biospecimen collection.^78^ DNA extraction, basic biospecimen quality control, and genotyping using the Affymetrix Axiom Smokescreen Array were performed at the Rutgers University Cell and DNA Repository. Genotypes were called from the raw intensities using the Affymetrix Power Tools and the Affymetrix Best Practice Workflow in batch processing. The genomic coordinates were aligned with genome build hg19. To ensure data quality, cohort-level quality control (QC) procedures were implemented, which are described in detail elsewhere.^9^ Samples missing more than 20% on genotype calls and variants with more than 10% missing rates were excluded during the initial QC. Samples with excessive relatedness were also removed. Individuals with sex discrepancies, such as those with ambiguous gender calls, were removed. Missing genotype data was inputted using the TOPMED reference panel with genome build GRCh38 and aligned with genome build hg19.^79^

After obtaining the minimally QCed genomic data, individuals from axiom plate 461, which was identified as problematic, were removed as recommended by the ABCD guidelines. Cryptic relatedness among samples was assessed, and one member of each first- and second-degree related pair was excluded to avoid confounding effects. Ambiguous SNPs and those with a missing allele were removed. PLINK 2.0^80^ was used for QC steps. Genetic variants with a call rate below 90% and minor allele frequency (MAF) less than 0.01 were removed to retain only common variants. Variants showing a significant departure from Hardy-Weinberg equilibrium (p < 1e-6) were filtered out to mitigate potential genotyping errors. Genetically related individuals with a first or second degree relative were removed. Pruning was performed to remove any highly correlated SNPs. Principal component analysis (PCA) was performed on the genotyping data to identify genetically inferred population ancestries to be used as covariates in the PRS calculation. Individuals with a heterozygosity rate of F>3sd from the population mean were removed. Duplicate SNPs and samples with a mislabelled sex chromosome were removed. A total of 6,535 individuals passed all filters and QC steps.

Polygenic risk scores (PRS) were calculated for each of the four mental health phenotypes (see ‘Mental health measures’ above) using PRSice-2 software^81^ based on a set of selected variants selected from the following case-control GWAS studies: (1) ADHD^82^; (2) Anxiety^83^; (3) Depression^84^; (4) Schizophrenia (for psychotic symptoms)^85^. Each of the studies defined cases based on clinical criteria, and controls were selected from individuals without the respective mental disorder. Variants reaching genome-wide significance (*p*<5e-8) and those showing suggestive associations (*p*<1e-5) were included in the PRS calculation.^86^ PRSice-2 was run with the recommended settings, such as clumping and LD pruning thresholds, to control for linkage disequilibrium (LD) between selected variants.^81^ Effect sizes and their standard errors for the selected variants were extracted from the GWAS summary statistics from each of the four studies. The PRS scores were calculated by summing the products of the effect sizes and the number of risk alleles carried by each individual at the selected variants for each of the four mental health phenotypes. The PRS calculation was also repeated separately for individuals from genetically inferred European (n= 5679) and non-European (n=865) ancestries for sensitivity analyses. Descriptive statistics for study variables are shown in Table S40.

### Statistical analysis

#### Identifying shared dimensions of genetic risk

Canonical Correlation Analysis (CCA) was performed using the ‘CCA’ package in R ^87^ to explore the underlying relationships between the four PRS scores (Set A) and their respective phenotypes (Set B) and to identify patterns of correlation across the two datasets. CCA is a multivariate method that is suitable for uncovering complex linear associations between two sets of data. The goal of this approach is to identify pairs of canonical components, or variates, that maximize the correlation between the two sets of variables while being orthogonal to each other.^88^ In this study, we used CCA as a data-reduction technique to identify possible genetic and phenotypic overlap across the four mental health conditions of interest and to identify linked dimensions of mental health. By extracting pairs of PRS components and phenotype components that are maximally correlated, while being uncorrelated with any other components extracted, CCA provides the ability to assess the extent of shared genetic variance across different mental health phenotypes and to test whether a single genetic component is sufficient for explaining a wide range of mental health conditions, or if multiple dimensions of genetic risk are needed. In contrast to other dimensionality-reduction techniques, CCA computes the components in the predictors and outcomes simultaneously such that they are maximally correlated with each other.^89^

In other words, it extracts the underlying structure *between* the two sets of variables, rather than optimising the variance within each set of variables first. As a result, CCA is well-suited to explaining associations between different PRS scores in terms of how they map onto different mental health symptoms.

PRS component scores are calculated from a weighted sum of the individual PRS scores for each mental health dimension, and phenotype component scores are calculated from a weighted sum of the four mental health phenotypes. The first and second canonical correlations denote the associations between the corresponding first and second variate pairs, ordered by decreasing correlation strength. To assess the statistical significance of the canonical correlations between each set of components, p-values are generated using F-approximations of different test statistics. Canonical coefficients (or weights) describe which variables most strongly influence each variate. Variables with larger coefficients have a greater relative influence on the overall variate score. A series of correlation analyses were conducted to assess the strength and significance of the association between each of the variables in the model and the component scores in a way that is easier to interpret than the canonical weights alone. As a sensitivity check to assess if our results were driven by confounding genetic population stratification, we conducted supplementary CCA analyses that included population substructure PCs as covariates for the entire sample and for individuals with European ancestries only.

Finally, multiple linear regressions were conducted to assess how much variance in each mental health phenotype is explained by each of the PRS components, controlling age, sex, and the first 6 population substructure PCs. This allowed us to assess whether the PRS components explain more (or less) variance than disorder-specific PRS scores for each mental health phenotype. Because CCA variates are orthogonal to each other, meaning that each additional variate explains only the remaining variance not accounted for by previous components, all PRS component scores were entered jointly as predictors in the regression models. Confidence intervals for the multiple regressions were generated using bootstrapping with 1000 replicates.

#### Relationship between genetic liability for mental-ill health and adversity

To assess possible overlapping liability between genetic risk and adversity, multiple linear regressions were performed to examine the unique contribution of each PRS component score to variation in cumulative adversity, controlling for the respective phenotype component score, age, sex, and the first 6 population substructure principle components (PCs). Genetic population PCs were included as covariates to mitigate spurious associations between ancestry-based population-level stratification and outcomes of interest. For instance, studies have demonstrated correlations between variation in population structure and certain brain imaging features, as well as the prevalence of poverty in the US.^90,91^ Failure to control for genetically-defined population structure can therefore in spurious associations between brain measures, mental health, and adversity driven by uncorrected for population stratification within a given sample.

Next, multiple hierarchical linear regressions were conducted to assess whether PRS and adversity both independently predict mental health phenotypes and whether there is an interaction effect between genetic risk and the environment. Cumulative adversity scores were entered alongside each PRS component as predictors for the respective phenotype component, controlling age, sex, and the first 6 population substructure PCs. Interaction terms between PRS variates and adversity were added to the models as a second step. Change in R squared between the steps was used to assess differences in the overall model after adding the interaction term with corresponding significance tests.

#### Cortico-limbic signature of genetic and environmental risk

In our final analyses, we leveraged a multivariate data-driven approach to assess how genetic and environmental risk covaries with cortico-limbic global and regional connectivity. Partial least squares (PLS) was used to model how genetic and environmental risk covaries with cortico-limbic connectivity. PLS is a data-reduction technique well suited to capturing covariance and explaining complex relationships between a large set of noisy or multicollinear variables. It models the relationship between predictor and outcome variables by simultaneously projecting them to a new space to obtain a set of orthogonal latent variables that represent linear combinations of the original variables.^92^ We conducted two separate PLS analyses to model the covariance between adversity/PRS and (1) cortico-limbic global connectivity; and (2) cortico-limbic regional connectivity (regional-PLS). In the first PLS, we modelled the four mental health PRSs and cumulative adversity as joint predictor variables (joint model), and connectivity between the limbic network and each of the 12 cortical networks as the outcome. We then repeated this using only the PRS scores (PRS-only model) and then only cumulative adversity (adversity-only model) as the predictor variables to assess unique associations between different dimensions of risk and cortico-limbic network connectivity. In our second PLS, we modelled the joint, PRS-only, and adversity-only predictors with connectivity between specific limbic regions (n=19) and each of the cortical networks as the outcome to assess possible regional heterogeneity within the limbic network. The *p*corr significance values for the PLS components were obtained by permuting the data 10,000 times and comparing the observed coefficients relative to their null distributions. Variable importance projection (VIP) was used to assess the relative importance of each variable in the model, with scores above 1 considered most influential in terms of their explanatory power. The stability of item loadings was assessed by averaging the mean squared error of prediction, R^2^, and Q^2^ across 10 cross-validation runs (nrepeat=10, folds=10), with a significance threshold of 0.01 for improvement in component error rate. To assess the stability of the model loadings, we employed a non-parametric bootstrapping approach with 1,000 resampling iterations to obtain standard error, t-values, confidence intervals and p-values for each item. Each bootstrap sample was created by randomly resampling with replacement from the original dataset ensuring that the matrix structure for both predictor and response variables was preserved. Variables were scaled to unit variance prior to analysis. We regressed the latent variables obtained from each PLS model to obtain the latent component slope, with age, sex, and the 6 PCs added as covariates. Mental health phenotype scores were added as a second set of covariates. In the PRS-only model, adversity was added as a final covariate. In the adversity-only model, PRS scores were added as final covariates. For any significant regression results, we tested for possible mediating effects of the component response scores (i.e., latent scores based on cortico-limbic connectivity measures) on the association between the component predictor scores (i.e., latent scores based on PRS/adversity measures) and mental health. One thousand bootstrap samples were used to estimate the 95% confidence intervals for indirect effects using the bias-corrected percentile method proposed by Biesanz and colleagues (2010). Analyses were performed in R using the *MixOmics* ^94^, *caret* ^95^ and *mdatools* ^96^.

## Acknowledgements

This study was funded by the UK Medical Research Council, Grant MC-A0606-5. DA is supported by the Gnodde Goldman Sachs endowed Professorship in Neuroinformatics, The James S. McDonnell Foundation Opportunity Awards, and by the Templeton World Charity Foundation, Inc. (funder DOI 501100011730) under grant TWCF-2002-30510. All research at the Department of Psychiatry at the University of Cambridge is supported by the National Institute for Health and Care Research Cambridge Biomedical Research Centre (NIHR203312) and the NIHR Applied Research Collaboration East England. The funders played no role in study design, data collection, analysis and interpretation of data, or the writing of this manuscript.

Data used in the preparation of this article were obtained from the Adolescent Brain Cognitive Development (ABCD) Study (https://abcdstudy.org), held in the NIMH Data Archive (NDA). ABCD consortium investigators designed and implemented the study and/or provided data but did not participate in the analysis or writing of this report. This manuscript reflects the views of the authors and may not reflect the opinions or views of the NIH or ABCD consortium investigators. The ABCD Study is supported by the National Institutes of Health and additional federal partners under award numbers U01DA041048, U01DA050989, U01DA051016, U01DA041022, U01DA051018, U01DA051037, U01DA050987, U01DA041174, U01DA041106, U01DA041117, U01DA041028, U01DA041134, U01DA050988, U01DA051039, U01DA041156, U01DA041025, U01DA041120, U01DA051038, U01DA041148, U01DA041093, U01DA041089, U24DA041123, U24DA041147. A full list of supporters is available at https://abcdstudy.org/federal-partners.html.

## Notes

### Competing Interest Statement

The authors have declared no competing interest.

