## Supplemental materials for "A common neural signature between genetic and environmental risk"

### 1. METHODS SUPPLEMENT

**Table S1.** Demographics for all ABCD participants at baseline

| | |
|---|---|
| *N* | 11,876 |
| Age mean *(SD)* | 9.9 *(0.6)* |
| % Female | 47.8% |
| *Race/Ethnicity* | |
| % White | 52.2% |
| % African American | 15.1% |
| % Hispanic | 3.9% |
| % Asian | 2.2% |
| % Other/Multi-racial | 25.9% |
| *Household characteristics* | |
| % Married caregivers | 65.5% |
| % College-level education | 59.4% |
| *Household income* | |
| < $25k | 17.2% |
| $25k - $49.99k | 22.4% |
| $50k -$74.99k | 14.5% |
| $75k -$99.99k | 30.5% |
| $100k + | 15.4% |

*Notes*. Age reported in years. College education reported if one or more caregivers has a college-level degree.

**Table S2**. Adversity questions

| Description | ABCD variable name | Timescale | % missing |
|---|---|---|---|
| parent alcohol problem | famhx_ss_momdad_alc_p | Lifetime (age 0-9) | 3.77 |
| parent drug use problem | famhx_ss_momdad_dg_p | Lifetime (age 0-9) | 5.45 |
| parent trouble with job/fights/police | famhx_ss_momdad_trb_p | Lifetime (age 0-9) | 3.05 |
| community shooting or stabbing | ksads_ptsd_raw_760_p | Lifetime (age 0-9) | 2.41 |
| shot, stabbed, beaten by non-family | ksads_ptsd_raw_761_p | Lifetime (age 0-9) | 2.41 |
| shot, stabbed, beaten by caregiver | ksads_ptsd_raw_762_p | Lifetime (age 0-9) | 2.41 |
| severely beaten by caregiver | ksads_ptsd_raw_763_p | Lifetime (age 0-9) | 2.41 |
| death threat by non-family | ksads_ptsd_raw_764_p | Lifetime (age 0-9) | 2.41 |
| death threat by family | ksads_ptsd_raw_765_p | Lifetime (age 0-9) | 2.41 |
| interparental violence | ksads_ptsd_raw_766_p | Lifetime (age 0-9) | 2.41 |
| sexual abuse by caregiver | ksads_ptsd_raw_767_p | Lifetime (age 0-9) | 2.41 |
| sexual abuse by non-family | ksads_ptsd_raw_768_p | Lifetime (age 0-9) | 2.41 |
| sexual abuse by peer | ksads_ptsd_raw_769_p | Lifetime (age 0-9) | 2.41 |
| unsafe community | nsc_p_ss_mean_3_items | Lifetime (age 0-9) | 0.07 |
| poor parental supervision | pmq_y_ss_mean | Lifetime (age 0-9) | 0.18 |
| low caregiver acceptance | crpbi_y_ss_parent | Lifetime (age 0-9) | 0.29 |
| financial difficulties: food | demo_fam_exp1_v2 | Past 12 months | 0.64 |
| financial difficulties: phone service | demo_fam_exp2_v2 | Past 12 months | 0.39 |
| financial difficulties: rent payment | demo_fam_exp3_v2 | Past 12 months | 0.51 |
| financial difficulties: eviction | demo_fam_exp4_v2 | Past 12 months | 0.28 |
| financial difficulties: gas and electric | demo_fam_exp5_v2 | Past 12 months | 0.37 |
| financial difficulties: medical care | demo_fam_exp6_v2 | Past 12 months | 0.34 |
| financial difficulties: dentist | demo_fam_exp7_v2 | Past 12 months | 0.41 |
| sudden death of loved one | ksads_ptsd_raw_770_p | Lifetime (age 0-9) | 2.41 |

*Notes*. Adversity items taken at baseline assessment (T1). Parental report used. Missingness represents the percentage of participants with missing data on a given question.

#### 1.2 Early life adversity

Missing adversity data was coded as "0" because sensitivity analyses revealed that either coding it as 1 or imputing it resulted in an overestimation of adversity in the sample relative to population prevalence estimates (Finkelhor et al., 2005; McLaughlin et al., 2012; Struck et al., 2020). In the first sensitivity analysis, we coded missing data as 1 instead of 0. This resulted in an additional 524 participants with adversity exposures, and an ELA group representing 27% of the total sample. This was much higher than population prevalence estimates would suggest, meaning this approach was heavily biased unlikely representative of real-world data.

Next, we used a multiple imputation package for mixed-type data, missForest in R (Stekhoven et al., 2012) to impute the missing values. Imputing missing values resulted in an additional 512 participants with adversity exposures (1251 in; 26.5% of the total sample), again much higher than population prevalence estimates (Finkelhor et al., 2005; McLaughlin et al., 2012; Struck et al., 2020). Imputation algorithms are heavily biased towards rare cases with binary data (e.g., exposures to ELA). This means adversity is likely over-estimated, explaining why using this method resulted in an unusually high number of children classified as having experienced adversity relative to population prevalence estimates (Finkelhor et al., 2005; McLaughlin et al., 2012; Struck et al., 2020). There were other reasons for not using imputation. First, it would have increased the standard error, which would have been problematic for our subsequent analyses. Second, the data were not missing at random. We tested for associations between missingness and several key cognitive and demographic variables. Missingness was associated interview ethnicity ($p$<.001); parental education ($p$<.001); parental income ($p$<.001) and 3 out of 5 measures of cognition that were tested ($p$s= .001-.05). The missing at random assumption was therefore not plausible, meaning that imputation would be heavily biased. Although imputation is beneficial in some cases, it must be balanced against the possible risks of inducing bias and overfitting, particularly in the case of non-normally distributed binary data (Sterne et al., 2009). While some procedures can handle non-normally distributed data better than others (Van Buuren et al., 1999), it is an ongoing area of development (Horton et al., 2007; Bernaards et al, 2007) that currently has no well-defined solution (Lee & Carlin, 2016; Sullivan et al., 2017). For these reasons, we decided imputation was not appropriate for our data.

### 2. RESULTS SUPPLEMENT

**Figure S1.** Results for PRS ADHD

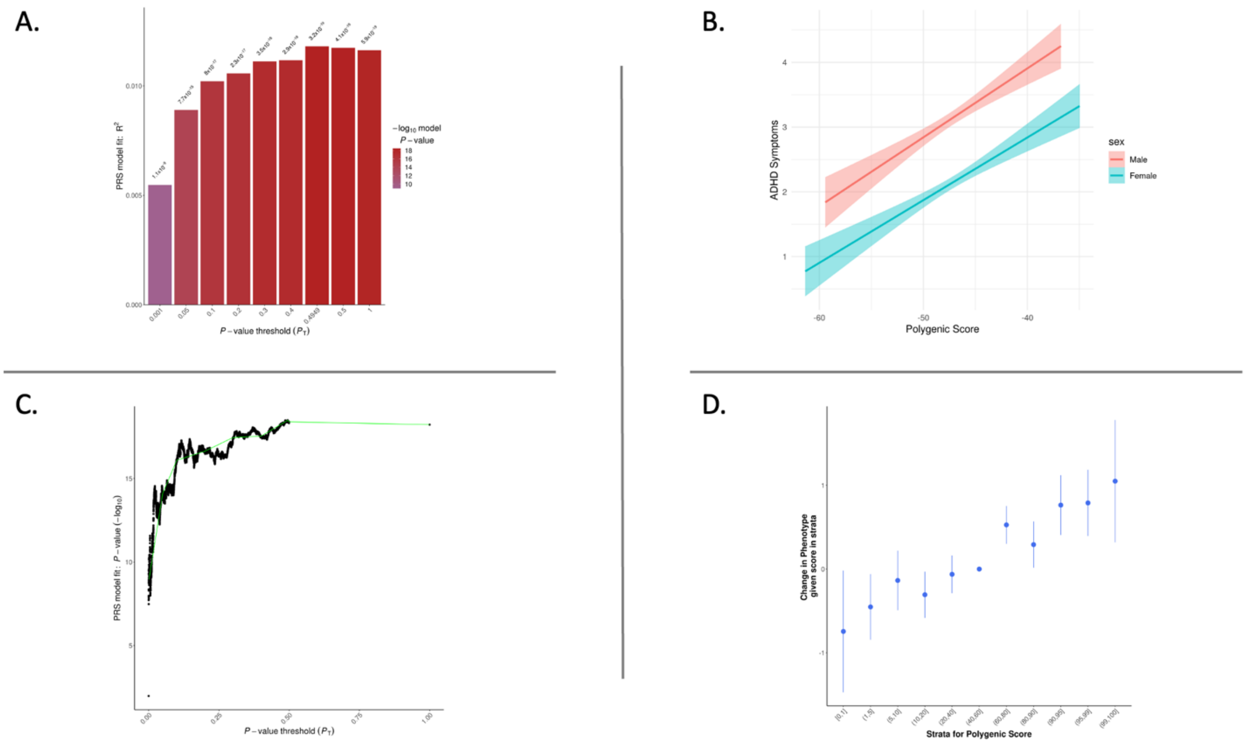


*Notes.* (A) Bar plot displaying the model fit of the PRS at different p-value thresholds; (B) PRS best fit results; (C) High resolution plot showing model fit at all p-value thresholds. The green line connects points showing the model fit at broad p-value thresholds used in the corresponding bar plot; (D) Strata plot providing an illustration of the effect of increased PRS on predicted phenotype using an uneven distribution of deciles.

**Figure S2.** Results for PRS Anxiety

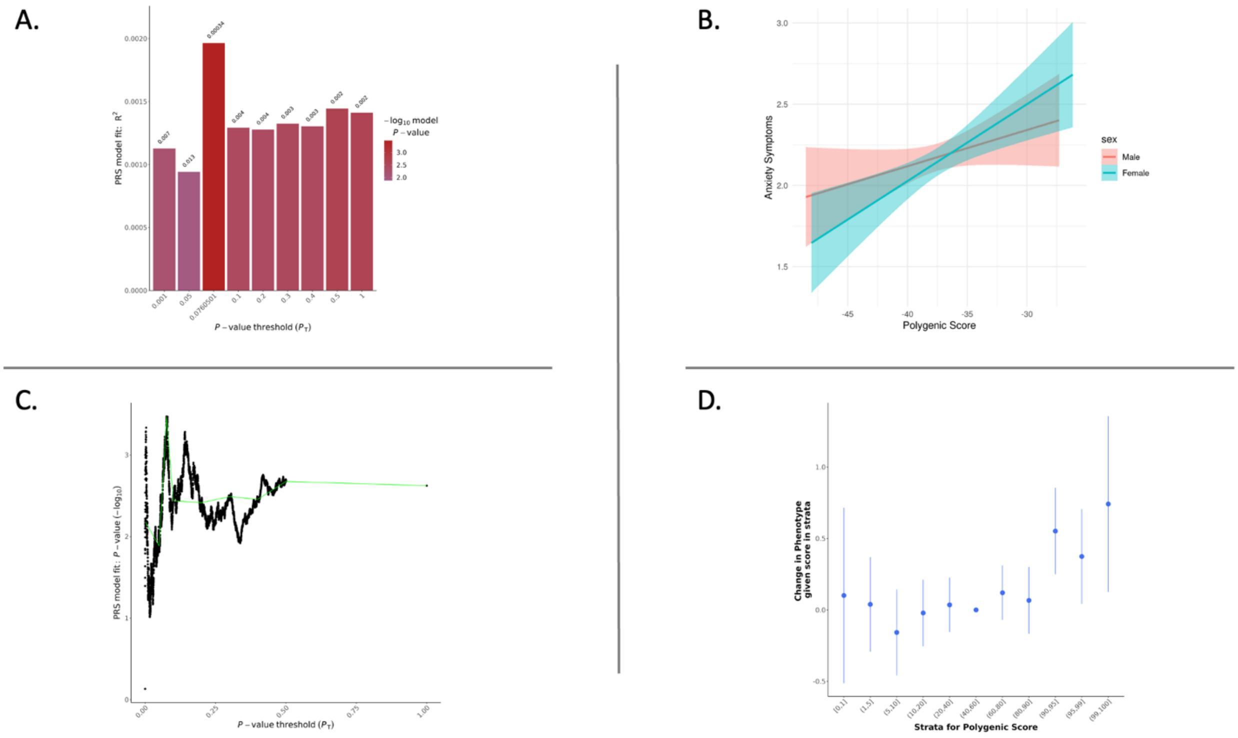


*Notes.* (A) Bar plot displaying the model fit of the PRS at different p-value thresholds; (B) PRS best fit results; (C) High resolution plot showing model fit at all p-value thresholds. The green line connects points showing the model fit at broad p-value thresholds used in the corresponding bar plot; (D) Strata plot providing an illustration of the effect of increased PRS on predicted phenotype using an uneven distribution of deciles.

**Figure S3.** Results for PRS Depression

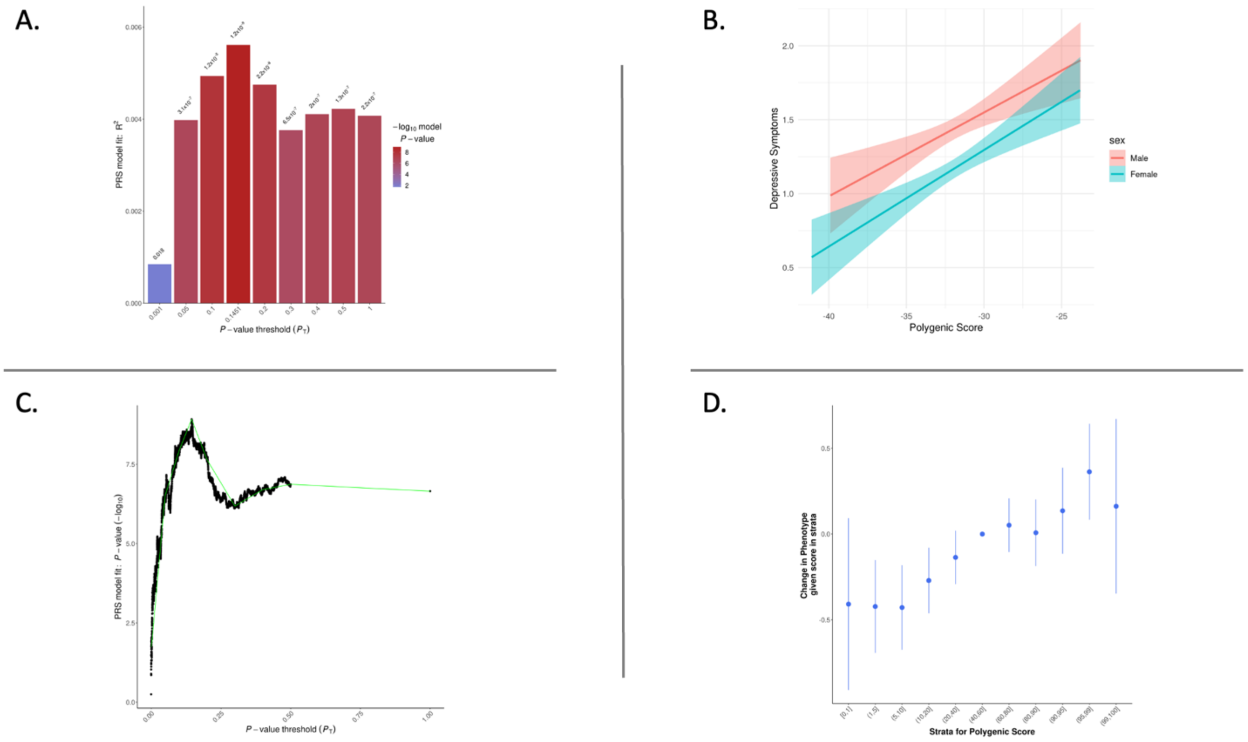


*Notes.* (A) Bar plot displaying the model fit of the PRS at different p-value thresholds; (B) PRS best fit results; (C) High resolution plot showing model fit at all p-value thresholds. The green line connects points showing the model fit at broad p-value thresholds used in the corresponding bar plot; (D) Strata plot providing an illustration of the effect of increased PRS on predicted phenotype using an uneven distribution of deciles.

**Figure S4.** Results for PRS Psychosis

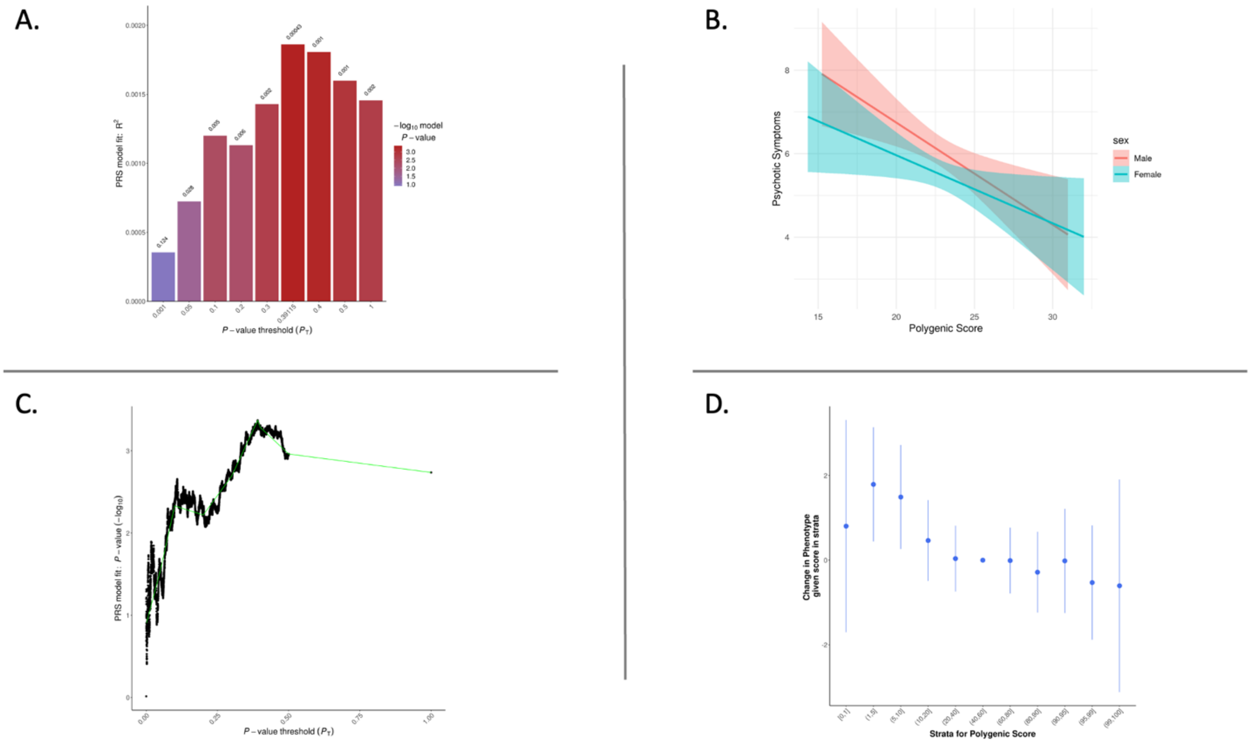


*Notes.* (A) Bar plot displaying the model fit of the PRS at different p-value thresholds; (B) PRS best fit results; (C) High resolution plot showing model fit at all p-value thresholds. The green line connects points showing the model fit at broad p-value thresholds used in the corresponding bar plot; (D) Strata plot providing an illustration of the effect of increased PRS on predicted phenotype using an uneven distribution of deciles.

**Figure S5.** Distribution of the PRS scores

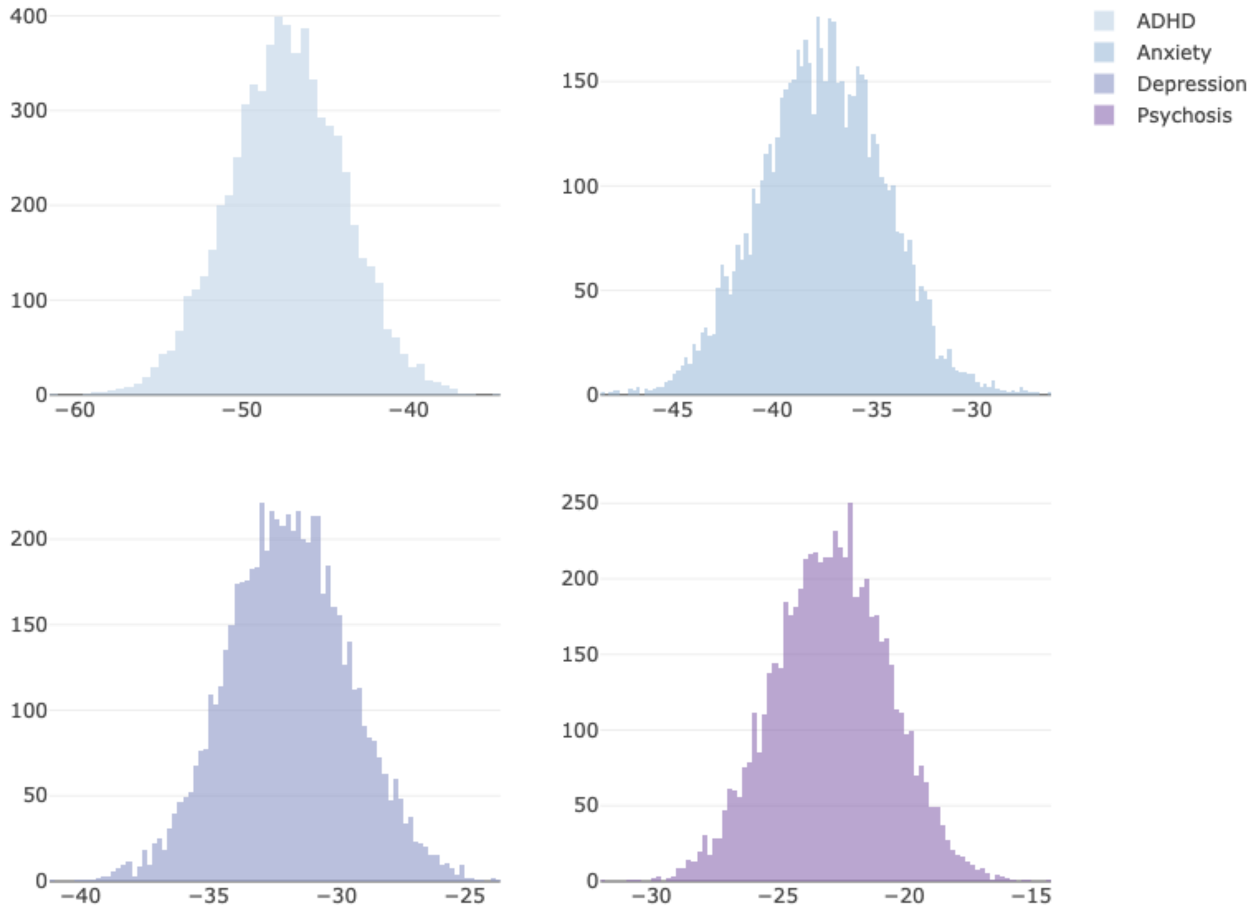


*Notes*. X-axis: unstandardised PRS; Y-axis: participant density.

**Figure S6.** PRS variance explained

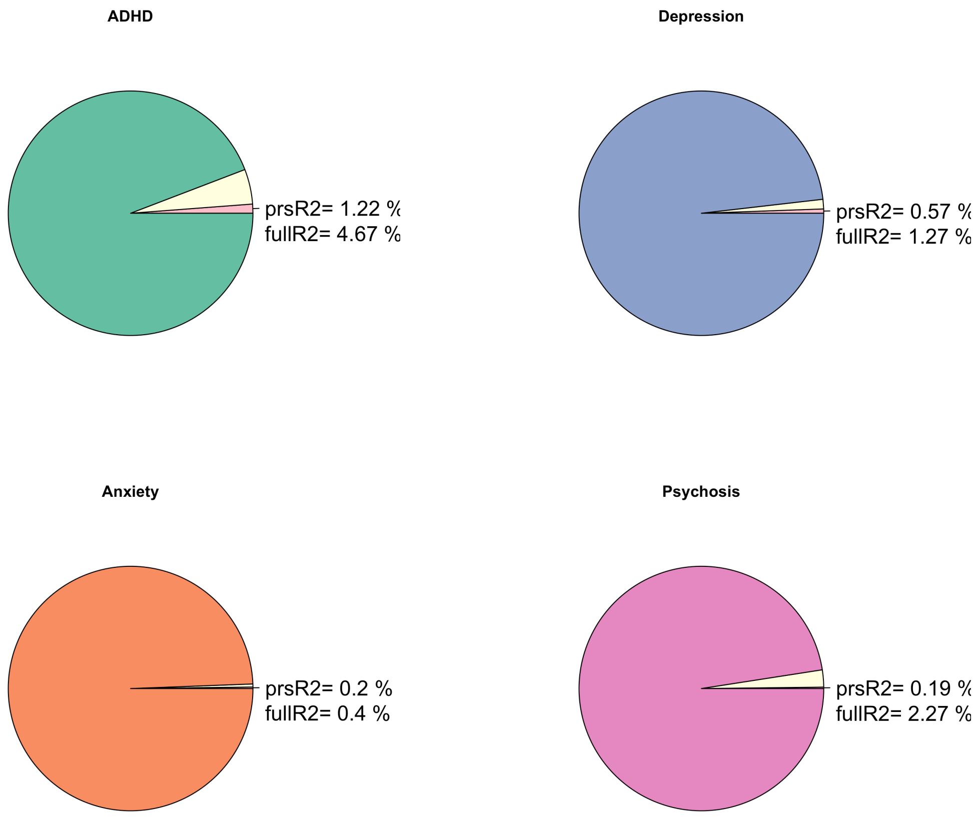


*Notes*. prsR2= Variance explained by the PRS controlling for age, sex and 6 population components (PCs). fullR2= Variance explained by the full model (including covariates age, sex, and 6 PCs).

**Table S3.** PRS best fit results

| Phenotype | Base GWAS | Thresh | PRS R2 | Full R2 | Null R2 | Coef | Std Err | P | SNPs |
|---|---|---|---|---|---|---|---|---|---|
| *Full sample* | | | | | | | | | |
| ADHD | Demontis 2023 | 0.495 | 0.012 | 0.047 | 0.035 | 0.100 | 0.011 | 0.0000 | 55120 |
| Anxiety | Meier 2019 | 0.076 | 0.002 | 0.004 | 0.002 | 0.037 | 0.010 | 0.0003 | 9026 |
| Depression | Wray 2018 | 0.145 | 0.006 | 0.013 | 0.007 | 0.072 | 0.012 | 0.0000 | 18925 |
| Psychosis | Richards 2022 | 0.391 | 0.002 | 0.023 | 0.021 | -0.197 | 0.056 | 0.0004 | 49762 |
| *European descent* | | | | | | | | | |
| ADHD | Demontis 2023 | 1.000 | 0.014 | 0.042 | 0.029 | 0.102 | 0.012 | 0.0000 | 79009 |
| Anxiety | Meier 2019 | 0.079 | 0.003 | 0.005 | 0.003 | 0.042 | 0.011 | 0.0002 | 9240 |
| Depression | Wray 2018 | 0.093 | 0.007 | 0.014 | 0.007 | 0.086 | 0.014 | 0.0000 | 13169 |
| Psychosis | Richards 2022 | 0.391 | 0.001 | 0.014 | 0.013 | -0.158 | 0.059 | 0.0074 | 48515 |
| *Non-European descent* | | | | | | | | | |
| ADHD | Demontis 2023 | 0.000 | 0.011 | 0.071 | 0.060 | 0.775 | 0.211 | 0.0003 | 183 |
| Anxiety | Meier 2019 | 0.000 | 0.007 | 0.023 | 0.016 | 0.427 | 0.148 | 0.0040 | 97 |
| Depression | Wray 2018 | 0.000 | 0.004 | 0.016 | 0.011 | -4.074 | 1.786 | 0.0227 | 1 |
| Psychosis | Richards 2022 | 0.109 | 0.008 | 0.026 | 0.018 | -0.624 | 0.206 | 0.0025 | 19598 |

*Notes*. Thresh= best p-value threshold. PRS R2= Variance explained by the PRS. Full R2= Variance explained by the full model (including covariates age, sex, and 6 PCs). Null R2= Variance explained by the covariates. Coef, Std Err and P= Regression coefficient, standard error, and significance for the model. SNPs= Number of SNPs included in the model.

**Table S4.** Hierarchical linear regression for variance explained in ADHD symptoms by ADHD PRS

| **Model** | | **Coef** | **Std Err** | **Std Coef** | **t** | **p** |
|---|---|---|---|---|---|---|
| $H_0$ | *(Intercept)* | 5.05 | 0.59 | | 8.58 | 0.000 |
| | age | -0.02 | 0.01 | -0.04 | -3.29 | 0.001 |
| | sex | -0.99 | 0.07 | | -13.53 | 0.000 |
| | PC1 | -12.57 | 2.95 | -0.05 | -4.26 | 0.000 |
| | PC2 | 9.95 | 2.95 | 0.04 | 3.37 | 0.001 |
| | PC3 | -3.60 | 2.95 | -0.02 | -1.22 | 0.222 |
| | PC4 | -8.73 | 2.95 | -0.04 | -2.96 | 0.003 |
| | PC5 | -5.20 | 2.94 | -0.02 | -1.77 | 0.078 |
| | PC6 | -0.14 | 2.95 | 0.00 | -0.05 | 0.962 |
| $H_1$ | *(Intercept)* | 9.79 | 0.79 | | 12.43 | 0.000 |
| | prsADHD | 0.10 | 0.01 | 0.12 | 8.99 | 0.000 |
| | age | -0.02 | 0.01 | -0.04 | -3.34 | 0.001 |
| | sex | -1.00 | 0.07 | | -13.76 | 0.000 |
| | PC1 | -4.39 | 3.07 | -0.02 | -1.43 | 0.153 |
| | PC2 | 9.81 | 2.93 | 0.04 | 3.35 | 0.001 |
| | PC3 | -6.00 | 2.94 | -0.03 | -2.04 | 0.042 |
| | PC4 | -7.37 | 2.93 | -0.03 | -2.51 | 0.012 |
| | PC5 | -7.36 | 2.94 | -0.03 | -2.51 | 0.012 |
| | PC6 | -0.63 | 2.93 | 0.00 | -0.22 | 0.830 |

| **Model** | **$R^2$** | **Adjusted $R^2$** | **$R^2$ Change** | **df1** | **df2** | **p** |
|---|---|---|---|---|---|---|
| $H_0$ | 0.035 | 0.034 | 0.035 | 8 | 6526 | 0.000 |
| $H_1$ | 0.047 | 0.045 | 0.012 | 1 | 6525 | 0.000 |

*Notes*. Null model includes age, sex, PC1, PC2, PC3, PC4, PC5, PC6.

**Table S5.** Hierarchical linear regression for variance explained in anxiety symptoms by Anxiety PRS

| Model | | Coef | Std Err | Std Coef | t | p |
|---|---|---|---|---|---|---|
| $H_0$ | *(Intercept)* | 2.76 | 0.50 | | 5.58 | 0.000 |
| | age | -0.01 | 0.00 | -0.02 | -1.19 | 0.234 |
| | sex | -0.03 | 0.06 | | -0.41 | 0.680 |
| | PC1 | -1.67 | 2.48 | -0.01 | -0.67 | 0.501 |
| | PC2 | 1.58 | 2.48 | 0.01 | 0.64 | 0.523 |
| | PC3 | -5.55 | 2.48 | -0.03 | -2.24 | 0.025 |
| | PC4 | -0.18 | 2.48 | 0.00 | -0.07 | 0.943 |
| | PC5 | -4.27 | 2.48 | -0.02 | -1.73 | 0.084 |
| | PC6 | -4.02 | 2.48 | -0.02 | -1.62 | 0.105 |
| $H_1$ | *(Intercept)* | 4.17 | 0.63 | | 6.62 | 0.000 |
| | prsAnxiety | 0.04 | 0.01 | 0.05 | 3.60 | 0.000 |
| | sex | -0.03 | 0.06 | | -0.45 | 0.650 |
| | age | -0.01 | 0.00 | -0.02 | -1.24 | 0.217 |
| | PC1 | 0.62 | 2.56 | 0.00 | 0.24 | 0.809 |
| | PC2 | 2.78 | 2.50 | 0.01 | 1.11 | 0.266 |
| | PC3 | -5.91 | 2.48 | -0.03 | -2.39 | 0.017 |
| | PC4 | -1.06 | 2.49 | -0.01 | -0.42 | 0.671 |
| | PC5 | -4.17 | 2.47 | -0.02 | -1.68 | 0.092 |
| | PC6 | -3.91 | 2.48 | -0.02 | -1.58 | 0.114 |

| Model | $R^2$ | Adjusted $R^2$ | $R^2$ Change | df1 | df2 | p |
|---|---|---|---|---|---|---|
| $H_0$ | 0.002 | 0.001 | 0.002 | 8 | 6526 | 0.103 |
| $H_1$ | 0.004 | 0.003 | 0.002 | 1 | 6525 | 0.000 |

*Notes*. Null model includes age, sex, PC1, PC2, PC3, PC4, PC5, PC6.

**Table S6.** Hierarchical linear regression for variance explained in depression symptoms by Depression PRS

| Model | | Coef | Std Err | Std Coef | t | p |
|---|---|---|---|---|---|---|
| $H_0$ | *(Intercept)* | 0.91 | 0.41 | | 2.25 | 0.025 |
| | age | 0.00 | 0.00 | 0.02 | 1.32 | 0.186 |
| | sex | -0.27 | 0.05 | | -5.34 | 0.000 |
| | PC1 | -4.58 | 2.03 | -0.03 | -2.26 | 0.024 |
| | PC2 | 3.09 | 2.03 | 0.02 | 1.52 | 0.128 |
| | PC3 | -3.97 | 2.03 | -0.02 | -1.96 | 0.050 |
| | PC4 | -0.23 | 2.03 | 0.00 | -0.11 | 0.911 |
| | PC5 | -3.92 | 2.03 | -0.02 | -1.94 | 0.053 |
| | PC6 | -1.80 | 2.03 | -0.01 | -0.89 | 0.374 |
| $H_1$ | *(Intercept)* | 3.25 | 0.56 | | 5.83 | 0.000 |
| | prsDepression | 0.07 | 0.01 | 0.09 | 6.09 | 0.000 |
| | age | 0.00 | 0.00 | 0.02 | 1.20 | 0.230 |
| | sex | -0.27 | 0.05 | | -5.32 | 0.000 |
| | PC1 | 1.28 | 2.24 | 0.01 | 0.57 | 0.569 |
| | PC2 | 6.01 | 2.08 | 0.04 | 2.89 | 0.004 |
| | PC3 | -4.39 | 2.02 | -0.03 | -2.17 | 0.030 |
| | PC4 | -1.55 | 2.03 | -0.01 | -0.76 | 0.446 |
| | PC5 | -4.16 | 2.02 | -0.03 | -2.06 | 0.039 |
| | PC6 | -2.00 | 2.02 | -0.01 | -0.99 | 0.322 |

| Model | $R^2$ | Adjusted $R^2$ | $R^2$ Change | df1 | df2 | p |
|---|---|---|---|---|---|---|
| $H_0$ | 0.007 | 0.006 | 0.007 | 8 | 6526 | 0.000 |
| $H_1$ | 0.013 | 0.011 | 0.006 | 1 | 6525 | 0.000 |

*Notes*. Null model includes age, sex, PC1, PC2, PC3, PC4, PC5, PC6.

**Table S7.** Hierarchical linear regression for variance explained in psychotic symptoms by Psychosis PRS

| Model | | Coef | Std Err | Std Coef | t | p |
|---|---|---|---|---|---|---|
| $H_0$ | (Intercept) | 16.14 | 2.02 | | 7.98 | 0.000 |
| | age | -0.09 | 0.02 | -0.06 | -5.01 | 0.000 |
| | sex | -0.57 | 0.25 | | -2.27 | 0.023 |
| | PC1 | -80.18 | 10.15 | -0.10 | -7.90 | 0.000 |
| | PC2 | -42.00 | 10.14 | -0.05 | -4.14 | 0.000 |
| | PC3 | -47.60 | 10.14 | -0.06 | -4.70 | 0.000 |
| | PC4 | 19.33 | 10.13 | 0.02 | 1.91 | 0.057 |
| | PC5 | -2.14 | 10.13 | 0.00 | -0.21 | 0.833 |
| | PC6 | -3.63 | 10.14 | 0.00 | -0.36 | 0.720 |
| $H_1$ | (Intercept) | 20.53 | 2.38 | | 8.62 | 0.000 |
| | prsPsychosis | 0.20 | 0.06 | 0.04 | 3.49 | 0.000 |
| | age | -0.08 | 0.02 | -0.06 | -4.97 | 0.000 |
| | sex | -0.56 | 0.25 | | -2.24 | 0.025 |
| | PC1 | -77.46 | 10.17 | -0.09 | -7.61 | 0.000 |
| | PC2 | -44.62 | 10.16 | -0.05 | -4.39 | 0.000 |
| | PC3 | -46.65 | 10.14 | -0.06 | -4.60 | 0.000 |
| | PC4 | 22.44 | 10.16 | 0.03 | 2.21 | 0.027 |
| | PC5 | -5.90 | 10.18 | -0.01 | -0.58 | 0.562 |
| | PC6 | -4.98 | 10.13 | -0.01 | -0.49 | 0.623 |
| **Model** | **$R^2$** | **Adjusted $R^2$** | **$R^2$ Change** | **df1** | **df2** | **p** |
| $H_0$ | 0.021 | 0.02 | 0.021 | 8 | 6526 | 0.000 |
| $H_1$ | 0.023 | 0.021 | 0.002 | 1 | 6525 | 0.000 |

*Notes*. Null model includes age, sex, PC1, PC2, PC3, PC4, PC5, PC6.

**Table S8**. Canonical loadings for mental health PRS scores and phenotypes

| | | | Pearson's r | Lower 95% CI | Upper 95% CI | p | |
|---|---|---|---|---|---|---|---|
| *Canonical Variate 1* | | | | | | | |
| CV1prs | - | prsADHD | -0.87 | -0.88 | -0.87 | 0.000 | *** |
| CV1prs | - | prsAnxiety | -0.35 | -0.37 | -0.33 | 0.000 | *** |
| CV1prs | - | prsDepression | -0.69 | -0.71 | -0.68 | 0.000 | *** |
| CV1prs | - | prsPsychosis | -0.35 | -0.37 | -0.33 | 0.000 | *** |
| CV1prs | - | phenoADHD | -0.11 | -0.14 | -0.09 | 0.000 | *** |
| CV1prs | - | phenoAnxiety | -0.05 | -0.07 | -0.02 | 0.000 | *** |
| CV1prs | - | phenoDepression | -0.07 | -0.09 | -0.05 | 0.000 | *** |
| CV1prs | - | phenoPsychosis | -0.12 | -0.14 | -0.09 | 0.000 | *** |
| CV1pheno | - | prsADHD | -0.13 | -0.16 | -0.11 | 0.000 | *** |
| CV1pheno | - | prsAnxiety | -0.05 | -0.08 | -0.03 | 0.000 | *** |
| CV1pheno | - | prsDepression | -0.11 | -0.13 | -0.08 | 0.000 | *** |
| CV1pheno | - | prsPsychosis | -0.05 | -0.08 | -0.03 | 0.000 | *** |
| CV1pheno | - | phenoADHD | -0.74 | -0.75 | -0.73 | 0.000 | *** |
| CV1pheno | - | phenoAnxiety | -0.31 | -0.33 | -0.29 | 0.000 | *** |
| CV1pheno | - | phenoDepression | -0.46 | -0.48 | -0.44 | 0.000 | *** |
| CV1pheno | - | phenoPsychosis | -0.77 | -0.78 | -0.76 | 0.000 | *** |
| *Canonical Variate 2* | | | | | | | |
| CV2prs | - | prsADHD | 0.40 | 0.38 | 0.42 | 0.000 | *** |
| CV2prs | - | prsAnxiety | -0.51 | -0.52 | -0.49 | 0.000 | *** |
| CV2prs | - | prsDepression | -0.67 | -0.68 | -0.66 | 0.000 | *** |
| CV2prs | - | prsPsychosis | -0.12 | -0.15 | -0.10 | 0.000 | *** |
| CV2prs | - | phenoADHD | 0.04 | 0.02 | 0.06 | 0.001 | ** |
| CV2prs | - | phenoAnxiety | -0.04 | -0.06 | -0.01 | 0.002 | ** |
| CV2prs | - | phenoDepression | -0.04 | -0.06 | -0.01 | 0.004 | ** |
| CV2prs | - | phenoPsychosis | -0.04 | -0.06 | -0.01 | 0.004 | ** |
| CV2pheno | - | prsADHD | 0.04 | 0.01 | 0.06 | 0.003 | ** |
| CV2pheno | - | prsAnxiety | -0.05 | -0.07 | -0.02 | 0.000 | *** |
| CV2pheno | - | prsDepression | -0.06 | -0.09 | -0.04 | 0.000 | *** |
| CV2pheno | - | prsPsychosis | -0.01 | -0.04 | 0.01 | 0.364 | |
| CV2pheno | - | phenoADHD | 0.42 | 0.40 | 0.44 | 0.000 | *** |
| CV2pheno | - | phenoAnxiety | -0.41 | -0.43 | -0.38 | 0.000 | *** |
| CV2pheno | - | phenoDepression | -0.38 | -0.40 | -0.36 | 0.000 | *** |
| CV2pheno | - | phenoPsychosis | -0.38 | -0.40 | -0.36 | 0.000 | *** |

*Notes*. CV1= First canonical variate. CV2=Second canonical variate. CV#prs= Canonical component for PRS scores. CV#pheno= Canonical component for phenotypes. * $p < .05$, ** $p < .01$, *** $p < .001$.

**Figure S7.** Canonical correlations for the first and second canonical variates

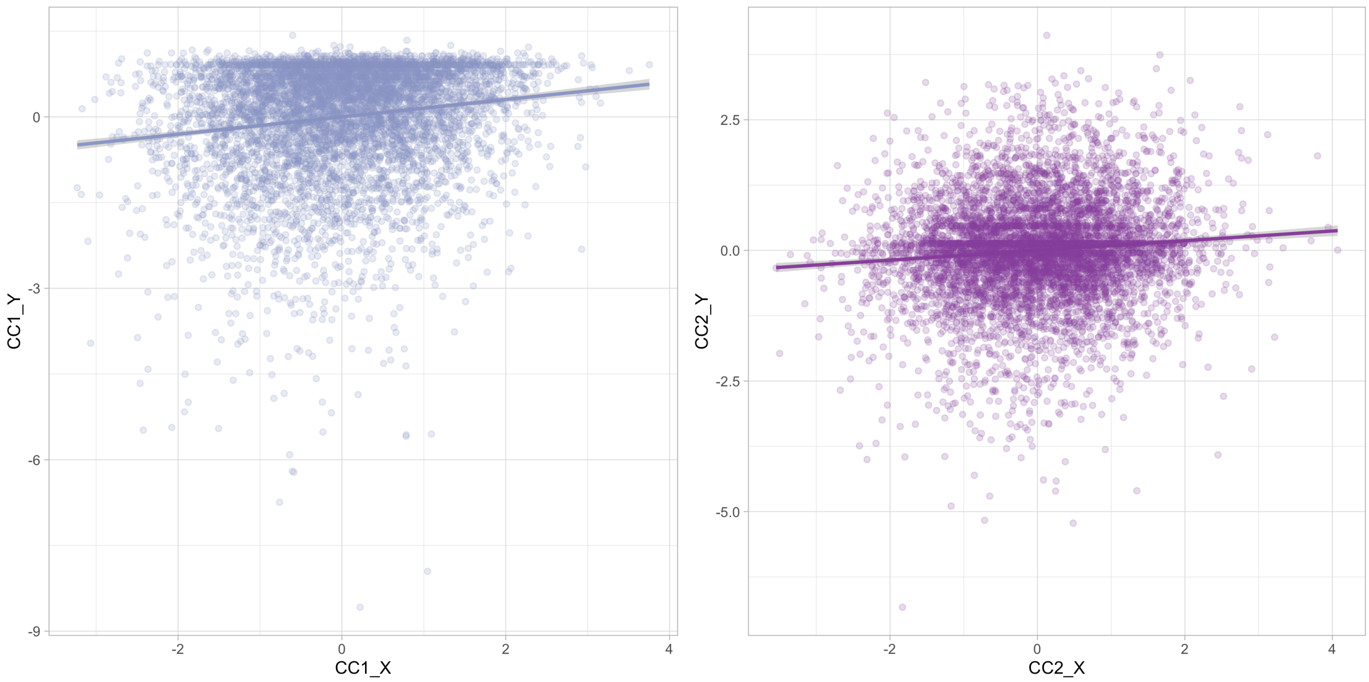


*Notes*. Correlation between predictor (X= PRS scores) and outcome (Y= phenotype scores) for each canonical variate.

**Table S9.** Hierarchical linear regression for variance explained in ADHD symptoms by the PRS component scores

| Model | | Coef | Std Err | Std Coef | t | p |
|---|---|---|---|---|---|---|
| $H_0$ | *(Intercept)* | 5.05 | 0.59 | | 8.58 | 0.000 |
| | age | -0.02 | 0.01 | -0.04 | -3.29 | 0.001 |
| | sex | -0.99 | 0.07 | | -13.54 | 0.000 |
| | PC1 | -12.63 | 2.95 | -0.05 | -4.28 | 0.000 |
| | PC2 | 9.91 | 2.95 | 0.04 | 3.36 | 0.001 |
| | PC3 | -3.63 | 2.95 | -0.02 | -1.23 | 0.218 |
| | PC4 | -8.71 | 2.95 | -0.04 | -2.96 | 0.003 |
| | PC5 | -5.21 | 2.95 | -0.02 | -1.77 | 0.077 |
| | PC6 | -0.14 | 2.95 | 0.00 | -0.05 | 0.962 |
| $H_1$ | *(Intercept)* | 5.08 | 0.59 | | 8.69 | 0.000 |
| | prsCVgeneral | 0.35 | 0.04 | 0.12 | 8.66 | 0.000 |
| | prsCVneurodev | 0.09 | 0.04 | 0.03 | 2.44 | 0.015 |
| | age | -0.02 | 0.01 | -0.04 | -3.35 | 0.001 |
| | sex | -1.00 | 0.07 | | -13.73 | 0.000 |
| | PC1 | -2.51 | 3.30 | -0.01 | -0.76 | 0.447 |
| | PC2 | 10.44 | 3.02 | 0.04 | 3.45 | 0.001 |
| | PC3 | -5.83 | 2.94 | -0.02 | -1.99 | 0.047 |
| | PC4 | -7.28 | 2.96 | -0.03 | -2.46 | 0.014 |
| | PC5 | -7.93 | 2.94 | -0.03 | -2.70 | 0.007 |
| | PC6 | -0.95 | 2.93 | 0.00 | -0.33 | 0.745 |
| **Model** | **$R^2$** | **Adjusted $R^2$** | **$R^2$ Change** | **df1** | **df2** | **p** |
| $H_0$ | 0.035 | 0.034 | 0.035 | 8 | 6524 | 0.000 |
| $H_1$ | 0.048 | 0.046 | 0.013 | 2 | 6522 | 0.000 |

*Notes*. prsCVgeneral = PRS component scores from the first canonical variate representing general psychopathology. prsCVneurodev = PRS component scores from the second canonical variate representing neurodevelopmental-specific variance. Null model includes age, sex, PC1, PC2, PC3, PC4, PC5, PC6.

**Table S10.** Hierarchical linear regression for variance explained in Anxiety symptoms by the PRS component scores

| Model | | Coef | Std Err | Std Coef | t | p |
|---|---|---|---|---|---|---|
| $H_0$ | *(Intercept)* | 2.76 | 0.50 | | 5.58 | 0.000 |
| | age | -0.01 | 0.00 | -0.02 | -1.19 | 0.233 |
| | sex | -0.03 | 0.06 | | -0.42 | 0.675 |
| | PC1 | -1.74 | 2.48 | -0.01 | -0.70 | 0.485 |
| | PC2 | 1.60 | 2.48 | 0.01 | 0.65 | 0.518 |
| | PC3 | -5.56 | 2.48 | -0.03 | -2.25 | 0.025 |
| | PC4 | -0.17 | 2.48 | 0.00 | -0.07 | 0.946 |
| | PC5 | -4.29 | 2.48 | -0.02 | -1.73 | 0.083 |
| | PC6 | -4.03 | 2.48 | -0.02 | -1.63 | 0.104 |
| $H_1$ | *(Intercept)* | 2.81 | 0.49 | | 5.70 | 0.000 |
| | prsCVgeneral | 0.16 | 0.03 | 0.07 | 4.70 | 0.000 |
| | prsCVneurodev | -0.12 | 0.03 | -0.05 | -3.54 | 0.000 |
| | age | -0.01 | 0.00 | -0.02 | -1.30 | 0.193 |
| | sex | -0.03 | 0.06 | | -0.42 | 0.673 |
| | PC1 | 5.32 | 2.79 | 0.03 | 1.91 | 0.056 |
| | PC2 | 4.58 | 2.55 | 0.02 | 1.79 | 0.073 |
| | PC3 | -6.27 | 2.48 | -0.03 | -2.53 | 0.011 |
| | PC4 | -1.42 | 2.50 | -0.01 | -0.57 | 0.570 |
| | PC5 | -4.83 | 2.48 | -0.02 | -1.95 | 0.052 |
| | PC6 | -4.28 | 2.47 | -0.02 | -1.73 | 0.084 |

| Model | $R^2$ | Adjusted $R^2$ | $R^2$ Change | df1 | df2 | p |
|---|---|---|---|---|---|---|
| $H_0$ | 0.045 | 0.002 | 0.002 | 8 | 6524 | 0.100 |
| $H_1$ | 0.083 | 0.007 | 0.005 | 2 | 6522 | 0.000 |

*Notes*. prsCVgeneral = PRS component scores from the first canonical variate representing general psychopathology. prsCVneurodev = PRS component scores from the second canonical variate representing neurodevelopmental-specific variance. Null model includes age, sex, PC1, PC2, PC3, PC4, PC5, PC6.

**Table S11.** Hierarchical linear regression for variance explained in Depression symptoms by the PRS component scores

| Model | | Coef | Std Err | Std Coef | t | p |
|---|---|---|---|---|---|---|
| $H_0$ | *(Intercept)* | 0.91 | 0.41 | | 2.25 | 0.024 |
| | age | 0.00 | 0.00 | 0.02 | 1.32 | 0.186 |
| | sex | -0.27 | 0.05 | | -5.36 | 0.000 |
| | PC1 | -4.67 | 2.03 | -0.03 | -2.30 | 0.021 |
| | PC2 | 3.09 | 2.03 | 0.02 | 1.53 | 0.127 |
| | PC3 | -3.99 | 2.03 | -0.02 | -1.97 | 0.049 |
| | PC4 | -0.21 | 2.03 | 0.00 | -0.10 | 0.917 |
| | PC5 | -3.95 | 2.03 | -0.02 | -1.95 | 0.051 |
| | PC6 | -1.82 | 2.03 | -0.01 | -0.90 | 0.371 |
| $H_1$ | *(Intercept)* | 0.96 | 0.40 | | 2.38 | 0.018 |
| | prsCVgeneral | 0.17 | 0.03 | 0.08 | 6.06 | 0.000 |
| | prsCVneurodev | -0.09 | 0.03 | -0.04 | -3.33 | 0.001 |
| | age | 0.00 | 0.00 | 0.02 | 1.21 | 0.227 |
| | sex | -0.27 | 0.05 | | -5.39 | 0.000 |
| | PC1 | 2.27 | 2.28 | 0.01 | 1.00 | 0.320 |
| | PC2 | 5.66 | 2.08 | 0.03 | 2.72 | 0.007 |
| | PC3 | -4.80 | 2.03 | -0.03 | -2.37 | 0.018 |
| | PC4 | -1.14 | 2.04 | -0.01 | -0.56 | 0.579 |
| | PC5 | -4.67 | 2.03 | -0.03 | -2.30 | 0.022 |
| | PC6 | -2.10 | 2.02 | -0.01 | -1.04 | 0.298 |

| Model | $R^2$ | Adjusted $R^2$ | $R^2$ Change | df1 | df2 | p |
|---|---|---|---|---|---|---|
| $H_0$ | 0.007 | 0.006 | 0.007 | 8 | 6524 | 0.000 |
| $H_1$ | 0.014 | 0.012 | 0.007 | 2 | 6522 | 0.000 |

*Notes*. prsCVgeneral = PRS component scores from the first canonical variate representing general psychopathology. prsCVneurodev = PRS component scores from the second canonical variate representing neurodevelopmental-specific variance. Null model includes age, sex, PC1, PC2, PC3, PC4, PC5, PC6.

**Table S12.** Hierarchical linear regression for variance explained in Psychosis symptoms by the PRS component scores

| Model | | Coef | Std Err | Std Coef | t | p |
|---|---|---|---|---|---|---|
| $H_0$ | *(Intercept)* | 16.13 | 2.02 | | 7.97 | 0.000 |
| | age | -0.09 | 0.02 | -0.06 | -5.00 | 0.000 |
| | sex | -0.57 | 0.25 | | -2.26 | 0.024 |
| | PC1 | -80.09 | 10.16 | -0.10 | -7.89 | 0.000 |
| | PC2 | -41.84 | 10.15 | -0.05 | -4.12 | 0.000 |
| | PC3 | -47.52 | 10.14 | -0.06 | -4.69 | 0.000 |
| | PC4 | 19.34 | 10.14 | 0.02 | 1.91 | 0.056 |
| | PC5 | -2.17 | 10.13 | 0.00 | -0.21 | 0.830 |
| | PC6 | -3.60 | 10.14 | 0.00 | -0.36 | 0.723 |
| $H_1$ | *(Intercept)* | 16.31 | 2.02 | | 8.09 | 0.000 |
| | prsCVgeneral | 1.00 | 0.14 | 0.10 | 7.17 | 0.000 |
| | prsCVneurodev | -0.11 | 0.13 | -0.01 | -0.85 | 0.397 |
| | age | -0.09 | 0.02 | -0.06 | -5.11 | 0.000 |
| | sex | -0.58 | 0.25 | | -2.32 | 0.021 |
| | PC1 | -45.36 | 11.38 | -0.06 | -3.98 | 0.000 |
| | PC2 | -33.77 | 10.42 | -0.04 | -3.24 | 0.001 |
| | PC3 | -53.09 | 10.14 | -0.06 | -5.24 | 0.000 |
| | PC4 | 18.83 | 10.22 | 0.02 | 1.84 | 0.066 |
| | PC5 | -8.24 | 10.15 | -0.01 | -0.81 | 0.417 |
| | PC6 | -5.63 | 10.11 | -0.01 | -0.56 | 0.578 |

| Model | $R^2$ | Adjusted $R^2$ | $R^2$ Change | df1 | df2 | p |
|---|---|---|---|---|---|---|
| $H_0$ | 0.021 | 0.020 | 0.021 | 8 | 6524 | 0.000 |
| $H_1$ | 0.028 | 0.027 | 0.008 | 2 | 6522 | 0.000 |

*Notes*. prsCVgeneral = PRS component scores from the first canonical variate representing general psychopathology. prsCVneurodev = PRS component scores from the second canonical variate representing neurodevelopmental-specific variance. Null model includes age, sex, PC1, PC2, PC3, PC4, PC5, PC6.

**Table S13.** Association between the general psychopathology PRS component and cumulative adversity

| | Coef | Std Err | Std Coef | t | Lower 95% CI | Upper 95% CI | p |
|---|---|---|---|---|---|---|---|
| (Intercept) | 1.670 | 0.357 | | 4.681 | 0.971 | 2.370 | 0.000 |
| CV1prs | 0.166 | 0.025 | 0.089 | 6.762 | 0.118 | 0.215 | 0.000 |
| CV1pheno | 0.447 | 0.023 | 0.239 | 19.800 | 0.403 | 0.492 | 0.000 |
| age | -0.002 | 0.003 | -0.010 | -0.812 | -0.008 | 0.003 | 0.417 |
| sex | 0.129 | 0.045 | | 2.888 | 0.041 | 0.216 | 0.004 |
| PC1 | -18.741 | 1.959 | -0.124 | -9.566 | -22.581 | -14.901 | 0.000 |
| PC2 | 10.671 | 1.791 | 0.070 | 5.958 | 7.160 | 14.182 | 0.000 |
| PC3 | -10.877 | 1.792 | -0.072 | -6.071 | -14.389 | -7.365 | 0.000 |
| PC4 | -0.521 | 1.773 | -0.003 | -0.294 | -3.996 | 2.954 | 0.769 |
| PC5 | -14.017 | 1.789 | -0.093 | -7.836 | -17.524 | -10.511 | 0.000 |
| PC6 | -4.093 | 1.781 | -0.027 | -2.298 | -7.584 | -0.601 | 0.022 |

*Notes*. prsCVgeneral = PRS component scores from the first canonical variate representing general psychopathology. phenoCVgeneral= Phenotype component scores from the first canonical variate representing general psychopathology.

**Table S14.** Association between the neurodevelopmental-specific PRS component and cumulative adversity

| | Coef | Std Err | Std Coef | t | Lower 95% CI | Upper 95% CI | p |
|---|---|---|---|---|---|---|---|
| (Intercept) | 2.213 | 0.368 | | 6.019 | 1.492 | 2.934 | 0.000 |
| CV2prs | 0.000 | 0.024 | 0.000 | -0.012 | -0.048 | 0.047 | 0.991 |
| CV2pheno | -0.175 | 0.023 | -0.093 | -7.574 | -0.220 | -0.130 | 0.000 |
| age | -0.006 | 0.003 | -0.025 | -2.071 | -0.012 | 0.000 | 0.038 |
| sex | -0.025 | 0.046 | | -0.545 | -0.115 | 0.065 | 0.586 |
| PC1 | -27.744 | 1.877 | -0.183 | -14.777 | -31.424 | -24.063 | 0.000 |
| PC2 | 10.035 | 1.891 | 0.066 | 5.307 | 6.328 | 13.741 | 0.000 |
| PC3 | -11.177 | 1.842 | -0.074 | -6.067 | -14.789 | -7.566 | 0.000 |
| PC4 | -1.625 | 1.854 | -0.011 | -0.876 | -5.260 | 2.011 | 0.381 |
| PC5 | -13.520 | 1.842 | -0.089 | -7.339 | -17.132 | -9.909 | 0.000 |
| PC6 | -3.654 | 1.839 | -0.024 | -1.987 | -7.258 | -0.049 | 0.047 |

*Notes*. prsCVneurodev= PRS component scores from the second canonical variate representing neurodevelopmental-specific variance. phenoCVneurodev= Phenotype component scores from the second canonical variate representing neurodevelopmental-specific variance.

**Table S15.** Association between PRS ADHD and cumulative adversity

| | Coef | Std Err | Std Coef | t | Lower 95% CI | Upper 95% CI | p |
|---|---|---|---|---|---|---|---|
| (Intercept) | 3.376 | 0.486 | | 6.947 | 2.423 | 4.328 | 0.000 |
| prsADHD | 0.042 | 0.007 | 0.077 | 6.123 | 0.028 | 0.055 | 0.000 |
| phenoADHD | 0.148 | 0.008 | 0.238 | 19.724 | 0.134 | 0.163 | 0.000 |
| age | -0.004 | 0.003 | -0.014 | -1.219 | -0.009 | 0.002 | 0.223 |
| sex | 0.158 | 0.045 | | 3.519 | 0.070 | 0.246 | 0.000 |
| PC1 | -22.549 | 1.868 | -0.149 | -12.074 | -26.210 | -18.888 | 0.000 |
| PC2 | 7.828 | 1.789 | 0.052 | 4.377 | 4.322 | 11.334 | 0.000 |
| PC3 | -12.082 | 1.793 | -0.080 | -6.737 | -15.598 | -8.567 | 0.000 |
| PC4 | 0.935 | 1.779 | 0.006 | 0.525 | -2.553 | 4.422 | 0.599 |
| PC5 | -13.612 | 1.791 | -0.090 | -7.601 | -17.123 | -10.102 | 0.000 |
| PC6 | -4.081 | 1.784 | -0.027 | -2.288 | -7.577 | -0.585 | 0.022 |

**Table S16.** Association between PRS Anxiety and cumulative adversity

| | Coef | Std Err | Std Coef | t | Lower 95% CI | Upper 95% CI | p |
|---|---|---|---|---|---|---|---|
| (Intercept) | 3.179 | 0.457 | | 6.956 | 2.283 | 4.074 | 0.000 |
| prsAnxiety | 0.039 | 0.007 | 0.065 | 5.272 | 0.025 | 0.054 | 0.000 |
| phenoAnxiety | 0.180 | 0.009 | 0.237 | 20.061 | 0.162 | 0.197 | 0.000 |
| age | -0.006 | 0.003 | -0.022 | -1.871 | -0.011 | 0.000 | 0.061 |
| sex | 0.016 | 0.044 | | 0.371 | -0.070 | 0.103 | 0.710 |
| PC1 | -25.045 | 1.844 | -0.165 | -13.584 | -28.659 | -21.431 | 0.000 |
| PC2 | 10.264 | 1.806 | 0.068 | 5.682 | 6.723 | 13.804 | 0.000 |
| PC3 | -11.066 | 1.790 | -0.073 | -6.181 | -14.575 | -7.556 | 0.000 |
| PC4 | -1.825 | 1.787 | -0.012 | -1.021 | -5.327 | 1.678 | 0.307 |
| PC5 | -12.511 | 1.787 | -0.083 | -7.000 | -16.015 | -9.007 | 0.000 |
| PC6 | -2.962 | 1.786 | -0.020 | -1.658 | -6.464 | 0.540 | 0.097 |

**Table S17.** Association between PRS Depression and cumulative adversity

| | Coef | Std Err | Std Coef | t | Lower 95% CI | Upper 95% CI | p |
|---|---|---|---|---|---|---|---|
| (Intercept) | 3.396 | 0.486 | | 6.980 | 2.442 | 4.349 | 0.000 |
| prsDepression | 0.046 | 0.010 | 0.059 | 4.413 | 0.025 | 0.066 | 0.000 |
| phenoDepression | 0.267 | 0.011 | 0.289 | 24.737 | 0.246 | 0.288 | 0.000 |
| age | -0.008 | 0.003 | -0.030 | -2.559 | -0.013 | -0.002 | 0.011 |
| sex | 0.089 | 0.044 | | 2.037 | 0.003 | 0.175 | 0.042 |
| PC1 | -22.779 | 1.950 | -0.151 | -11.683 | -26.601 | -18.957 | 0.000 |
| PC2 | 10.398 | 1.814 | 0.069 | 5.733 | 6.843 | 13.953 | 0.000 |
| PC3 | -10.865 | 1.763 | -0.072 | -6.163 | -14.321 | -7.409 | 0.000 |
| PC4 | -1.672 | 1.761 | -0.011 | -0.949 | -5.124 | 1.781 | 0.343 |
| PC5 | -12.518 | 1.760 | -0.083 | -7.111 | -15.969 | -9.067 | 0.000 |
| PC6 | -3.273 | 1.759 | -0.022 | -1.861 | -6.722 | 0.176 | 0.063 |

**Table S18.** Association between PRS Psychosis and cumulative adversity

| | Coef | Std Err | Std Coef | t | Lower 95% CI | Upper 95% CI | p |
|---|---|---|---|---|---|---|---|
| (Intercept) | 2.352 | 0.433 | | 5.429 | 1.503 | 3.201 | 0.000 |
| prsPsychosis | 0.026 | 0.010 | 0.031 | 2.528 | 0.006 | 0.046 | 0.011 |
| phenoPsychosis | 0.023 | 0.002 | 0.123 | 10.082 | 0.018 | 0.027 | 0.000 |
| age | -0.004 | 0.003 | -0.016 | -1.280 | -0.010 | 0.002 | 0.201 |
| sex | 0.029 | 0.045 | | 0.631 | -0.060 | 0.118 | 0.528 |
| PC1 | -25.726 | 1.847 | -0.170 | -13.932 | -29.346 | -22.106 | 0.000 |
| PC2 | 9.944 | 1.842 | 0.066 | 5.398 | 6.333 | 13.555 | 0.000 |
| PC3 | -10.496 | 1.837 | -0.069 | -5.715 | -14.097 | -6.896 | 0.000 |
| PC4 | -1.006 | 1.830 | -0.007 | -0.550 | -4.594 | 2.581 | 0.582 |
| PC5 | -14.029 | 1.842 | -0.093 | -7.615 | -17.640 | -10.417 | 0.000 |
| PC6 | -4.023 | 1.832 | -0.027 | -2.196 | -7.614 | -0.432 | 0.028 |

**Table S19.** Association of general psychopathology PRS and adversity with the general psychopathology phenotype

| Model | | Coef | Std Err | Std Coef | t | Lower 95% CI | Upper 95% CI | p |
|---|---|---|---|---|---|---|---|---|
| $H_0$ | *(Intercept)* | 0.844 | 0.192 | | 4.393 | 0.468 | 1.221 | 0.000 |
| | prsCVgeneral | 0.116 | 0.013 | 0.116 | 8.768 | 0.090 | 0.142 | 0.000 |
| | adversity | 0.130 | 0.007 | 0.243 | 19.800 | 0.117 | 0.143 | 0.000 |
| | age | -0.008 | 0.002 | -0.056 | -4.754 | -0.011 | -0.004 | 0.000 |
| | sex | -0.262 | 0.024 | | -11.005 | -0.308 | -0.215 | 0.000 |
| | PC1 | -0.697 | 1.063 | -0.009 | -0.656 | -2.780 | 1.386 | 0.512 |
| | PC2 | -1.044 | 0.967 | -0.013 | -1.080 | -2.940 | 0.852 | 0.280 |
| | PC3 | -3.128 | 0.967 | -0.039 | -3.235 | -5.023 | -1.233 | 0.001 |
| | PC4 | -0.444 | 0.955 | -0.006 | -0.465 | -2.315 | 1.428 | 0.642 |
| | PC5 | -0.439 | 0.968 | -0.005 | -0.453 | -2.336 | 1.459 | 0.650 |
| | PC6 | -0.280 | 0.960 | -0.003 | -0.292 | -2.161 | 1.601 | 0.770 |
| $H_1$ | *(Intercept)* | 0.846 | 0.192 | | 4.400 | 0.469 | 1.223 | 0.000 |
| | prsCVgeneral | 0.123 | 0.016 | 0.123 | 7.744 | 0.092 | 0.155 | 0.000 |
| | adversity | 0.131 | 0.007 | 0.245 | 19.591 | 0.118 | 0.144 | 0.000 |
| | age | -0.008 | 0.002 | -0.057 | -4.760 | -0.011 | -0.004 | 0.000 |
| | sex | -0.262 | 0.024 | | -10.994 | -0.308 | -0.215 | 0.000 |
| | PC1 | -0.765 | 1.066 | -0.009 | -0.718 | -2.854 | 1.324 | 0.473 |
| | PC2 | -0.975 | 0.971 | -0.012 | -1.004 | -2.878 | 0.928 | 0.315 |
| | PC3 | -3.148 | 0.967 | -0.039 | -3.255 | -5.044 | -1.252 | 0.001 |
| | PC4 | -0.438 | 0.955 | -0.005 | -0.459 | -2.310 | 1.433 | 0.646 |
| | PC5 | -0.464 | 0.968 | -0.006 | -0.479 | -2.362 | 1.435 | 0.632 |
| | PC6 | -0.277 | 0.960 | -0.003 | -0.288 | -2.158 | 1.604 | 0.773 |
| | prsCVgeneral ✻ adversity | -0.005 | 0.006 | -0.013 | -0.842 | -0.018 | 0.007 | 0.400 |

| Model | R | $R^2$ | Adjusted $R^2$ | RMSE | $R^2$ Change | df1 | df2 | p |
|---|---|---|---|---|---|---|---|---|
| $H_0$ | 0.326 | 0.106 | 0.105 | 0.948 | 0.106 | 10 | 6364 | 0.000 |
| $H_1$ | 0.326 | 0.106 | 0.105 | 0.948 | 0.000 | 1 | 6363 | 0.400 |

*Notes*. prsCVgeneral= PRS component scores from the first canonical variate representing general psychopathology. $H_1$ model includes interaction between prsCVgeneral and adversity.

**Table S20.** Association of neurodevelopmental-specific PRS and adversity with the general psychopathology phenotype

| Model | | Coef | Std Err | Std Coef | t | Lower 95% CI | Upper 95% CI | p |
|---|---|---|---|---|---|---|---|---|
| $H_0$ | *(Intercept)* | 0.515 | 0.199 | | 2.589 | 0.125 | 0.905 | 0.010 |
| | prsCVneurodev | 0.084 | 0.013 | 0.084 | 6.409 | 0.058 | 0.109 | 0.000 |
| | adversity | -0.051 | 0.007 | -0.096 | -7.574 | -0.064 | -0.038 | 0.000 |
| | age | -0.003 | 0.002 | -0.021 | -1.667 | -0.006 | 0.000 | 0.096 |
| | sex | -0.230 | 0.025 | | -9.349 | -0.279 | -0.182 | 0.000 |
| | PC1 | -1.694 | 1.031 | -0.021 | -1.644 | -3.715 | 0.327 | 0.100 |
| | PC2 | 2.763 | 1.023 | 0.034 | 2.702 | 0.758 | 4.768 | 0.007 |
| | PC3 | 2.139 | 0.997 | 0.027 | 2.145 | 0.184 | 4.094 | 0.032 |
| | PC4 | -2.541 | 1.001 | -0.032 | -2.539 | -4.504 | -0.579 | 0.011 |
| | PC5 | -0.665 | 0.999 | -0.008 | -0.665 | -2.623 | 1.294 | 0.506 |
| | PC6 | 1.598 | 0.993 | 0.020 | 1.610 | -0.348 | 3.545 | 0.108 |
| $H_1$ | *(Intercept)* | 0.519 | 0.199 | | 2.608 | 0.129 | 0.909 | 0.009 |
| | prsCVneurodev | 0.057 | 0.016 | 0.057 | 3.529 | 0.025 | 0.089 | 0.000 |
| | adversity | -0.050 | 0.007 | -0.094 | -7.378 | -0.063 | -0.037 | 0.000 |
| | age | -0.003 | 0.002 | -0.021 | -1.691 | -0.006 | 0.000 | 0.091 |
| | sex | -0.229 | 0.025 | | -9.308 | -0.277 | -0.181 | 0.000 |
| | PC1 | -1.741 | 1.030 | -0.022 | -1.690 | -3.761 | 0.279 | 0.091 |
| | PC2 | 2.705 | 1.022 | 0.033 | 2.645 | 0.700 | 4.709 | 0.008 |
| | PC3 | 2.030 | 0.998 | 0.025 | 2.035 | 0.075 | 3.986 | 0.042 |
| | PC4 | -2.624 | 1.001 | -0.033 | -2.622 | -4.586 | -0.662 | 0.009 |
| | PC5 | -0.585 | 0.999 | -0.007 | -0.586 | -2.544 | 1.373 | 0.558 |
| | PC6 | 1.600 | 0.992 | 0.020 | 1.612 | -0.345 | 3.546 | 0.107 |
| | prsCVneurodev ✻ adversity | 0.019 | 0.007 | 0.043 | 2.759 | 0.005 | 0.032 | 0.006 |

| Model | R | $R^2$ | Adjusted $R^2$ | RMSE | $R^2$ Change | df1 | df2 | p |
|---|---|---|---|---|---|---|---|---|
| $H_0$ | 0.185 | 0.034 | 0.033 | 0.981 | 0.034 | 10 | 6364 | 0.000 |
| $H_1$ | 0.189 | 0.036 | 0.034 | 0.981 | 0.001 | 1 | 6363 | 0.006 |

*Notes*. prsCVneurodev= PRS component scores from the second canonical variate representing neurodevelopmental-specific variance. phenoCVneurodev= Phenotype component scores from the second canonical variate representing neurodevelopmental-specific variance. $H_1$ model includes interaction between prsCVneurodev and adversity.

**Table S21.** Association of PRS ADHD and adversity with the ADHD phenotype

| Model | | Coef | Std Err | Std Coef | t | Lower 95% CI | Upper 95% CI | p |
|---|---|---|---|---|---|---|---|---|
| $H_0$ | *(Intercept)* | 8.077 | 0.782 | | 10.326 | 6.544 | 9.611 | 0.000 |
| | prsADHD | 0.082 | 0.011 | 0.094 | 7.445 | 0.060 | 0.103 | 0.000 |
| | adversity | 0.388 | 0.020 | 0.242 | 19.724 | 0.349 | 0.427 | 0.000 |
| | age | -0.014 | 0.005 | -0.034 | -2.845 | -0.023 | -0.004 | 0.004 |
| | sex | -1.007 | 0.072 | | -14.078 | -1.147 | -0.867 | 0.000 |
| | PC1 | 4.904 | 3.054 | 0.020 | 1.606 | -1.082 | 10.891 | 0.108 |
| | PC2 | 5.853 | 2.895 | 0.024 | 2.021 | 0.177 | 11.529 | 0.043 |
| | PC3 | -1.431 | 2.910 | -0.006 | -0.492 | -7.136 | 4.273 | 0.623 |
| | PC4 | -7.519 | 2.875 | -0.031 | -2.615 | -13.155 | -1.883 | 0.009 |
| | PC5 | -2.253 | 2.909 | -0.009 | -0.775 | -7.955 | 3.449 | 0.439 |
| | PC6 | 0.547 | 2.885 | 0.002 | 0.190 | -5.109 | 6.203 | 0.850 |
| $H_1$ | *(Intercept)* | 7.861 | 0.871 | | 9.027 | 6.154 | 9.568 | 0.000 |
| | prsADHD | 0.077 | 0.014 | 0.089 | 5.678 | 0.051 | 0.104 | 0.000 |
| | adversity | 0.542 | 0.273 | 0.338 | 1.986 | 0.007 | 1.077 | 0.047 |
| | age | -0.014 | 0.005 | -0.034 | -2.842 | -0.023 | -0.004 | 0.004 |
| | sex | -1.007 | 0.072 | | -14.079 | -1.148 | -0.867 | 0.000 |
| | PC1 | 5.007 | 3.059 | 0.021 | 1.637 | -0.990 | 11.004 | 0.102 |
| | PC2 | 5.724 | 2.904 | 0.024 | 1.971 | 0.030 | 11.418 | 0.049 |
| | PC3 | -1.424 | 2.910 | -0.006 | -0.489 | -7.129 | 4.282 | 0.625 |
| | PC4 | -7.545 | 2.876 | -0.031 | -2.624 | -13.182 | -1.908 | 0.009 |
| | PC5 | -2.196 | 2.911 | -0.009 | -0.754 | -7.901 | 3.510 | 0.451 |
| | PC6 | 0.544 | 2.885 | 0.002 | 0.189 | -5.112 | 6.201 | 0.850 |
| | prsADHD ✻ adversity | 0.003 | 0.006 | 0.096 | 0.566 | -0.008 | 0.015 | 0.571 |

| Model | R | $R^2$ | Adjusted $R^2$ | RMSE | $R^2$ Change | df1 | df2 | p |
|---|---|---|---|---|---|---|---|---|
| $H_0$ | 0.320 | 0.103 | 0.101 | 2.850 | 0.103 | 10 | 6366 | 0.000 |
| $H_1$ | 0.320 | 0.103 | 0.101 | 2.850 | 0.000 | 1 | 6365 | 0.571 |

*Notes*. $H_1$ model includes interaction between prsADHD and adversity.

**Table S22.** Association of PRS Anxiety and adversity with the Anxiety phenotype

| Model | | Coef | Std Err | Std Coef | t | Lower 95% CI | Upper 95% CI | p |
|---|---|---|---|---|---|---|---|---|
| $H_0$ | *(Intercept)* | 2.674 | 0.621 | | 4.307 | 1.457 | 3.891 | 0.000 |
| | prsAnxiety | 0.023 | 0.010 | 0.029 | 2.246 | 0.003 | 0.042 | 0.025 |
| | adversity | 0.331 | 0.016 | 0.251 | 20.061 | 0.298 | 0.363 | 0.000 |
| | age | -0.001 | 0.004 | -0.003 | -0.282 | -0.009 | 0.007 | 0.778 |
| | sex | -0.025 | 0.060 | | -0.423 | -0.143 | 0.092 | 0.672 |
| | PC1 | 8.526 | 2.533 | 0.043 | 3.366 | 3.561 | 13.490 | 0.001 |
| | PC2 | -0.571 | 2.454 | -0.003 | -0.233 | -5.383 | 4.240 | 0.816 |
| | PC3 | -1.884 | 2.434 | -0.009 | -0.774 | -6.655 | 2.887 | 0.439 |
| | PC4 | -0.871 | 2.422 | -0.004 | -0.360 | -5.619 | 3.877 | 0.719 |
| | PC5 | -0.550 | 2.432 | -0.003 | -0.226 | -5.318 | 4.217 | 0.821 |
| | PC6 | -3.500 | 2.421 | -0.018 | -1.445 | -8.247 | 1.247 | 0.148 |
| $H_1$ | *(Intercept)* | 3.005 | 0.681 | | 4.416 | 1.671 | 4.340 | 0.000 |
| | prsAnxiety | 0.031 | 0.013 | 0.040 | 2.513 | 0.007 | 0.056 | 0.012 |
| | adversity | 0.104 | 0.191 | 0.079 | 0.543 | -0.271 | 0.479 | 0.587 |
| | age | -0.001 | 0.004 | -0.003 | -0.286 | -0.009 | 0.007 | 0.775 |
| | sex | -0.025 | 0.060 | | -0.421 | -0.143 | 0.092 | 0.673 |
| | PC1 | 8.474 | 2.533 | 0.042 | 3.345 | 3.508 | 13.439 | 0.001 |
| | PC2 | -0.537 | 2.455 | -0.003 | -0.219 | -5.349 | 4.274 | 0.827 |
| | PC3 | -1.933 | 2.434 | -0.010 | -0.794 | -6.704 | 2.838 | 0.427 |
| | PC4 | -0.909 | 2.422 | -0.005 | -0.375 | -5.657 | 3.839 | 0.707 |
| | PC5 | -0.516 | 2.432 | -0.003 | -0.212 | -5.283 | 4.252 | 0.832 |
| | PC6 | -3.536 | 2.422 | -0.018 | -1.460 | -8.283 | 1.211 | 0.144 |
| | prsAnxiety ✻ adversity | -0.006 | 0.005 | -0.172 | -1.187 | -0.016 | 0.004 | 0.235 |

| Model | R | $R^2$ | Adjusted $R^2$ | RMSE | $R^2$ Change | df1 | df2 | p |
|---|---|---|---|---|---|---|---|---|
| $H_0$ | 0.252 | 0.064 | 0.062 | 2.393 | 0.064 | 10 | 6366 | 0.000 |
| $H_1$ | 0.253 | 0.064 | 0.062 | 2.392 | 0.000 | 1 | 6365 | 0.235 |

*Notes*. $H_1$ model includes interaction between prsAnxiety and adversity.

**Table S23.** Association of PRS Depression and adversity with the Depression phenotype

| Model | | Coef | Std Err | Std Coef | t | Lower 95% CI | Upper 95% CI | p |
|---|---|---|---|---|---|---|---|---|
| $H_0$ | *(Intercept)* | 1.806 | 0.541 | | 3.339 | 0.746 | 2.866 | 0.001 |
| | prsDepression | 0.052 | 0.011 | 0.061 | 4.508 | 0.029 | 0.074 | 0.000 |
| | adversity | 0.328 | 0.013 | 0.303 | 24.737 | 0.302 | 0.354 | 0.000 |
| | age | 0.007 | 0.003 | 0.025 | 2.081 | 0.000 | 0.013 | 0.037 |
| | sex | -0.276 | 0.048 | | -5.707 | -0.371 | -0.181 | 0.000 |
| | PC1 | 8.358 | 2.182 | 0.051 | 3.831 | 4.081 | 12.636 | 0.000 |
| | PC2 | 1.963 | 2.015 | 0.012 | 0.974 | -1.988 | 5.914 | 0.330 |
| | PC3 | -0.680 | 1.960 | -0.004 | -0.347 | -4.523 | 3.162 | 0.728 |
| | PC4 | -1.224 | 1.952 | -0.007 | -0.627 | -5.051 | 2.603 | 0.531 |
| | PC5 | -0.128 | 1.959 | -0.001 | -0.065 | -3.968 | 3.713 | 0.948 |
| | PC6 | -1.843 | 1.950 | -0.011 | -0.945 | -5.666 | 1.981 | 0.345 |
| $H_1$ | *(Intercept)* | 1.531 | 0.593 | | 2.582 | 0.369 | 2.694 | 0.010 |
| | prsDepression | 0.043 | 0.014 | 0.051 | 3.153 | 0.016 | 0.070 | 0.002 |
| | adversity | 0.520 | 0.170 | 0.480 | 3.051 | 0.186 | 0.854 | 0.002 |
| | age | 0.007 | 0.003 | 0.025 | 2.096 | 0.000 | 0.013 | 0.036 |
| | sex | -0.277 | 0.048 | | -5.732 | -0.372 | -0.182 | 0.000 |
| | PC1 | 8.516 | 2.186 | 0.052 | 3.895 | 4.230 | 12.801 | 0.000 |
| | PC2 | 1.863 | 2.017 | 0.011 | 0.924 | -2.092 | 5.818 | 0.356 |
| | PC3 | -0.574 | 1.962 | -0.003 | -0.292 | -4.420 | 3.273 | 0.770 |
| | PC4 | -1.188 | 1.952 | -0.007 | -0.609 | -5.016 | 2.639 | 0.543 |
| | PC5 | -0.116 | 1.959 | -0.001 | -0.059 | -3.957 | 3.724 | 0.953 |
| | PC6 | -1.864 | 1.951 | -0.011 | -0.956 | -5.688 | 1.959 | 0.339 |
| | prsDepression ✻ adversity | 0.006 | 0.005 | 0.177 | 1.129 | -0.004 | 0.017 | 0.259 |

| Model | R | $R^2$ | Adjusted $R^2$ | RMSE | $R^2$ Change | df1 | df2 | p |
|---|---|---|---|---|---|---|---|---|
| $H_0$ | 0.316 | 0.100 | 0.099 | 1.927 | 0.100 | 10 | 6366 | 0.000 |
| $H_1$ | 0.317 | 0.100 | 0.099 | 1.927 | 0.000 | 1 | 6365 | 0.259 |

*Notes*. $H_1$ model includes interaction between prsDepression and adversity.

**Table S24.** Association of PRS Psychosis and adversity with the Psychosis phenotype

| Model | | Coef | Std Err | Std Coef | t | Lower 95% CI | Upper 95% CI | p |
|---|---|---|---|---|---|---|---|---|
| $H_0$ | *(Intercept)* | 18.267 | 2.397 | | 7.621 | 13.568 | 22.965 | 0.000 |
| | prsPsychosis | 0.166 | 0.056 | 0.037 | 2.959 | 0.056 | 0.277 | 0.003 |
| | adversity | 0.695 | 0.069 | 0.128 | 10.082 | 0.560 | 0.830 | 0.000 |
| | age | -0.079 | 0.017 | -0.058 | -4.670 | -0.112 | -0.046 | 0.000 |
| | sex | -0.533 | 0.252 | | -2.118 | -1.027 | -0.040 | 0.034 |
| | PC1 | -58.792 | 10.367 | -0.071 | -5.671 | -79.114 | -38.470 | 0.000 |
| | PC2 | -48.825 | 10.219 | -0.059 | -4.778 | -68.857 | -28.793 | 0.000 |
| | PC3 | -38.411 | 10.197 | -0.046 | -3.767 | -58.400 | -18.422 | 0.000 |
| | PC4 | 21.282 | 10.143 | 0.026 | 2.098 | 1.398 | 41.167 | 0.036 |
| | PC5 | 3.117 | 10.260 | 0.004 | 0.304 | -16.996 | 23.230 | 0.761 |
| | PC6 | -5.519 | 10.160 | -0.007 | -0.543 | -25.436 | 14.397 | 0.587 |
| $H_1$ | *(Intercept)* | 20.919 | 2.584 | | 8.097 | 15.854 | 25.983 | 0.000 |
| | prsPsychosis | 0.285 | 0.071 | 0.063 | 4.017 | 0.146 | 0.424 | 0.000 |
| | adversity | -1.144 | 0.675 | -0.210 | -1.696 | -2.467 | 0.178 | 0.090 |
| | age | -0.079 | 0.017 | -0.057 | -4.647 | -0.112 | -0.046 | 0.000 |
| | sex | -0.533 | 0.252 | | -2.116 | -1.026 | -0.039 | 0.034 |
| | PC1 | -59.836 | 10.368 | -0.072 | -5.771 | -80.162 | -39.511 | 0.000 |
| | PC2 | -47.153 | 10.232 | -0.057 | -4.609 | -67.211 | -27.095 | 0.000 |
| | PC3 | -38.122 | 10.192 | -0.046 | -3.740 | -58.102 | -18.142 | 0.000 |
| | PC4 | 21.731 | 10.140 | 0.026 | 2.143 | 1.854 | 41.608 | 0.032 |
| | PC5 | 2.548 | 10.257 | 0.003 | 0.248 | -17.559 | 22.654 | 0.804 |
| | PC6 | -5.955 | 10.156 | -0.007 | -0.586 | -25.864 | 13.954 | 0.558 |
| | prsPsychosis ✻ adversity | -0.081 | 0.030 | -0.339 | -2.741 | -0.139 | -0.023 | 0.006 |

| Model | R | $R^2$ | Adjusted $R^2$ | RMSE | $R^2$ Change | df1 | df2 | p |
|---|---|---|---|---|---|---|---|---|
| $H_0$ | 0.194 | 0.037 | 0.036 | 10.031 | 0.037 | 10 | 6365 | 0.000 |
| $H_1$ | 0.196 | 0.039 | 0.037 | 10.026 | 0.001 | 1 | 6364 | 0.006 |

*Notes*. $H_1$ model includes interaction between prsPsychosis and adversity.

**Figure S8.** Prediction of mental health phenotypes by PRS components, adversity, and their interaction

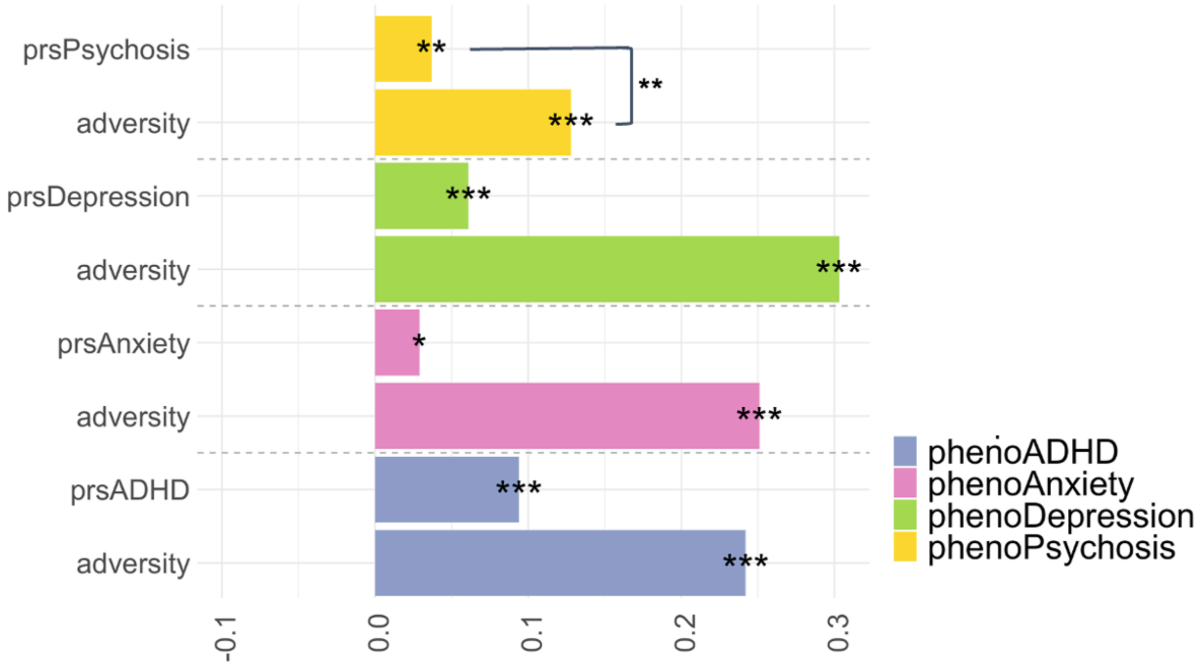


*Notes*. Regressions model the predictive value for each phenotype separately, including age, sex, and the first 6 population components (PCs) as covariates. Y-axis represents standardised coefficients. Only significant interactions shown.

**Table S25.** PLS: cortico-limbic network connectivity VIP scores and loadings for the joint model

| | VIP | Loading | Std. err | t | p | 2.50% | 97.50% |
|---|---|---|---|---|---|---|---|
| *Predictor (adversity & PRS)* | | | | | | | |
| **prsDepression** | **1.334** | **-0.597** | **0.086** | **-6.796** | **0.000** | **-0.710** | **-0.444** |
| **adversity** | **1.280** | **-0.572** | **0.100** | **-5.541** | **0.000** | **-0.705** | **-0.387** |
| **prsADHD** | **1.107** | **-0.495** | **0.084** | **-5.815** | **0.000** | **-0.625** | **-0.329** |
| prsAnxiety | 0.594 | -0.266 | 0.089 | -2.895 | 0.004 | -0.421 | -0.074 |
| prsPsychosis | 0.055 | -0.024 | 0.104 | -0.225 | 0.822 | -0.219 | 0.187 |
| *Outcome (Network connectivity)* | | | | | | | |
| **Cingulo opercular** | **1.499** | **0.433** | **0.048** | **8.840** | **0.000** | **0.368** | **0.483** |
| **Sensorimotor hand** | **1.467** | **0.423** | **0.049** | **8.526** | **0.000** | **0.358** | **0.478** |
| **Cingulo parietal** | **1.281** | **0.370** | **0.047** | **7.705** | **0.000** | **0.298** | **0.428** |
| **Visual** | **1.166** | **0.337** | **0.047** | **7.058** | **0.000** | **0.260** | **0.402** |
| **Default** | **1.040** | **0.300** | **0.045** | **6.557** | **0.000** | **0.223** | **0.372** |
| **Retrosplenial temporal** | **1.011** | **0.292** | **0.057** | **5.055** | **0.000** | **0.184** | **0.382** |
| Sensorimotor mouth | 0.987 | 0.285 | 0.046 | 6.080 | 0.000 | 0.197 | 0.353 |
| Salience | 0.635 | 0.183 | 0.048 | 3.740 | 0.000 | 0.094 | 0.266 |
| Ventral attention | 0.615 | 0.177 | 0.048 | 3.619 | 0.000 | 0.078 | 0.261 |
| Auditory | 0.590 | 0.170 | 0.059 | 2.847 | 0.004 | 0.058 | 0.285 |
| Fronto-parietal | 0.510 | 0.147 | 0.048 | 3.026 | 0.002 | 0.052 | 0.238 |
| Dorsal attention | 0.369 | 0.106 | 0.052 | 2.050 | 0.040 | 0.004 | 0.202 |

*Notes*. Variable importance in projection (VIP) was used to assess the relative importance of each item in the PLS model. Bold indicates VIP scores considered most influential in terms of their explanatory power (>1). Bootstrapping with 1,000 resampling iterations was used to obtain standard errors, t-values confidence intervals, and p-values.

**Table S26.** PLS: cortico-limbic network connectivity VIP scores and loadings for the PRS-only model

| | VIP | Loading | Std. err | t | p | 2.50% | 97.50% |
|---|---|---|---|---|---|---|---|
| *Predictor (PRS)* | | | | | | | |
| **prsDepression** | **1.457** | **-0.728** | **0.069** | **-10.265** | **0.000** | **-0.839** | **-0.562** |
| **prsADHD** | **1.205** | **-0.602** | **0.086** | **-6.906** | **0.000** | **-0.753** | **-0.423** |
| prsAnxiety | 0.649 | -0.324 | 0.101 | -3.116 | 0.002 | -0.501 | -0.102 |
| prsPsychosis | 0.069 | -0.034 | 0.129 | -0.262 | 0.794 | -0.266 | 0.226 |
| *Outcome (Network connectivity)* | | | | | | | |
| **Cingulo opercular** | **1.534** | **0.443** | **0.034** | **12.628** | **0.000** | **0.367** | **0.501** |
| **Sensorimotor hand** | **1.424** | **0.411** | **0.036** | **11.267** | **0.000** | **0.333** | **0.472** |
| **Cingulo parietal** | **1.269** | **0.366** | **0.042** | **8.636** | **0.000** | **0.273** | **0.437** |
| **Visual** | **1.154** | **0.333** | **0.044** | **7.387** | **0.000** | **0.237** | **0.406** |
| **Default** | **1.042** | **0.301** | **0.045** | **6.594** | **0.000** | **0.198** | **0.382** |
| **Retrosplenial temporal** | **1.033** | **0.298** | **0.063** | **4.628** | **0.000** | **0.164** | **0.412** |
| **Sensorimotor mouth** | **1.005** | **0.290** | **0.049** | **5.764** | **0.000** | **0.184** | **0.376** |
| Ventral attention | 0.652 | 0.188 | 0.055 | 3.348 | 0.001 | 0.075 | 0.286 |
| Auditory | 0.640 | 0.185 | 0.071 | 2.575 | 0.010 | 0.054 | 0.330 |
| Salience | 0.591 | 0.171 | 0.055 | 3.044 | 0.002 | 0.045 | 0.268 |
| Fronto-parietal | 0.491 | 0.142 | 0.058 | 2.404 | 0.016 | 0.021 | 0.252 |
| Dorsal attention | 0.296 | 0.086 | 0.063 | 1.364 | 0.173 | -0.044 | 0.203 |

*Notes*. Variable importance in projection (VIP) was used to assess the relative importance of each adversity item in the PLS model. Bold indicates VIP scores considered most influential in terms of their explanatory power (>1). Bootstrapping with 1,000 resampling iterations was used to obtain standard errors, t-values confidence intervals, and p-values.

**Table S27.** PLS: cortico-limbic network connectivity VIP scores and loadings for the adversity-only model

| | VIP | Loading | Std. err | t | p | 2.50% | 97.50% |
|---|---|---|---|---|---|---|---|
| *Predictor (adversity)* | | | | | | | |
| **adversity** | **-** | **1.000** | **0.000** | **-** | **0.000** | **1** | **1** |
| *Outcome (Network connectivity)* | | | | | | | |
| **Sensorimotor hand** | **1.549** | **-0.447** | **0.046** | **-9.644** | **0.000** | **-0.531** | **-0.359** |
| **Cingulo opercular** | **1.420** | **-0.410** | **0.045** | **-8.860** | **0.000** | **-0.487** | **-0.310** |
| **Cingulo parietal** | **1.298** | **-0.375** | **0.044** | **-8.312** | **0.000** | **-0.448** | **-0.269** |
| **Visual** | **1.185** | **-0.342** | **0.053** | **-6.309** | **0.000** | **-0.439** | **-0.225** |
| **Default** | **1.032** | **-0.298** | **0.061** | **-4.768** | **0.000** | **-0.407** | **-0.168** |
| Retrosplenial temporal | 0.966 | -0.279 | 0.070 | -3.933 | 0.000 | -0.418 | -0.145 |
| Sensorimotor mouth | 0.943 | -0.272 | 0.056 | -4.703 | 0.000 | -0.361 | -0.148 |
| Salience | 0.720 | -0.208 | 0.063 | -3.183 | 0.001 | -0.319 | -0.070 |
| Fronto-parietal | 0.547 | -0.158 | 0.068 | -2.190 | 0.029 | -0.269 | -0.004 |
| Ventral attention | 0.534 | -0.154 | 0.067 | -2.192 | 0.028 | -0.271 | -0.006 |
| Dorsal attention | 0.514 | -0.148 | 0.069 | -2.047 | 0.041 | -0.268 | 0.006 |
| Auditory | 0.489 | -0.141 | 0.075 | -1.864 | 0.062 | -0.289 | 0.001 |

*Notes*. Variable importance in projection (VIP) was used to assess the relative importance of each item in the PLS model. Bold indicates VIP scores considered most influential in terms of their explanatory power (>1). Bootstrapping with 1,000 resampling iterations was used to obtain standard errors, t-values confidence intervals, and p-values.

**Table S28.** Mediation effect of the PLS cortico-limbic network connectivity component on the association between the adversity component and mental health phenotypes

| *Direct Effects* | | | | | Est | Std Err | z | Lower 95% | Upper 95% | p |
|---|---|---|---|---|---|---|---|---|---|---|
| adversity component | | → | | phenoADHD | 0.716 | 0.048 | 14.892 | 0.622 | 0.810 | 0.000 |
| adversity component | | → | | phenoAnxiety | 0.638 | 0.043 | 14.984 | 0.554 | 0.721 | 0.000 |
| adversity component | | → | | phenoDepression | 0.604 | 0.033 | 18.053 | 0.538 | 0.670 | 0.000 |
| adversity component | | → | | phenoPsychosis | 1.093 | 0.162 | 6.731 | 0.775 | 1.411 | 0.000 |
| *Indirect Effects* | | | | | **Est** | **Std Err** | **z** | **Lower 95%** | **Upper 95%** | **p** |
| adversity component | → | cortico-limbic component | → | phenoADHD | -0.007 | 0.004 | -1.840 | -0.014 | 0.000 | 0.066 |
| adversity component | → | cortico-limbic component | → | phenoAnxiety | -0.006 | 0.003 | -1.830 | -0.012 | 0.000 | 0.067 |
| adversity component | → | cortico-limbic component | → | phenoDepression | -0.006 | 0.003 | -2.146 | -0.011 | 0.000 | 0.032 |
| adversity component | → | cortico-limbic component | → | phenoPsychosis | 0.027 | 0.013 | 2.130 | 0.002 | 0.052 | 0.033 |

*Notes*. Components derived from the adversity-only cortico-limbic network connectivity PLS model. Controlling for age, sex and 6 PCs.

**Table S29.** PLS: cortico-limbic regional connectivity VIP scores and loadings for the joint model

| | VIP | Loading | Std. err | t | p | 2.50% | 97.50% |
|---|---|---|---|---|---|---|---|
| *Predictor (adversity & PRS)* | | | | | | | |
| **prsDepression** | **1.296** | **-0.579** | **0.052** | **-11.014** | **0.000** | **-0.673** | **-0.467** |
| **adversity** | **1.257** | **-0.562** | **0.068** | **-7.981** | **0.000** | **-0.679** | **-0.413** |
| **prsADHD** | **1.162** | **-0.520** | **0.054** | **-9.483** | **0.000** | **-0.619** | **-0.403** |
| prsAnxiety | 0.623 | -0.279 | 0.070 | -4.148 | 0.000 | -0.414 | -0.141 |
| prsPsychosis | 0.052 | -0.023 | 0.080 | -0.476 | 0.634 | -0.192 | 0.128 |
| *Outcome (regional connectivity)* | | | | | | | |
| **sensorimotor hand : left-pallidum** | **2.406** | **0.159** | **0.012** | **12.353** | **0.000** | **0.128** | **0.176** |
| **sensorimotor hand : right-accumbens** | **2.360** | **0.156** | **0.014** | **10.975** | **0.000** | **0.122** | **0.176** |
| **cingulo-parietal : left-thalamus** | **2.334** | **0.155** | **0.014** | **10.653** | **0.000** | **0.119** | **0.176** |
| **cingulo-opercular : left-amygdala** | **2.328** | **0.154** | **0.013** | **11.164** | **0.000** | **0.121** | **0.171** |
| **auditory : right-putamen** | **2.302** | **0.152** | **0.012** | **12.037** | **0.000** | **0.122** | **0.170** |
| **visual : left-hippocampus** | **2.153** | **0.143** | **0.016** | **8.553** | **0.000** | **0.107** | **0.167** |
| **sensorimotor hand : brain-stem** | **2.112** | **0.140** | **0.016** | **8.518** | **0.000** | **0.104** | **0.167** |
| **sensorimotor hand : right-caudate** | **2.066** | **0.137** | **0.014** | **9.355** | **0.000** | **0.104** | **0.158** |
| **retrosplenial temporal : right-thalamus** | **2.061** | **0.136** | **0.017** | **7.565** | **0.000** | **0.095** | **0.164** |
| **retrosplenial temporal : right-cerebellum** | **2.059** | **0.136** | **0.013** | **9.983** | **0.000** | **0.106** | **0.157** |
| **sensorimotor hand : right-putamen** | **2.057** | **0.136** | **0.016** | **8.219** | **0.000** | **0.099** | **0.164** |
| **sensorimotor mouth : right-caudate** | **2.024** | **0.134** | **0.018** | **7.118** | **0.000** | **0.091** | **0.162** |
| **sensorimotor mouth : right-thalamus** | **2.012** | **0.133** | **0.015** | **8.718** | **0.000** | **0.098** | **0.157** |
| **cingulo-opercular : right-amygdala** | **2.003** | **0.133** | **0.016** | **8.044** | **0.000** | **0.096** | **0.160** |
| **sensorimotor hand : right-ventraldc** | **1.928** | **0.128** | **0.018** | **6.835** | **0.000** | **0.086** | **0.156** |
| **cingulo-opercular : left-putamen** | **1.906** | **0.126** | **0.014** | **8.427** | **0.000** | **0.092** | **0.149** |
| **cingulo-parietal : left-caudate** | **1.874** | **0.124** | **0.018** | **6.581** | **0.000** | **0.083** | **0.152** |
| **cingulo-opercular : right-hippocampus** | **1.872** | **0.124** | **0.015** | **8.176** | **0.000** | **0.091** | **0.147** |
| **salience : left-thalamus** | **1.869** | **0.124** | **0.016** | **7.454** | **0.000** | **0.085** | **0.147** |

| | | | | | | | |
|---|---|---|---|---|---|---|---|
| **cingulo-opercular : brain-stem** | **1.823** | **0.121** | **0.018** | **6.394** | **0.000** | **0.081** | **0.153** |
| **cingulo-parietal : right-pallidum** | **1.814** | **0.120** | **0.016** | **7.394** | **0.000** | **0.084** | **0.147** |
| **auditory : right-pallidum** | **1.802** | **0.119** | **0.017** | **6.830** | **0.000** | **0.082** | **0.147** |
| **cingulo-opercular : left-accumbens** | **1.775** | **0.118** | **0.017** | **6.808** | **0.000** | **0.079** | **0.145** |
| **ventral-attention : left-caudate** | **1.760** | **0.117** | **0.018** | **6.115** | **0.000** | **0.077** | **0.145** |
| **cingulo-opercular : left-caudate** | **1.735** | **0.115** | **0.015** | **7.626** | **0.000** | **0.081** | **0.137** |
| **default : left-pallidum** | **1.707** | **0.113** | **0.017** | **6.404** | **0.000** | **0.075** | **0.141** |
| **cingulo-parietal : left-ventraldc** | **1.686** | **0.112** | **0.017** | **6.364** | **0.000** | **0.073** | **0.139** |
| **retrosplenial temporal : right-ventraldc** | **1.674** | **0.111** | **0.015** | **6.955** | **0.000** | **0.075** | **0.135** |
| **cingulo-opercular : right-ventraldc** | **1.673** | **0.111** | **0.017** | **6.223** | **0.000** | **0.072** | **0.140** |
| **ventral-attention : left-hippocampus** | **1.673** | **0.111** | **0.019** | **5.526** | **0.000** | **0.069** | **0.144** |
| **visual : right-hippocampus** | **1.646** | **0.109** | **0.017** | **6.268** | **0.000** | **0.070** | **0.138** |
| **sensorimotor hand : left-cerebellum** | **1.628** | **0.108** | **0.015** | **7.031** | **0.000** | **0.076** | **0.133** |
| **visual : right-pallidum** | **1.606** | **0.106** | **0.017** | **5.983** | **0.000** | **0.066** | **0.134** |
| **cingulo-parietal : right-cerebellum** | **1.586** | **0.105** | **0.016** | **6.218** | **0.000** | **0.066** | **0.129** |
| **cingulo-opercular : left-ventraldc** | **1.576** | **-0.104** | **0.020** | **-5.084** | **0.000** | **-0.140** | **-0.062** |
| **sensorimotor mouth : left-amygdala** | **1.568** | **0.104** | **0.016** | **6.008** | **0.000** | **0.066** | **0.130** |
| **sensorimotor mouth : left-hippocampus** | **1.564** | **0.104** | **0.017** | **5.686** | **0.000** | **0.064** | **0.134** |
| **auditory : left-ventraldc** | **1.550** | **0.103** | **0.019** | **5.357** | **0.000** | **0.063** | **0.136** |
| **dorsal attention : left-hippocampus** | **1.530** | **-0.101** | **0.019** | **-4.999** | **0.000** | **-0.133** | **-0.059** |
| **cingulo-parietal : right-hippocampus** | **1.508** | **0.100** | **0.018** | **5.362** | **0.000** | **0.058** | **0.127** |
| **dorsal attention : right-ventraldc** | **1.496** | **0.099** | **0.019** | **5.097** | **0.000** | **0.060** | **0.133** |
| **default : left-hippocampus** | **1.490** | **0.099** | **0.020** | **4.826** | **0.000** | **0.057** | **0.133** |
| **default : left-accumbens** | **1.487** | **0.098** | **0.016** | **5.802** | **0.000** | **0.060** | **0.124** |
| **sensorimotor mouth : left-ventraldc** | **1.399** | **0.093** | **0.018** | **4.838** | **0.000** | **0.052** | **0.124** |
| **cingulo-parietal : right-caudate** | **1.399** | **0.093** | **0.018** | **4.942** | **0.000** | **0.055** | **0.123** |
| **fronto-parietal : right-hippocampus** | **1.392** | **0.092** | **0.020** | **4.396** | **0.000** | **0.047** | **0.127** |
| **visual : left-amygdala** | **1.377** | **0.091** | **0.019** | **4.529** | **0.000** | **0.050** | **0.126** |
| **ventral-attention : brain-stem** | **1.339** | **0.089** | **0.018** | **4.575** | **0.000** | **0.049** | **0.119** |

| | | | | | | | |
|---|---|---|---|---|---|---|---|
| **dorsal attention : brain-stem** | **1.319** | **0.087** | **0.019** | **4.355** | **0.000** | **0.043** | **0.120** |
| **auditory : left-putamen** | **1.311** | **-0.087** | **0.020** | **-4.218** | **0.000** | **-0.121** | **-0.043** |
| **default : left-putamen** | **1.275** | **0.084** | **0.018** | **4.477** | **0.000** | **0.044** | **0.115** |
| **auditory : left-cerebellum** | **1.257** | **0.083** | **0.019** | **4.191** | **0.000** | **0.043** | **0.121** |
| **salience : left-ventraldc** | **1.249** | **-0.083** | **0.020** | **-3.975** | **0.000** | **-0.117** | **-0.038** |
| **ventral-attention : right-ventraldc** | **1.194** | **0.079** | **0.018** | **4.083** | **0.000** | **0.038** | **0.108** |
| **salience : left-accumbens** | **1.190** | **0.079** | **0.018** | **4.071** | **0.000** | **0.036** | **0.107** |
| **salience : left-putamen** | **1.187** | **0.079** | **0.019** | **3.938** | **0.000** | **0.037** | **0.114** |
| **default : right-accumbens** | **1.180** | **0.078** | **0.018** | **4.201** | **0.000** | **0.041** | **0.109** |
| **sensorimotor hand : left-caudate** | **1.144** | **0.076** | **0.019** | **3.900** | **0.000** | **0.036** | **0.109** |
| **fronto-parietal : left-caudate** | **1.136** | **0.075** | **0.018** | **3.981** | **0.000** | **0.037** | **0.111** |
| **visual : right-accumbens** | **1.090** | **0.072** | **0.019** | **3.621** | **0.000** | **0.033** | **0.108** |
| **fronto-parietal : right-pallidum** | **1.089** | **-0.072** | **0.019** | **-3.502** | **0.000** | **-0.107** | **-0.031** |
| **ventral-attention : left-putamen** | **1.089** | **-0.072** | **0.020** | **-3.483** | **0.001** | **-0.107** | **-0.029** |
| **auditory : right-hippocampus** | **1.064** | **-0.070** | **0.019** | **-3.620** | **0.000** | **-0.108** | **-0.033** |
| **visual : left-thalamus** | **1.063** | **0.070** | **0.021** | **3.298** | **0.001** | **0.028** | **0.110** |
| **auditory : right-ventraldc** | **1.061** | **-0.070** | **0.019** | **-3.490** | **0.000** | **-0.102** | **-0.028** |
| **retrosplenial temporal : left-hippocampus** | **1.056** | **0.070** | **0.020** | **3.470** | **0.001** | **0.030** | **0.107** |
| **salience : left-cerebellum** | **1.029** | **0.068** | **0.019** | **3.516** | **0.000** | **0.029** | **0.102** |
| **sensorimotor hand : right-hippocampus** | **1.020** | **0.068** | **0.017** | **3.692** | **0.000** | **0.028** | **0.096** |
| **fronto-parietal : right-accumbens** | **1.003** | **0.066** | **0.020** | **3.294** | **0.001** | **0.025** | **0.101** |
| **visual : right-thalamus** | **1.002** | **-0.066** | **0.019** | **-3.280** | **0.001** | **-0.099** | **-0.024** |
| retrosplenial temporal : brain-stem | 0.988 | 0.065 | 0.019 | 3.349 | 0.001 | 0.026 | 0.097 |
| fronto-parietal : left-putamen | 0.979 | 0.065 | 0.019 | 3.214 | 0.001 | 0.024 | 0.096 |
| auditory : brain-stem | 0.976 | -0.065 | 0.020 | -3.177 | 0.002 | -0.101 | -0.024 |
| default : left-ventraldc | 0.963 | 0.064 | 0.019 | 3.200 | 0.001 | 0.024 | 0.101 |
| fronto-parietal : left-ventraldc | 0.950 | 0.063 | 0.019 | 3.152 | 0.002 | 0.021 | 0.100 |
| cingulo-parietal : right-amygdala | 0.927 | -0.061 | 0.021 | -2.758 | 0.006 | -0.102 | -0.019 |
| sensorimotor hand : left-amygdala | 0.908 | -0.060 | 0.020 | -2.875 | 0.004 | -0.096 | -0.018 |

| | | | | | | | |
|---|---|---|---|---|---|---|---|
| sensorimotor hand : left-ventraldc | 0.903 | -0.060 | 0.020 | -2.943 | 0.003 | -0.096 | -0.020 |
| ventral-attention : right-caudate | 0.900 | 0.060 | 0.019 | 2.976 | 0.003 | 0.018 | 0.096 |
| default : right-hippocampus | 0.900 | 0.060 | 0.020 | 2.823 | 0.005 | 0.019 | 0.096 |
| sensorimotor hand : left-hippocampus | 0.899 | -0.060 | 0.019 | -2.945 | 0.003 | -0.095 | -0.018 |
| retrosplenial temporal : right-hippocampus | 0.898 | -0.059 | 0.019 | -2.969 | 0.003 | -0.091 | -0.017 |
| retrosplenial temporal : right-accumbens | 0.887 | 0.059 | 0.020 | 2.803 | 0.005 | 0.018 | 0.096 |
| cingulo-opercular : right-putamen | 0.880 | 0.058 | 0.020 | 2.743 | 0.006 | 0.017 | 0.098 |
| ventral-attention : left-pallidum | 0.876 | -0.058 | 0.020 | -2.651 | 0.008 | -0.098 | -0.017 |
| cingulo-parietal : right-thalamus | 0.871 | 0.058 | 0.020 | 2.764 | 0.006 | 0.018 | 0.096 |
| dorsal attention : left-accumbens | 0.865 | 0.057 | 0.020 | 2.847 | 0.004 | 0.018 | 0.093 |
| fronto-parietal : left-cerebellum | 0.859 | -0.057 | 0.019 | -2.825 | 0.005 | -0.092 | -0.018 |
| cingulo-opercular : right-caudate | 0.859 | -0.057 | 0.020 | -2.748 | 0.006 | -0.094 | -0.017 |
| retrosplenial temporal : left-cerebellum | 0.845 | -0.056 | 0.019 | -2.813 | 0.005 | -0.094 | -0.015 |
| fronto-parietal : right-putamen | 0.829 | -0.055 | 0.020 | -2.579 | 0.010 | -0.090 | -0.012 |
| dorsal attention : left-amygdala | 0.811 | -0.054 | 0.021 | -2.404 | 0.016 | -0.095 | -0.012 |
| default : right-putamen | 0.810 | -0.054 | 0.020 | -2.488 | 0.013 | -0.091 | -0.013 |
| auditory : left-accumbens | 0.809 | 0.054 | 0.018 | 2.856 | 0.004 | 0.015 | 0.085 |
| cingulo-opercular : right-accumbens | 0.807 | 0.053 | 0.020 | 2.550 | 0.011 | 0.013 | 0.090 |
| ventral-attention : right-accumbens | 0.806 | -0.053 | 0.020 | -2.581 | 0.010 | -0.090 | -0.012 |
| cingulo-opercular : left-thalamus | 0.800 | 0.053 | 0.019 | 2.632 | 0.009 | 0.013 | 0.088 |
| default : left-cerebellum | 0.777 | -0.051 | 0.020 | -2.455 | 0.014 | -0.087 | -0.010 |
| default : right-amygdala | 0.777 | 0.051 | 0.019 | 2.607 | 0.009 | 0.011 | 0.086 |
| cingulo-parietal : left-amygdala | 0.777 | 0.051 | 0.021 | 2.409 | 0.016 | 0.012 | 0.090 |
| retrosplenial temporal : left-amygdala | 0.776 | -0.051 | 0.019 | -2.639 | 0.008 | -0.084 | -0.012 |
| cingulo-opercular : left-cerebellum | 0.754 | 0.050 | 0.021 | 2.307 | 0.021 | 0.010 | 0.090 |
| auditory : left-hippocampus | 0.752 | -0.050 | 0.020 | -2.351 | 0.019 | -0.088 | -0.007 |
| cingulo-parietal : right-ventraldc | 0.752 | -0.050 | 0.020 | -2.387 | 0.017 | -0.087 | -0.010 |
| retrosplenial temporal : right-pallidum | 0.735 | 0.049 | 0.020 | 2.337 | 0.020 | 0.008 | 0.084 |
| fronto-parietal : left-accumbens | 0.733 | -0.049 | 0.020 | -2.408 | 0.016 | -0.082 | -0.008 |

| | | | | | | | |
|---|---|---|---|---|---|---|---|
| dorsal attention : right-accumbens | 0.720 | 0.048 | 0.020 | 2.236 | 0.025 | 0.006 | 0.086 |
| retrosplenial temporal : left-caudate | 0.700 | -0.046 | 0.019 | -2.395 | 0.017 | -0.083 | -0.009 |
| default : right-pallidum | 0.686 | 0.045 | 0.020 | 2.208 | 0.027 | 0.007 | 0.082 |
| salience : right-thalamus | 0.683 | 0.045 | 0.019 | 2.255 | 0.024 | 0.005 | 0.081 |
| ventral-attention : left-amygdala | 0.679 | -0.045 | 0.020 | -2.219 | 0.027 | -0.081 | -0.004 |
| auditory : right-cerebellum | 0.676 | -0.045 | 0.020 | -2.143 | 0.032 | -0.079 | -0.003 |
| fronto-parietal : right-caudate | 0.671 | 0.044 | 0.019 | 2.190 | 0.029 | 0.002 | 0.080 |
| visual : left-caudate | 0.670 | 0.044 | 0.019 | 2.271 | 0.023 | 0.004 | 0.081 |
| sensorimotor mouth : left-pallidum | 0.653 | 0.043 | 0.021 | 1.978 | 0.048 | 0.003 | 0.082 |
| fronto-parietal : right-amygdala | 0.650 | 0.043 | 0.020 | 1.973 | 0.049 | 0.000 | 0.078 |
| default : left-caudate | 0.624 | -0.041 | 0.020 | -1.945 | 0.052 | -0.077 | 0.001 |
| ventral-attention : left-thalamus | 0.603 | 0.040 | 0.019 | 2.056 | 0.040 | 0.001 | 0.073 |
| sensorimotor mouth : left-accumbens | 0.592 | 0.039 | 0.020 | 1.898 | 0.058 | 0.000 | 0.078 |
| dorsal attention : left-thalamus | 0.583 | 0.039 | 0.021 | 1.764 | 0.078 | -0.003 | 0.077 |
| sensorimotor hand : left-putamen | 0.581 | 0.038 | 0.020 | 1.809 | 0.071 | -0.003 | 0.076 |
| visual : right-putamen | 0.568 | 0.038 | 0.019 | 1.844 | 0.065 | -0.002 | 0.073 |
| fronto-parietal : left-thalamus | 0.544 | -0.036 | 0.020 | -1.650 | 0.099 | -0.072 | 0.005 |
| cingulo-parietal : left-hippocampus | 0.544 | -0.036 | 0.020 | -1.752 | 0.080 | -0.076 | 0.004 |
| visual : right-ventraldc | 0.537 | 0.036 | 0.019 | 1.765 | 0.078 | -0.003 | 0.070 |
| sensorimotor hand : right-pallidum | 0.532 | 0.035 | 0.020 | 1.714 | 0.087 | -0.004 | 0.071 |
| visual : right-caudate | 0.528 | 0.035 | 0.019 | 1.741 | 0.082 | -0.004 | 0.074 |
| sensorimotor mouth : left-putamen | 0.512 | -0.034 | 0.020 | -1.651 | 0.099 | -0.070 | 0.006 |
| ventral-attention : right-putamen | 0.497 | 0.033 | 0.021 | 1.496 | 0.135 | -0.008 | 0.073 |
| ventral-attention : right-cerebellum | 0.489 | 0.032 | 0.020 | 1.520 | 0.128 | -0.010 | 0.070 |
| sensorimotor mouth : right-pallidum | 0.479 | -0.032 | 0.021 | -1.500 | 0.134 | -0.070 | 0.010 |
| salience : left-amygdala | 0.478 | 0.032 | 0.021 | 1.404 | 0.160 | -0.010 | 0.068 |
| ventral-attention : right-thalamus | 0.460 | -0.030 | 0.019 | -1.531 | 0.126 | -0.064 | 0.009 |
| auditory : left-thalamus | 0.459 | 0.030 | 0.021 | 1.469 | 0.142 | -0.009 | 0.070 |
| salience : right-caudate | 0.446 | 0.030 | 0.020 | 1.327 | 0.185 | -0.014 | 0.067 |

| | | | | | | | |
|---|---|---|---|---|---|---|---|
| fronto-parietal : left-amygdala | 0.439 | 0.029 | 0.020 | 1.460 | 0.144 | -0.008 | 0.069 |
| salience : right-cerebellum | 0.429 | -0.028 | 0.020 | -1.294 | 0.196 | -0.067 | 0.014 |
| retrosplenial temporal : right-putamen | 0.427 | -0.028 | 0.020 | -1.367 | 0.172 | -0.063 | 0.013 |
| fronto-parietal : brain-stem | 0.398 | 0.026 | 0.020 | 1.301 | 0.193 | -0.012 | 0.067 |
| sensorimotor mouth : right-ventraldc | 0.382 | -0.025 | 0.019 | -1.252 | 0.211 | -0.062 | 0.011 |
| dorsal attention : right-caudate | 0.381 | 0.025 | 0.021 | 1.146 | 0.252 | -0.015 | 0.064 |
| salience : right-hippocampus | 0.380 | 0.025 | 0.021 | 1.152 | 0.249 | -0.019 | 0.065 |
| salience : right-putamen | 0.378 | -0.025 | 0.021 | -1.165 | 0.244 | -0.064 | 0.019 |
| dorsal attention : left-caudate | 0.366 | 0.024 | 0.021 | 1.133 | 0.257 | -0.014 | 0.065 |
| sensorimotor mouth : right-amygdala | 0.364 | -0.024 | 0.019 | -1.174 | 0.240 | -0.060 | 0.015 |
| fronto-parietal : right-ventraldc | 0.360 | 0.024 | 0.020 | 1.185 | 0.236 | -0.016 | 0.062 |
| dorsal attention : left-cerebellum | 0.360 | 0.024 | 0.020 | 1.194 | 0.232 | -0.016 | 0.061 |
| sensorimotor hand : left-accumbens | 0.359 | 0.024 | 0.021 | 1.126 | 0.260 | -0.015 | 0.065 |
| visual : brain-stem | 0.357 | -0.024 | 0.020 | -1.113 | 0.266 | -0.061 | 0.016 |
| ventral-attention : left-cerebellum | 0.352 | -0.023 | 0.020 | -1.069 | 0.285 | -0.063 | 0.016 |
| visual : left-accumbens | 0.338 | -0.022 | 0.019 | -1.158 | 0.247 | -0.060 | 0.016 |
| dorsal attention : right-pallidum | 0.331 | -0.022 | 0.021 | -1.044 | 0.297 | -0.063 | 0.018 |
| default : right-caudate | 0.324 | 0.021 | 0.020 | 1.090 | 0.276 | -0.016 | 0.062 |
| auditory : right-caudate | 0.322 | 0.021 | 0.021 | 0.981 | 0.327 | -0.022 | 0.059 |
| visual : left-ventraldc | 0.316 | -0.021 | 0.019 | -1.040 | 0.298 | -0.057 | 0.018 |
| dorsal attention : right-hippocampus | 0.315 | 0.021 | 0.020 | 1.009 | 0.313 | -0.019 | 0.060 |
| visual : right-cerebellum | 0.312 | 0.021 | 0.020 | 0.979 | 0.327 | -0.019 | 0.060 |
| dorsal attention : right-putamen | 0.311 | 0.021 | 0.019 | 1.075 | 0.283 | -0.016 | 0.058 |
| default : left-thalamus | 0.309 | 0.020 | 0.021 | 0.915 | 0.360 | -0.021 | 0.061 |
| auditory : right-accumbens | 0.308 | 0.020 | 0.020 | 0.993 | 0.321 | -0.020 | 0.060 |
| sensorimotor mouth : right-putamen | 0.302 | -0.020 | 0.020 | -0.969 | 0.333 | -0.061 | 0.018 |
| retrosplenial temporal : left-putamen | 0.302 | -0.020 | 0.020 | -1.017 | 0.309 | -0.055 | 0.018 |
| default : right-thalamus | 0.300 | -0.020 | 0.019 | -0.979 | 0.328 | -0.058 | 0.018 |
| default : brain-stem | 0.289 | 0.019 | 0.020 | 0.905 | 0.366 | -0.019 | 0.054 |

| | | | | | | | |
|---|---|---|---|---|---|---|---|
| dorsal attention : left-putamen | 0.289 | -0.019 | 0.020 | -0.874 | 0.382 | -0.058 | 0.023 |
| cingulo-parietal : left-accumbens | 0.272 | 0.018 | 0.020 | 0.913 | 0.362 | -0.020 | 0.057 |
| salience : left-hippocampus | 0.271 | -0.018 | 0.021 | -0.790 | 0.429 | -0.057 | 0.025 |
| salience : right-accumbens | 0.271 | -0.018 | 0.020 | -0.892 | 0.372 | -0.057 | 0.021 |
| sensorimotor mouth : brain-stem | 0.266 | 0.018 | 0.019 | 0.892 | 0.373 | -0.020 | 0.055 |
| cingulo-parietal : left-pallidum | 0.264 | 0.018 | 0.021 | 0.819 | 0.413 | -0.024 | 0.057 |
| sensorimotor hand : right-cerebellum | 0.257 | 0.017 | 0.021 | 0.770 | 0.441 | -0.024 | 0.057 |
| fronto-parietal : right-cerebellum | 0.249 | 0.016 | 0.021 | 0.803 | 0.422 | -0.025 | 0.057 |
| fronto-parietal : right-thalamus | 0.241 | -0.016 | 0.020 | -0.821 | 0.412 | -0.056 | 0.022 |
| visual : left-cerebellum | 0.231 | 0.015 | 0.020 | 0.738 | 0.460 | -0.025 | 0.054 |
| dorsal attention : right-thalamus | 0.231 | -0.015 | 0.020 | -0.717 | 0.473 | -0.054 | 0.022 |
| retrosplenial temporal : right-caudate | 0.229 | -0.015 | 0.019 | -0.837 | 0.402 | -0.051 | 0.021 |
| sensorimotor mouth : left-thalamus | 0.223 | 0.015 | 0.021 | 0.666 | 0.506 | -0.028 | 0.053 |
| cingulo-opercular : left-hippocampus | 0.211 | 0.014 | 0.021 | 0.638 | 0.523 | -0.027 | 0.055 |
| cingulo-opercular : left-pallidum | 0.207 | 0.014 | 0.020 | 0.661 | 0.509 | -0.024 | 0.054 |
| ventral-attention : right-pallidum | 0.207 | 0.014 | 0.021 | 0.574 | 0.566 | -0.026 | 0.055 |
| sensorimotor mouth : right-accumbens | 0.197 | -0.013 | 0.021 | -0.607 | 0.544 | -0.055 | 0.027 |
| cingulo-parietal : left-putamen | 0.196 | -0.013 | 0.020 | -0.685 | 0.493 | -0.055 | 0.025 |
| sensorimotor hand : right-amygdala | 0.190 | 0.013 | 0.021 | 0.614 | 0.539 | -0.027 | 0.052 |
| ventral-attention : right-amygdala | 0.189 | -0.013 | 0.019 | -0.573 | 0.566 | -0.049 | 0.027 |
| sensorimotor hand : right-thalamus | 0.185 | -0.012 | 0.019 | -0.626 | 0.531 | -0.052 | 0.028 |
| fronto-parietal : left-hippocampus | 0.166 | 0.011 | 0.020 | 0.572 | 0.567 | -0.027 | 0.050 |
| default : right-cerebellum | 0.163 | -0.011 | 0.019 | -0.576 | 0.564 | -0.047 | 0.027 |
| retrosplenial temporal : right-amygdala | 0.158 | 0.010 | 0.021 | 0.484 | 0.629 | -0.031 | 0.053 |
| ventral-attention : left-ventraldc | 0.154 | 0.010 | 0.020 | 0.543 | 0.587 | -0.026 | 0.051 |
| salience : right-ventraldc | 0.151 | -0.010 | 0.020 | -0.511 | 0.610 | -0.047 | 0.026 |
| cingulo-opercular : right-cerebellum | 0.151 | -0.010 | 0.020 | -0.492 | 0.623 | -0.049 | 0.032 |
| dorsal attention : right-cerebellum | 0.148 | -0.010 | 0.021 | -0.425 | 0.671 | -0.049 | 0.033 |
| default : left-amygdala | 0.145 | -0.010 | 0.020 | -0.459 | 0.646 | -0.051 | 0.027 |

| | | | | | | | |
|---|---|---|---|---|---|---|---|
| visual : left-putamen | 0.145 | 0.010 | 0.020 | 0.477 | 0.633 | -0.028 | 0.048 |
| ventral-attention : right-hippocampus | 0.138 | -0.009 | 0.020 | -0.438 | 0.662 | -0.048 | 0.029 |
| cingulo-parietal : brain-stem | 0.135 | -0.009 | 0.020 | -0.422 | 0.673 | -0.046 | 0.031 |
| sensorimotor mouth : right-cerebellum | 0.132 | -0.009 | 0.021 | -0.372 | 0.710 | -0.046 | 0.034 |
| cingulo-parietal : right-putamen | 0.123 | -0.008 | 0.020 | -0.359 | 0.720 | -0.045 | 0.035 |
| visual : right-amygdala | 0.122 | -0.008 | 0.021 | -0.367 | 0.714 | -0.048 | 0.034 |
| salience : right-pallidum | 0.121 | -0.008 | 0.020 | -0.383 | 0.701 | -0.043 | 0.034 |
| cingulo-opercular : right-pallidum | 0.117 | 0.008 | 0.020 | 0.386 | 0.699 | -0.030 | 0.047 |
| fronto-parietal : left-pallidum | 0.104 | -0.007 | 0.020 | -0.351 | 0.725 | -0.049 | 0.033 |
| auditory : right-amygdala | 0.094 | 0.006 | 0.020 | 0.270 | 0.787 | -0.033 | 0.047 |
| visual : left-pallidum | 0.088 | -0.006 | 0.020 | -0.261 | 0.794 | -0.046 | 0.033 |
| sensorimotor mouth : right-hippocampus | 0.082 | 0.005 | 0.021 | 0.265 | 0.791 | -0.034 | 0.047 |
| sensorimotor mouth : left-caudate | 0.081 | -0.005 | 0.021 | -0.249 | 0.803 | -0.045 | 0.034 |
| salience : left-caudate | 0.076 | 0.005 | 0.020 | 0.267 | 0.789 | -0.034 | 0.046 |
| retrosplenial temporal : left-thalamus | 0.075 | 0.005 | 0.021 | 0.283 | 0.777 | -0.036 | 0.046 |
| cingulo-opercular : right-thalamus | 0.074 | 0.005 | 0.020 | 0.240 | 0.810 | -0.034 | 0.045 |
| cingulo-parietal : left-cerebellum | 0.071 | 0.005 | 0.021 | 0.243 | 0.808 | -0.036 | 0.047 |
| auditory : left-caudate | 0.067 | 0.004 | 0.020 | 0.200 | 0.841 | -0.033 | 0.043 |
| auditory : right-thalamus | 0.066 | 0.004 | 0.021 | 0.216 | 0.829 | -0.037 | 0.045 |
| salience : right-amygdala | 0.066 | -0.004 | 0.020 | -0.169 | 0.866 | -0.042 | 0.034 |
| dorsal attention : left-pallidum | 0.065 | -0.004 | 0.020 | -0.266 | 0.791 | -0.044 | 0.034 |
| retrosplenial temporal : left-ventraldc | 0.062 | -0.004 | 0.021 | -0.160 | 0.873 | -0.044 | 0.038 |
| dorsal attention : left-ventraldc | 0.050 | 0.003 | 0.020 | 0.158 | 0.874 | -0.037 | 0.038 |
| salience : left-pallidum | 0.037 | -0.002 | 0.021 | -0.072 | 0.943 | -0.042 | 0.040 |
| auditory : left-pallidum | 0.021 | 0.001 | 0.022 | 0.096 | 0.924 | -0.042 | 0.044 |
| retrosplenial temporal : left-accumbens | 0.020 | -0.001 | 0.020 | -0.054 | 0.957 | -0.040 | 0.039 |
| dorsal attention : right-amygdala | 0.016 | -0.001 | 0.020 | -0.011 | 0.991 | -0.038 | 0.039 |
| sensorimotor mouth : left-cerebellum | 0.014 | -0.001 | 0.021 | -0.071 | 0.943 | -0.041 | 0.039 |
| salience : brain-stem | 0.011 | 0.001 | 0.021 | 0.026 | 0.979 | -0.038 | 0.042 |

| | | | | | | | |
|---|---|---|---|---|---|---|---|
| retrosplenial temporal : left-pallidum | 0.010 | -0.001 | 0.020 | -0.022 | 0.983 | -0.039 | 0.039 |
| cingulo-parietal : right-accumbens | 0.009 | 0.001 | 0.020 | 0.052 | 0.959 | -0.037 | 0.041 |
| auditory : left-amygdala | 0.007 | 0.000 | 0.020 | -0.061 | 0.952 | -0.041 | 0.036 |
| default : right-ventraldc | 0.006 | 0.000 | 0.020 | -0.006 | 0.995 | -0.037 | 0.039 |
| sensorimotor hand : left-thalamus | 0.003 | 0.000 | 0.020 | 0.023 | 0.982 | -0.038 | 0.040 |
| ventral-attention : left-accumbens | 0.000 | 0.000 | 0.019 | 0.012 | 0.991 | -0.034 | 0.041 |

*Notes*. Variable importance in projection (VIP) was used to assess the relative importance of each item in the PLS model. Bold indicates VIP scores considered most influential in terms of their explanatory power (>1). Bootstrapping with 1,000 resampling iterations was used to obtain standard errors, t-values confidence intervals, and p-values.

**Table S30.** PLS: cortico-limbic regional connectivity VIP scores and loadings for the PRS-only model

| | VIP | Loading | Std. err | t | p | 2.50% | 97.50% |
|---|---|---|---|---|---|---|---|
| *Predictor (PRS)* | | | | | | | |
| **prsDepression** | **1.391** | **-0.696** | **0.051** | **-13.477** | **0.000** | **-0.782** | **-0.581** |
| **prsADHD** | **1.255** | **-0.628** | **0.059** | **-10.365** | **0.000** | **-0.726** | **-0.499** |
| prsAnxiety | 0.694 | -0.347 | 0.072 | -4.989 | 0.000 | -0.493 | -0.203 |
| prsPsychosis | 0.082 | -0.041 | 0.094 | -0.675 | 0.500 | -0.235 | 0.139 |
| *Outcome (regional connectivity)* | | | | | | | |
| **auditory : right-putamen** | **2.339** | **0.155** | **0.015** | **9.799** | **0.000** | **0.117** | **0.174** |
| **sensorimotor hand : left-pallidum** | **2.333** | **0.154** | **0.015** | **9.406** | **0.000** | **0.113** | **0.174** |
| **cingulo-opercular : left-amygdala** | **2.327** | **0.154** | **0.016** | **9.307** | **0.000** | **0.115** | **0.173** |
| **visual : left-hippocampus** | **2.258** | **0.150** | **0.019** | **7.499** | **0.000** | **0.103** | **0.178** |
| **cingulo-parietal : left-thalamus** | **2.217** | **0.147** | **0.016** | **8.488** | **0.000** | **0.105** | **0.169** |
| **sensorimotor hand : brain-stem** | **2.205** | **0.146** | **0.019** | **7.244** | **0.000** | **0.101** | **0.175** |
| **sensorimotor hand : right-accumbens** | **2.165** | **0.143** | **0.016** | **8.174** | **0.000** | **0.100** | **0.164** |

| | | | | | | | |
|---|---|---|---|---|---|---|---|
| **retrosplenial temporal : right-cerebellum** | **2.069** | **0.137** | **0.016** | **8.194** | **0.000** | **0.099** | **0.158** |
| **sensorimotor hand : right-caudate** | **2.059** | **0.136** | **0.017** | **7.545** | **0.000** | **0.094** | **0.160** |
| **salience : left-thalamus** | **2.010** | **0.133** | **0.018** | **6.777** | **0.000** | **0.088** | **0.160** |
| **sensorimotor hand : right-putamen** | **1.995** | **0.132** | **0.019** | **6.466** | **0.000** | **0.087** | **0.162** |
| **cingulo-opercular : brain-stem** | **1.982** | **0.131** | **0.020** | **6.073** | **0.000** | **0.085** | **0.165** |
| **cingulo-opercular : right-amygdala** | **1.970** | **0.130** | **0.020** | **6.263** | **0.000** | **0.085** | **0.160** |
| **retrosplenial temporal : right-thalamus** | **1.924** | **0.127** | **0.021** | **5.804** | **0.000** | **0.080** | **0.160** |
| **cingulo-opercular : left-putamen** | **1.918** | **0.127** | **0.017** | **6.848** | **0.000** | **0.085** | **0.151** |
| **sensorimotor mouth : right-thalamus** | **1.857** | **0.123** | **0.018** | **6.456** | **0.000** | **0.080** | **0.151** |
| **auditory : right-pallidum** | **1.835** | **0.122** | **0.020** | **5.823** | **0.000** | **0.076** | **0.152** |
| **cingulo-opercular : left-accumbens** | **1.828** | **0.121** | **0.020** | **5.837** | **0.000** | **0.075** | **0.152** |
| **sensorimotor mouth : right-caudate** | **1.821** | **0.121** | **0.021** | **5.331** | **0.000** | **0.069** | **0.154** |
| **cingulo-opercular : right-hippocampus** | **1.810** | **0.120** | **0.018** | **6.447** | **0.000** | **0.078** | **0.147** |
| **visual : right-hippocampus** | **1.799** | **0.119** | **0.019** | **5.774** | **0.000** | **0.072** | **0.148** |
| **cingulo-parietal : left-caudate** | **1.767** | **0.117** | **0.020** | **5.317** | **0.000** | **0.069** | **0.146** |
| **sensorimotor hand : right-ventraldc** | **1.729** | **0.115** | **0.021** | **5.132** | **0.000** | **0.064** | **0.145** |
| **dorsal attention : left-hippocampus** | **1.724** | **-0.114** | **0.023** | **-4.633** | **0.000** | **-0.152** | **-0.063** |
| **cingulo-opercular : left-caudate** | **1.672** | **0.111** | **0.018** | **5.863** | **0.000** | **0.070** | **0.137** |
| **cingulo-opercular : right-ventraldc** | **1.663** | **0.110** | **0.020** | **5.213** | **0.000** | **0.065** | **0.142** |
| **cingulo-opercular : left-ventraldc** | **1.637** | **-0.108** | **0.023** | **-4.426** | **0.000** | **-0.150** | **-0.059** |
| **retrosplenial temporal : right-ventraldc** | **1.623** | **0.108** | **0.019** | **5.354** | **0.000** | **0.064** | **0.138** |
| **cingulo-parietal : right-pallidum** | **1.612** | **0.107** | **0.018** | **5.506** | **0.000** | **0.063** | **0.136** |
| **auditory : left-ventraldc** | **1.600** | **0.106** | **0.022** | **4.618** | **0.000** | **0.058** | **0.145** |
| **sensorimotor hand : left-cerebellum** | **1.592** | **0.105** | **0.018** | **5.555** | **0.000** | **0.063** | **0.133** |
| **default : left-hippocampus** | **1.566** | **0.104** | **0.023** | **4.290** | **0.000** | **0.054** | **0.146** |
| **default : left-pallidum** | **1.566** | **0.104** | **0.020** | **4.838** | **0.000** | **0.061** | **0.136** |
| **dorsal attention : right-ventraldc** | **1.547** | **0.102** | **0.022** | **4.377** | **0.000** | **0.052** | **0.141** |
| **visual : left-amygdala** | **1.545** | **0.102** | **0.023** | **4.244** | **0.000** | **0.052** | **0.141** |
| **ventral-attention : left-hippocampus** | **1.540** | **0.102** | **0.023** | **4.200** | **0.000** | **0.052** | **0.139** |

| | | | | | | | |
|---|---|---|---|---|---|---|---|
| **cingulo-parietal : right-caudate** | **1.529** | **0.101** | **0.020** | **4.612** | **0.000** | **0.055** | **0.134** |
| **cingulo-parietal : left-ventraldc** | **1.526** | **0.101** | **0.020** | **4.717** | **0.000** | **0.055** | **0.132** |
| **ventral-attention : left-caudate** | **1.495** | **0.099** | **0.022** | **4.208** | **0.000** | **0.050** | **0.137** |
| **cingulo-parietal : right-hippocampus** | **1.477** | **0.098** | **0.021** | **4.296** | **0.000** | **0.048** | **0.131** |
| **visual : right-pallidum** | **1.473** | **0.098** | **0.021** | **4.503** | **0.000** | **0.053** | **0.131** |
| **default : left-accumbens** | **1.473** | **0.098** | **0.019** | **4.755** | **0.000** | **0.053** | **0.129** |
| **cingulo-parietal : right-cerebellum** | **1.472** | **0.097** | **0.019** | **4.712** | **0.000** | **0.050** | **0.126** |
| **fronto-parietal : left-caudate** | **1.407** | **0.093** | **0.021** | **4.148** | **0.000** | **0.045** | **0.128** |
| **sensorimotor mouth : left-amygdala** | **1.392** | **0.092** | **0.020** | **4.369** | **0.000** | **0.046** | **0.123** |
| **sensorimotor mouth : left-ventraldc** | **1.388** | **0.092** | **0.021** | **4.125** | **0.000** | **0.046** | **0.126** |
| **sensorimotor mouth : left-hippocampus** | **1.381** | **0.091** | **0.021** | **4.030** | **0.000** | **0.044** | **0.124** |
| **ventral-attention : brain-stem** | **1.337** | **0.089** | **0.022** | **3.814** | **0.000** | **0.042** | **0.126** |
| **auditory : left-cerebellum** | **1.326** | **0.088** | **0.023** | **3.673** | **0.000** | **0.039** | **0.131** |
| **salience : left-ventraldc** | **1.320** | **-0.087** | **0.023** | **-3.463** | **0.001** | **-0.125** | **-0.037** |
| **auditory : right-hippocampus** | **1.299** | **-0.086** | **0.022** | **-3.699** | **0.000** | **-0.127** | **-0.041** |
| **retrosplenial temporal : left-hippocampus** | **1.288** | **0.085** | **0.023** | **3.587** | **0.000** | **0.037** | **0.124** |
| **sensorimotor hand : left-ventraldc** | **1.281** | **-0.085** | **0.024** | **-3.392** | **0.001** | **-0.124** | **-0.037** |
| **fronto-parietal : right-hippocampus** | **1.264** | **0.084** | **0.023** | **3.363** | **0.001** | **0.033** | **0.122** |
| **dorsal attention : brain-stem** | **1.247** | **0.083** | **0.022** | **3.430** | **0.001** | **0.030** | **0.120** |
| **visual : left-thalamus** | **1.234** | **0.082** | **0.023** | **3.322** | **0.001** | **0.030** | **0.126** |
| **dorsal attention : left-accumbens** | **1.219** | **0.081** | **0.022** | **3.419** | **0.001** | **0.033** | **0.120** |
| **default : right-accumbens** | **1.217** | **0.081** | **0.021** | **3.629** | **0.000** | **0.036** | **0.115** |
| **visual : right-accumbens** | **1.216** | **0.081** | **0.022** | **3.458** | **0.001** | **0.032** | **0.121** |
| **visual : right-thalamus** | **1.213** | **-0.080** | **0.023** | **-3.285** | **0.001** | **-0.119** | **-0.028** |
| **sensorimotor hand : left-caudate** | **1.153** | **0.076** | **0.022** | **3.340** | **0.001** | **0.032** | **0.115** |
| **cingulo-parietal : right-thalamus** | **1.134** | **0.075** | **0.023** | **2.998** | **0.003** | **0.025** | **0.115** |
| **default : left-putamen** | **1.060** | **0.070** | **0.021** | **3.120** | **0.002** | **0.023** | **0.105** |
| **ventral-attention : left-pallidum** | **1.056** | **-0.070** | **0.024** | **-2.671** | **0.008** | **-0.114** | **-0.018** |
| **cingulo-opercular : right-caudate** | **1.055** | **-0.070** | **0.024** | **-2.728** | **0.006** | **-0.114** | **-0.019** |

| | | | | | | | |
|---|---|---|---|---|---|---|---|
| **fronto-parietal : right-putamen** | **1.054** | **-0.070** | **0.023** | **-2.742** | **0.006** | **-0.111** | **-0.018** |
| **auditory : right-ventraldc** | **1.041** | **-0.069** | **0.022** | **-2.907** | **0.004** | **-0.105** | **-0.020** |
| **fronto-parietal : right-accumbens** | **1.032** | **0.068** | **0.023** | **2.868** | **0.004** | **0.018** | **0.111** |
| **salience : left-putamen** | **1.030** | **0.068** | **0.022** | **2.935** | **0.003** | **0.019** | **0.109** |
| **auditory : brain-stem** | **1.021** | **-0.068** | **0.022** | **-2.840** | **0.005** | **-0.106** | **-0.018** |
| **fronto-parietal : left-ventraldc** | **1.012** | **0.067** | **0.022** | **2.865** | **0.004** | **0.020** | **0.110** |
| **ventral-attention : right-caudate** | **1.004** | **0.066** | **0.022** | **2.815** | **0.005** | **0.018** | **0.106** |
| **fronto-parietal : left-cerebellum** | **1.003** | **-0.066** | **0.023** | **-2.714** | **0.007** | **-0.110** | **-0.017** |
| default : right-putamen | 0.998 | -0.066 | 0.024 | -2.486 | 0.013 | -0.109 | -0.016 |
| fronto-parietal : left-putamen | 0.989 | 0.065 | 0.023 | 2.690 | 0.007 | 0.018 | 0.104 |
| auditory : left-putamen | 0.979 | -0.065 | 0.023 | -2.657 | 0.008 | -0.104 | -0.017 |
| salience : left-accumbens | 0.978 | 0.065 | 0.022 | 2.746 | 0.006 | 0.015 | 0.100 |
| cingulo-parietal : right-amygdala | 0.968 | -0.064 | 0.025 | -2.395 | 0.017 | -0.110 | -0.011 |
| cingulo-opercular : right-accumbens | 0.944 | 0.063 | 0.023 | 2.518 | 0.012 | 0.017 | 0.104 |
| dorsal attention : left-amygdala | 0.944 | -0.062 | 0.025 | -2.339 | 0.019 | -0.110 | -0.013 |
| retrosplenial temporal : right-hippocampus | 0.925 | -0.061 | 0.022 | -2.555 | 0.011 | -0.099 | -0.012 |
| ventral-attention : left-putamen | 0.924 | -0.061 | 0.023 | -2.487 | 0.013 | -0.101 | -0.011 |
| default : right-hippocampus | 0.923 | 0.061 | 0.024 | 2.349 | 0.019 | 0.011 | 0.103 |
| fronto-parietal : right-pallidum | 0.920 | -0.061 | 0.023 | -2.517 | 0.012 | -0.104 | -0.014 |
| cingulo-opercular : right-putamen | 0.912 | 0.060 | 0.023 | 2.424 | 0.015 | 0.012 | 0.103 |
| default : left-ventraldc | 0.893 | 0.059 | 0.023 | 2.509 | 0.012 | 0.013 | 0.103 |
| retrosplenial temporal : brain-stem | 0.892 | 0.059 | 0.022 | 2.567 | 0.010 | 0.014 | 0.097 |
| salience : right-thalamus | 0.890 | 0.059 | 0.022 | 2.466 | 0.014 | 0.008 | 0.097 |
| retrosplenial temporal : right-accumbens | 0.883 | 0.058 | 0.023 | 2.314 | 0.021 | 0.010 | 0.101 |
| cingulo-parietal : left-amygdala | 0.858 | 0.057 | 0.023 | 2.289 | 0.022 | 0.010 | 0.102 |
| default : right-pallidum | 0.852 | 0.056 | 0.023 | 2.303 | 0.021 | 0.008 | 0.099 |
| cingulo-parietal : right-ventraldc | 0.838 | -0.056 | 0.023 | -2.275 | 0.023 | -0.100 | -0.010 |
| fronto-parietal : right-caudate | 0.835 | 0.055 | 0.023 | 2.256 | 0.024 | 0.005 | 0.095 |
| fronto-parietal : left-accumbens | 0.832 | -0.055 | 0.024 | -2.207 | 0.027 | -0.099 | -0.005 |

| | | | | | | | |
|---|---|---|---|---|---|---|---|
| cingulo-opercular : left-cerebellum | 0.823 | 0.055 | 0.024 | 2.119 | 0.034 | 0.005 | 0.100 |
| sensorimotor mouth : left-accumbens | 0.818 | 0.054 | 0.023 | 2.199 | 0.028 | 0.003 | 0.093 |
| ventral-attention : right-ventraldc | 0.817 | 0.054 | 0.021 | 2.404 | 0.016 | 0.009 | 0.092 |
| sensorimotor hand : left-amygdala | 0.804 | -0.053 | 0.024 | -2.074 | 0.038 | -0.097 | -0.003 |
| ventral-attention : right-putamen | 0.779 | 0.052 | 0.024 | 1.971 | 0.049 | 0.001 | 0.096 |
| sensorimotor hand : right-hippocampus | 0.774 | 0.051 | 0.021 | 2.338 | 0.019 | 0.006 | 0.088 |
| cingulo-parietal : left-hippocampus | 0.771 | -0.051 | 0.024 | -2.007 | 0.045 | -0.097 | 0.001 |
| ventral-attention : right-accumbens | 0.766 | -0.051 | 0.023 | -2.034 | 0.042 | -0.093 | -0.001 |
| sensorimotor hand : left-hippocampus | 0.754 | -0.050 | 0.023 | -2.079 | 0.038 | -0.093 | -0.004 |
| visual : right-caudate | 0.732 | 0.048 | 0.023 | 2.025 | 0.043 | 0.001 | 0.089 |
| sensorimotor mouth : left-pallidum | 0.725 | 0.048 | 0.024 | 1.879 | 0.060 | -0.001 | 0.092 |
| retrosplenial temporal : left-amygdala | 0.699 | -0.046 | 0.022 | -1.892 | 0.059 | -0.085 | 0.003 |
| fronto-parietal : left-thalamus | 0.692 | -0.046 | 0.024 | -1.744 | 0.081 | -0.087 | 0.006 |
| salience : left-cerebellum | 0.682 | 0.045 | 0.023 | 1.897 | 0.058 | -0.001 | 0.085 |
| default : right-amygdala | 0.674 | 0.045 | 0.023 | 1.900 | 0.058 | -0.003 | 0.086 |
| fronto-parietal : left-amygdala | 0.669 | 0.044 | 0.023 | 1.854 | 0.064 | -0.002 | 0.086 |
| sensorimotor mouth : left-putamen | 0.662 | -0.044 | 0.024 | -1.773 | 0.076 | -0.086 | 0.007 |
| default : brain-stem | 0.655 | 0.043 | 0.024 | 1.681 | 0.093 | -0.006 | 0.084 |
| auditory : left-hippocampus | 0.654 | -0.043 | 0.024 | -1.681 | 0.093 | -0.086 | 0.005 |
| retrosplenial temporal : left-cerebellum | 0.634 | -0.042 | 0.023 | -1.759 | 0.079 | -0.086 | 0.005 |
| auditory : right-cerebellum | 0.631 | -0.042 | 0.024 | -1.579 | 0.114 | -0.080 | 0.008 |
| cingulo-opercular : left-thalamus | 0.627 | 0.041 | 0.023 | 1.687 | 0.092 | -0.008 | 0.082 |
| auditory : left-accumbens | 0.607 | 0.040 | 0.021 | 1.848 | 0.065 | -0.004 | 0.078 |
| visual : left-caudate | 0.606 | 0.040 | 0.023 | 1.689 | 0.091 | -0.005 | 0.081 |
| dorsal attention : left-thalamus | 0.599 | 0.040 | 0.025 | 1.437 | 0.151 | -0.011 | 0.087 |
| dorsal attention : right-caudate | 0.588 | 0.039 | 0.023 | 1.546 | 0.122 | -0.009 | 0.080 |
| salience : right-hippocampus | 0.588 | 0.039 | 0.024 | 1.520 | 0.129 | -0.012 | 0.079 |
| ventral-attention : left-amygdala | 0.568 | -0.038 | 0.023 | -1.551 | 0.121 | -0.082 | 0.010 |
| retrosplenial temporal : left-caudate | 0.564 | -0.037 | 0.023 | -1.593 | 0.111 | -0.078 | 0.011 |

| | | | | | | | |
|---|---|---|---|---|---|---|---|
| visual : right-putamen | 0.563 | 0.037 | 0.024 | 1.454 | 0.146 | -0.009 | 0.080 |
| default : left-caudate | 0.560 | -0.037 | 0.023 | -1.503 | 0.133 | -0.079 | 0.011 |
| retrosplenial temporal : right-pallidum | 0.554 | 0.037 | 0.023 | 1.511 | 0.131 | -0.011 | 0.079 |
| salience : right-putamen | 0.536 | -0.036 | 0.024 | -1.355 | 0.175 | -0.076 | 0.015 |
| cingulo-parietal : left-pallidum | 0.523 | 0.035 | 0.024 | 1.366 | 0.172 | -0.016 | 0.078 |
| visual : left-accumbens | 0.518 | -0.034 | 0.023 | -1.433 | 0.152 | -0.078 | 0.010 |
| default : left-cerebellum | 0.518 | -0.034 | 0.023 | -1.382 | 0.167 | -0.077 | 0.013 |
| visual : right-ventraldc | 0.515 | 0.034 | 0.023 | 1.380 | 0.168 | -0.014 | 0.076 |
| sensorimotor hand : right-pallidum | 0.508 | 0.034 | 0.023 | 1.379 | 0.168 | -0.012 | 0.077 |
| ventral-attention : right-cerebellum | 0.507 | 0.034 | 0.024 | 1.285 | 0.199 | -0.016 | 0.080 |
| retrosplenial temporal : right-caudate | 0.492 | -0.033 | 0.023 | -1.371 | 0.170 | -0.075 | 0.012 |
| auditory : left-caudate | 0.490 | 0.032 | 0.023 | 1.279 | 0.201 | -0.014 | 0.078 |
| fronto-parietal : right-amygdala | 0.481 | 0.032 | 0.024 | 1.208 | 0.227 | -0.020 | 0.073 |
| dorsal attention : right-hippocampus | 0.480 | 0.032 | 0.023 | 1.291 | 0.197 | -0.016 | 0.077 |
| cingulo-opercular : left-pallidum | 0.476 | 0.032 | 0.023 | 1.299 | 0.194 | -0.014 | 0.075 |
| dorsal attention : right-pallidum | 0.474 | -0.031 | 0.023 | -1.292 | 0.197 | -0.076 | 0.017 |
| sensorimotor mouth : left-thalamus | 0.472 | 0.031 | 0.024 | 1.216 | 0.224 | -0.019 | 0.073 |
| visual : brain-stem | 0.451 | -0.030 | 0.024 | -1.165 | 0.244 | -0.073 | 0.019 |
| dorsal attention : right-putamen | 0.450 | 0.030 | 0.022 | 1.297 | 0.195 | -0.013 | 0.073 |
| dorsal attention : left-pallidum | 0.443 | -0.029 | 0.024 | -1.199 | 0.231 | -0.079 | 0.017 |
| sensorimotor hand : left-putamen | 0.436 | 0.029 | 0.024 | 1.111 | 0.266 | -0.024 | 0.074 |
| dorsal attention : left-cerebellum | 0.434 | 0.029 | 0.023 | 1.189 | 0.234 | -0.017 | 0.073 |
| salience : right-cerebellum | 0.424 | -0.028 | 0.023 | -1.068 | 0.286 | -0.071 | 0.020 |
| default : left-thalamus | 0.414 | 0.027 | 0.023 | 1.092 | 0.275 | -0.019 | 0.072 |
| dorsal attention : right-accumbens | 0.411 | 0.027 | 0.023 | 1.063 | 0.288 | -0.021 | 0.070 |
| fronto-parietal : right-thalamus | 0.406 | -0.027 | 0.024 | -1.101 | 0.271 | -0.070 | 0.019 |
| cingulo-parietal : brain-stem | 0.402 | -0.027 | 0.024 | -1.007 | 0.314 | -0.068 | 0.023 |
| dorsal attention : right-thalamus | 0.394 | -0.026 | 0.024 | -1.010 | 0.312 | -0.068 | 0.021 |
| auditory : left-thalamus | 0.379 | 0.025 | 0.025 | 1.047 | 0.295 | -0.019 | 0.074 |

| | | | | | | | |
|---|---|---|---|---|---|---|---|
| fronto-parietal : right-cerebellum | 0.378 | 0.025 | 0.024 | 1.028 | 0.304 | -0.021 | 0.073 |
| ventral-attention : right-thalamus | 0.374 | -0.025 | 0.023 | -0.972 | 0.331 | -0.066 | 0.025 |
| default : right-cerebellum | 0.371 | -0.025 | 0.023 | -1.017 | 0.309 | -0.068 | 0.023 |
| cingulo-parietal : right-accumbens | 0.366 | 0.024 | 0.024 | 0.958 | 0.338 | -0.022 | 0.071 |
| ventral-attention : left-thalamus | 0.366 | 0.024 | 0.023 | 1.012 | 0.312 | -0.025 | 0.066 |
| sensorimotor mouth : right-pallidum | 0.361 | -0.024 | 0.024 | -0.943 | 0.346 | -0.069 | 0.024 |
| sensorimotor mouth : right-ventraldc | 0.352 | -0.023 | 0.023 | -0.961 | 0.337 | -0.067 | 0.023 |
| sensorimotor mouth : brain-stem | 0.348 | 0.023 | 0.024 | 0.939 | 0.348 | -0.024 | 0.068 |
| salience : right-caudate | 0.347 | 0.023 | 0.024 | 0.843 | 0.399 | -0.028 | 0.067 |
| visual : left-pallidum | 0.340 | -0.023 | 0.025 | -0.849 | 0.396 | -0.076 | 0.025 |
| fronto-parietal : brain-stem | 0.340 | 0.023 | 0.023 | 0.946 | 0.344 | -0.022 | 0.067 |
| ventral-attention : right-hippocampus | 0.339 | -0.022 | 0.024 | -0.871 | 0.384 | -0.068 | 0.026 |
| cingulo-parietal : left-accumbens | 0.338 | 0.022 | 0.024 | 0.896 | 0.370 | -0.024 | 0.069 |
| sensorimotor mouth : right-accumbens | 0.334 | -0.022 | 0.024 | -0.854 | 0.393 | -0.069 | 0.027 |
| retrosplenial temporal : right-putamen | 0.329 | -0.022 | 0.024 | -0.845 | 0.398 | -0.064 | 0.031 |
| default : left-amygdala | 0.322 | -0.021 | 0.024 | -0.855 | 0.393 | -0.065 | 0.026 |
| retrosplenial temporal : right-amygdala | 0.322 | 0.021 | 0.024 | 0.826 | 0.409 | -0.027 | 0.071 |
| sensorimotor hand : left-accumbens | 0.316 | 0.021 | 0.024 | 0.865 | 0.387 | -0.023 | 0.068 |
| fronto-parietal : right-ventraldc | 0.299 | 0.020 | 0.024 | 0.845 | 0.398 | -0.027 | 0.067 |
| sensorimotor mouth : left-caudate | 0.298 | -0.020 | 0.024 | -0.757 | 0.449 | -0.062 | 0.028 |
| retrosplenial temporal : left-ventraldc | 0.296 | -0.020 | 0.026 | -0.723 | 0.469 | -0.071 | 0.030 |
| retrosplenial temporal : left-putamen | 0.294 | -0.019 | 0.023 | -0.820 | 0.412 | -0.064 | 0.028 |
| dorsal attention : left-caudate | 0.294 | 0.019 | 0.025 | 0.786 | 0.432 | -0.029 | 0.067 |
| auditory : left-pallidum | 0.285 | -0.019 | 0.026 | -0.645 | 0.519 | -0.068 | 0.033 |
| fronto-parietal : left-hippocampus | 0.281 | 0.019 | 0.023 | 0.770 | 0.441 | -0.028 | 0.063 |
| ventral-attention : left-ventraldc | 0.274 | 0.018 | 0.024 | 0.740 | 0.459 | -0.029 | 0.063 |
| salience : right-pallidum | 0.268 | -0.018 | 0.024 | -0.688 | 0.491 | -0.061 | 0.034 |
| visual : left-ventraldc | 0.263 | -0.017 | 0.023 | -0.701 | 0.483 | -0.061 | 0.032 |
| salience : right-accumbens | 0.262 | -0.017 | 0.023 | -0.731 | 0.465 | -0.063 | 0.029 |

| | | | | | | | |
|---|---|---|---|---|---|---|---|
| sensorimotor hand : right-amygdala | 0.259 | 0.017 | 0.024 | 0.655 | 0.513 | -0.031 | 0.063 |
| sensorimotor mouth : right-amygdala | 0.256 | -0.017 | 0.022 | -0.694 | 0.488 | -0.058 | 0.029 |
| cingulo-parietal : left-putamen | 0.247 | -0.016 | 0.023 | -0.700 | 0.484 | -0.063 | 0.025 |
| auditory : right-amygdala | 0.240 | 0.016 | 0.024 | 0.612 | 0.541 | -0.030 | 0.062 |
| dorsal attention : right-cerebellum | 0.230 | -0.015 | 0.025 | -0.520 | 0.603 | -0.062 | 0.037 |
| dorsal attention : right-amygdala | 0.227 | 0.015 | 0.023 | 0.639 | 0.523 | -0.029 | 0.059 |
| ventral-attention : right-pallidum | 0.216 | 0.014 | 0.024 | 0.505 | 0.614 | -0.034 | 0.063 |
| retrosplenial temporal : left-accumbens | 0.212 | 0.014 | 0.024 | 0.573 | 0.566 | -0.031 | 0.061 |
| salience : left-caudate | 0.211 | 0.014 | 0.024 | 0.574 | 0.566 | -0.034 | 0.064 |
| auditory : right-thalamus | 0.206 | 0.014 | 0.025 | 0.569 | 0.570 | -0.031 | 0.063 |
| sensorimotor hand : right-cerebellum | 0.203 | 0.013 | 0.024 | 0.534 | 0.593 | -0.032 | 0.061 |
| salience : left-amygdala | 0.193 | 0.013 | 0.024 | 0.459 | 0.646 | -0.036 | 0.056 |
| auditory : right-caudate | 0.183 | 0.012 | 0.025 | 0.416 | 0.677 | -0.041 | 0.055 |
| visual : left-putamen | 0.182 | 0.012 | 0.023 | 0.502 | 0.616 | -0.035 | 0.055 |
| cingulo-opercular : right-pallidum | 0.180 | 0.012 | 0.025 | 0.448 | 0.654 | -0.036 | 0.058 |
| retrosplenial temporal : left-thalamus | 0.164 | -0.011 | 0.024 | -0.345 | 0.730 | -0.056 | 0.039 |
| sensorimotor mouth : right-cerebellum | 0.156 | -0.010 | 0.024 | -0.389 | 0.697 | -0.054 | 0.037 |
| ventral-attention : right-amygdala | 0.145 | -0.010 | 0.023 | -0.328 | 0.743 | -0.055 | 0.038 |
| dorsal attention : left-ventraldc | 0.143 | -0.009 | 0.023 | -0.388 | 0.698 | -0.053 | 0.035 |
| cingulo-parietal : right-putamen | 0.142 | -0.009 | 0.024 | -0.359 | 0.720 | -0.052 | 0.039 |
| fronto-parietal : left-pallidum | 0.139 | -0.009 | 0.024 | -0.386 | 0.700 | -0.058 | 0.040 |
| default : right-thalamus | 0.139 | -0.009 | 0.023 | -0.384 | 0.701 | -0.055 | 0.035 |
| salience : brain-stem | 0.137 | -0.009 | 0.025 | -0.326 | 0.744 | -0.054 | 0.042 |
| salience : left-pallidum | 0.135 | 0.009 | 0.024 | 0.351 | 0.726 | -0.037 | 0.053 |
| sensorimotor hand : right-thalamus | 0.131 | -0.009 | 0.024 | -0.363 | 0.717 | -0.056 | 0.035 |
| auditory : left-amygdala | 0.128 | 0.008 | 0.024 | 0.278 | 0.781 | -0.042 | 0.051 |
| cingulo-opercular : right-thalamus | 0.122 | -0.008 | 0.023 | -0.291 | 0.771 | -0.051 | 0.039 |
| ventral-attention : left-accumbens | 0.117 | 0.008 | 0.022 | 0.327 | 0.744 | -0.031 | 0.052 |
| salience : left-hippocampus | 0.105 | -0.007 | 0.024 | -0.228 | 0.820 | -0.051 | 0.042 |

| | | | | | | | |
|---|---|---|---|---|---|---|---|
| retrosplenial temporal : left-pallidum | 0.097 | 0.006 | 0.023 | 0.236 | 0.813 | -0.041 | 0.052 |
| default : right-caudate | 0.068 | 0.005 | 0.023 | 0.271 | 0.787 | -0.039 | 0.051 |
| cingulo-opercular : right-cerebellum | 0.068 | 0.004 | 0.024 | 0.145 | 0.885 | -0.043 | 0.052 |
| visual : left-cerebellum | 0.065 | 0.004 | 0.023 | 0.203 | 0.839 | -0.040 | 0.050 |
| dorsal attention : left-putamen | 0.062 | -0.004 | 0.024 | -0.132 | 0.895 | -0.048 | 0.043 |
| salience : right-ventraldc | 0.061 | -0.004 | 0.023 | -0.201 | 0.841 | -0.046 | 0.039 |
| sensorimotor mouth : left-cerebellum | 0.060 | 0.004 | 0.024 | 0.148 | 0.882 | -0.043 | 0.050 |
| sensorimotor mouth : right-putamen | 0.056 | -0.004 | 0.024 | -0.146 | 0.884 | -0.050 | 0.047 |
| auditory : right-accumbens | 0.049 | -0.003 | 0.023 | -0.060 | 0.953 | -0.046 | 0.043 |
| default : right-ventraldc | 0.046 | 0.003 | 0.024 | 0.137 | 0.891 | -0.045 | 0.047 |
| sensorimotor mouth : right-hippocampus | 0.035 | -0.002 | 0.024 | -0.069 | 0.945 | -0.050 | 0.049 |
| ventral-attention : left-cerebellum | 0.022 | -0.001 | 0.023 | -0.067 | 0.947 | -0.048 | 0.044 |
| sensorimotor hand : left-thalamus | 0.022 | -0.001 | 0.024 | -0.002 | 0.998 | -0.046 | 0.048 |
| salience : right-amygdala | 0.018 | 0.001 | 0.023 | 0.080 | 0.936 | -0.045 | 0.047 |
| visual : right-amygdala | 0.018 | -0.001 | 0.024 | -0.049 | 0.961 | -0.046 | 0.045 |
| cingulo-opercular : left-hippocampus | 0.016 | -0.001 | 0.025 | -0.066 | 0.947 | -0.049 | 0.046 |
| cingulo-parietal : left-cerebellum | 0.006 | 0.000 | 0.025 | 0.012 | 0.990 | -0.046 | 0.051 |
| visual : right-cerebellum | 0.002 | 0.000 | 0.024 | 0.044 | 0.965 | -0.045 | 0.048 |

*Notes*. Variable importance in projection (VIP) was used to assess the relative importance of each item in the PLS model. Bold indicates VIP scores considered most influential in terms of their explanatory power (>1). Bootstrapping with 1,000 resampling iterations was used to obtain standard errors, t-values confidence intervals, and p-values.

**Table S31.** Mediation effect of the PLS cortico-limbic regional connectivity component on the association between the PRS component and mental health phenotypes

| *Direct Effects* | | | | | **Est** | **Std Err** | **z** | **Lower 95%** | **Upper 95%** | **p** |
|---|---|---|---|---|---|---|---|---|---|---|
| PRS component | → | | | phenoADHD | -0.149 | 0.042 | -3.501 | -0.232 | -0.065 | 0.000 |
| PRS component | → | | | phenoAnxiety | -0.083 | 0.037 | -2.233 | -0.157 | -0.010 | 0.026 |
| PRS component | → | | | phenoDepression | -0.083 | 0.029 | -2.822 | -0.141 | -0.025 | 0.005 |
| PRS component | → | | | phenoPsychosis | -0.464 | 0.143 | -3.253 | -0.743 | -0.184 | 0.001 |
| *Indirect Effects* | | | | | **Est** | **Std Err** | **z** | **Lower 95%** | **Upper 95%** | **p** |
| PRS component | → | cortico-limbic component | → | phenoADHD | 0.003 | 0.002 | 1.423 | -0.001 | 0.008 | 0.155 |
| PRS component | → | cortico-limbic component | → | phenoAnxiety | 0.003 | 0.002 | 1.555 | -0.001 | 0.008 | 0.120 |
| PRS component | → | cortico-limbic component | → | phenoDepression | 0.003 | 0.002 | 1.798 | 0.000 | 0.007 | 0.072 |
| PRS component | → | cortico-limbic component | → | phenoPsychosis | -0.021 | 0.010 | -2.080 | -0.041 | -0.001 | 0.038 |

*Notes*. Components derived from the PRS-only cortico-limbic regional connectivity PLS model. Controlling for age, sex and 6 PCs.

**Table S32.** PLS: cortico-limbic regional connectivity VIP scores and loadings for the adversity-only model

| | VIP | Loading | Std. err | t | p | 2.50% | 97.50% |
|---|---|---|---|---|---|---|---|
| *Predictor (adversity)* | | | | | | | |
| **adversity** | **-** | **1.000** | 0.000 | - | 0.00 | 1 | 1 |
| *Outcome (regional connectivity)* | | | | | | | |
| **sensorimotor hand : right-accumbens** | **2.554** | **-0.169** | **0.017** | **-9.194** | **0.000** | **-0.191** | **-0.123** |
| **cingulo-parietal : left-thalamus** | **2.371** | **-0.157** | **0.018** | **-8.036** | **0.000** | **-0.180** | **-0.111** |
| **sensorimotor hand : left-pallidum** | **2.356** | **-0.156** | **0.017** | **-8.245** | **0.000** | **-0.175** | **-0.107** |
| **sensorimotor mouth : right-caudate** | **2.264** | **-0.150** | **0.022** | **-6.300** | **0.000** | **-0.180** | **-0.092** |
| **retrosplenial temporal : right-thalamus** | **2.173** | **-0.144** | **0.021** | **-6.295** | **0.000** | **-0.173** | **-0.091** |
| **sensorimotor hand : right-ventraldc** | **2.165** | **-0.143** | **0.022** | **-6.067** | **0.000** | **-0.176** | **-0.091** |
| **ventral-attention : left-caudate** | **2.151** | **-0.142** | **0.023** | **-5.687** | **0.000** | **-0.179** | **-0.089** |
| **sensorimotor mouth : right-thalamus** | **2.145** | **-0.142** | **0.019** | **-6.927** | **0.000** | **-0.169** | **-0.094** |
| **cingulo-opercular : left-amygdala** | **2.135** | **-0.141** | **0.019** | **-6.878** | **0.000** | **-0.162** | **-0.091** |
| **cingulo-parietal : right-pallidum** | **2.074** | **-0.137** | **0.021** | **-6.040** | **0.000** | **-0.167** | **-0.086** |
| **auditory : right-putamen** | **2.029** | **-0.134** | **0.017** | **-7.382** | **0.000** | **-0.154** | **-0.088** |
| **sensorimotor hand : right-putamen** | **1.994** | **-0.132** | **0.021** | **-5.731** | **0.000** | **-0.163** | **-0.078** |
| **cingulo-parietal : left-caudate** | **1.936** | **-0.128** | **0.023** | **-5.113** | **0.000** | **-0.165** | **-0.075** |
| **sensorimotor hand : right-caudate** | **1.906** | **-0.126** | **0.022** | **-5.267** | **0.000** | **-0.154** | **-0.069** |
| **cingulo-opercular : right-amygdala** | **1.890** | **-0.125** | **0.020** | **-5.702** | **0.000** | **-0.152** | **-0.073** |
| **auditory : left-putamen** | **1.878** | **0.124** | **0.024** | **4.857** | **0.000** | **0.068** | **0.165** |
| **ventral-attention : right-ventraldc** | **1.869** | **-0.124** | **0.024** | **-4.704** | **0.000** | **-0.163** | **-0.067** |
| **cingulo-parietal : left-ventraldc** | **1.868** | **-0.124** | **0.022** | **-5.243** | **0.000** | **-0.154** | **-0.071** |
| **retrosplenial temporal : right-cerebellum** | **1.861** | **-0.123** | **0.019** | **-5.964** | **0.000** | **-0.147** | **-0.074** |
| **default : left-pallidum** | **1.848** | **-0.122** | **0.022** | **-5.147** | **0.000** | **-0.153** | **-0.067** |
| **cingulo-opercular : right-hippocampus** | **1.840** | **-0.122** | **0.022** | **-5.140** | **0.000** | **-0.149** | **-0.068** |
| **ventral-attention : left-hippocampus** | **1.806** | **-0.120** | **0.024** | **-4.644** | **0.000** | **-0.163** | **-0.067** |

| | | | | | | | |
|---|---|---|---|---|---|---|---|
| sensorimotor mouth : left-hippocampus | 1.797 | -0.119 | 0.021 | -5.119 | 0.000 | -0.151 | -0.068 |
| sensorimotor mouth : left-amygdala | 1.782 | -0.118 | 0.022 | -4.980 | 0.000 | -0.151 | -0.064 |
| sensorimotor hand : brain-stem | 1.752 | -0.116 | 0.022 | -4.980 | 0.000 | -0.150 | -0.064 |
| visual : left-hippocampus | 1.750 | -0.116 | 0.021 | -5.044 | 0.000 | -0.145 | -0.064 |
| visual : right-pallidum | 1.734 | -0.115 | 0.022 | -4.724 | 0.000 | -0.148 | -0.059 |
| cingulo-opercular : left-caudate | 1.711 | -0.113 | 0.021 | -4.933 | 0.000 | -0.141 | -0.060 |
| cingulo-opercular : left-putamen | 1.705 | -0.113 | 0.020 | -5.136 | 0.000 | -0.139 | -0.058 |
| cingulo-parietal : right-cerebellum | 1.674 | -0.111 | 0.022 | -4.720 | 0.000 | -0.142 | -0.061 |
| salience : left-cerebellum | 1.639 | -0.109 | 0.022 | -4.568 | 0.000 | -0.141 | -0.055 |
| retrosplenial temporal : right-ventraldc | 1.632 | -0.108 | 0.021 | -4.642 | 0.000 | -0.137 | -0.058 |
| default : left-putamen | 1.595 | -0.106 | 0.024 | -4.111 | 0.000 | -0.142 | -0.046 |
| auditory : right-pallidum | 1.589 | -0.105 | 0.021 | -4.506 | 0.000 | -0.138 | -0.054 |
| cingulo-opercular : right-ventraldc | 1.559 | -0.103 | 0.023 | -4.145 | 0.000 | -0.139 | -0.047 |
| sensorimotor hand : left-cerebellum | 1.545 | -0.102 | 0.021 | -4.561 | 0.000 | -0.130 | -0.048 |
| fronto-parietal : right-hippocampus | 1.538 | -0.102 | 0.026 | -3.694 | 0.000 | -0.143 | -0.044 |
| cingulo-opercular : left-accumbens | 1.522 | -0.101 | 0.022 | -4.186 | 0.000 | -0.135 | -0.050 |
| salience : left-accumbens | 1.516 | -0.100 | 0.022 | -4.222 | 0.000 | -0.133 | -0.047 |
| sensorimotor hand : right-hippocampus | 1.455 | -0.096 | 0.023 | -3.901 | 0.000 | -0.131 | -0.043 |
| cingulo-parietal : right-hippocampus | 1.441 | -0.095 | 0.022 | -3.934 | 0.000 | -0.129 | -0.043 |
| salience : left-thalamus | 1.412 | -0.094 | 0.023 | -3.787 | 0.000 | -0.128 | -0.038 |
| salience : left-putamen | 1.411 | -0.093 | 0.025 | -3.515 | 0.000 | -0.137 | -0.041 |
| default : left-accumbens | 1.379 | -0.091 | 0.023 | -3.715 | 0.000 | -0.126 | -0.038 |
| fronto-parietal : right-pallidum | 1.351 | 0.089 | 0.027 | 3.068 | 0.002 | 0.031 | 0.140 |
| dorsal attention : brain-stem | 1.349 | -0.089 | 0.023 | -3.566 | 0.000 | -0.123 | -0.034 |
| cingulo-opercular : brain-stem | 1.348 | -0.089 | 0.025 | -3.238 | 0.001 | -0.130 | -0.030 |
| ventral-attention : left-putamen | 1.343 | 0.089 | 0.024 | 3.442 | 0.001 | 0.037 | 0.131 |
| auditory : left-ventraldc | 1.327 | -0.088 | 0.025 | -3.344 | 0.001 | -0.131 | -0.033 |
| cingulo-opercular : left-ventraldc | 1.327 | 0.088 | 0.025 | 3.253 | 0.001 | 0.034 | 0.132 |
| sensorimotor mouth : left-ventraldc | 1.303 | -0.086 | 0.025 | -3.230 | 0.001 | -0.129 | -0.029 |

| | | | | | | | |
|---|---|---|---|---|---|---|---|
| **dorsal attention : right-accumbens** | **1.283** | **-0.085** | **0.025** | **-3.156** | **0.002** | **-0.132** | **-0.034** |
| **dorsal attention : right-ventraldc** | **1.262** | **-0.084** | **0.023** | **-3.405** | **0.001** | **-0.120** | **-0.033** |
| **default : left-cerebellum** | **1.232** | **0.082** | **0.025** | **2.978** | **0.003** | **0.027** | **0.127** |
| **ventral-attention : brain-stem** | **1.223** | **-0.081** | **0.025** | **-3.008** | **0.003** | **-0.123** | **-0.027** |
| **retrosplenial temporal : left-cerebellum** | **1.213** | **0.080** | **0.024** | **3.154** | **0.002** | **0.028** | **0.122** |
| **default : left-hippocampus** | **1.210** | **-0.080** | **0.026** | **-2.841** | **0.005** | **-0.126** | **-0.023** |
| **visual : right-hippocampus** | **1.183** | **-0.078** | **0.024** | **-3.020** | **0.003** | **-0.117** | **-0.023** |
| **auditory : left-accumbens** | **1.149** | **-0.076** | **0.024** | **-2.850** | **0.004** | **-0.114** | **-0.021** |
| **sensorimotor hand : left-hippocampus** | **1.130** | **0.075** | **0.024** | **2.923** | **0.003** | **0.021** | **0.117** |
| **retrosplenial temporal : brain-stem** | **1.094** | **-0.072** | **0.024** | **-2.793** | **0.005** | **-0.114** | **-0.017** |
| **cingulo-opercular : left-thalamus** | **1.072** | **-0.071** | **0.027** | **-2.528** | **0.012** | **-0.120** | **-0.016** |
| **retrosplenial temporal : right-pallidum** | **1.044** | **-0.069** | **0.026** | **-2.434** | **0.015** | **-0.118** | **-0.013** |
| **sensorimotor hand : left-amygdala** | **1.043** | **0.069** | **0.025** | **2.571** | **0.010** | **0.016** | **0.116** |
| **default : left-ventraldc** | **1.042** | **-0.069** | **0.026** | **-2.466** | **0.014** | **-0.113** | **-0.009** |
| **sensorimotor hand : left-caudate** | **1.036** | **-0.069** | **0.024** | **-2.627** | **0.009** | **-0.108** | **-0.015** |
| **ventral-attention : left-thalamus** | **1.035** | **-0.069** | **0.024** | **-2.689** | **0.007** | **-0.112** | **-0.020** |
| **auditory : left-cerebellum** | **1.030** | **-0.068** | **0.024** | **-2.633** | **0.008** | **-0.109** | **-0.017** |
| **auditory : right-accumbens** | **1.026** | **-0.068** | **0.027** | **-2.350** | **0.019** | **-0.118** | **-0.012** |
| **dorsal attention : left-hippocampus** | **1.016** | **0.067** | **0.025** | **2.515** | **0.012** | **0.016** | **0.112** |
| **salience : left-amygdala** | **1.015** | **-0.067** | **0.027** | **-2.328** | **0.020** | **-0.117** | **-0.011** |
| **ventral-attention : left-cerebellum** | **1.010** | **0.067** | **0.025** | **2.440** | **0.015** | **0.013** | **0.111** |
| **cingulo-parietal : right-caudate** | **1.009** | **-0.067** | **0.025** | **-2.463** | **0.014** | **-0.108** | **-0.012** |
| **salience : left-ventraldc** | **1.004** | **0.067** | **0.025** | **2.452** | **0.014** | **0.013** | **0.109** |
| default : right-accumbens | 0.999 | -0.066 | 0.025 | -2.382 | 0.017 | -0.108 | -0.010 |
| auditory : right-ventraldc | 0.996 | 0.066 | 0.027 | 2.257 | 0.024 | 0.007 | 0.113 |
| fronto-parietal : right-amygdala | 0.943 | -0.062 | 0.027 | -2.143 | 0.032 | -0.109 | -0.002 |
| default : right-amygdala | 0.919 | -0.061 | 0.026 | -2.134 | 0.033 | -0.105 | -0.004 |
| retrosplenial temporal : left-caudate | 0.914 | 0.061 | 0.024 | 2.380 | 0.017 | 0.009 | 0.100 |
| visual : right-cerebellum | 0.908 | -0.060 | 0.027 | -2.086 | 0.037 | -0.112 | -0.006 |

| | | | | | | | |
|---|---|---|---|---|---|---|---|
| visual : left-amygdala | 0.908 | -0.060 | 0.027 | -2.095 | 0.036 | -0.104 | -0.003 |
| auditory : left-hippocampus | 0.887 | 0.059 | 0.025 | 2.178 | 0.029 | 0.008 | 0.103 |
| fronto-parietal : left-putamen | 0.885 | -0.059 | 0.026 | -2.099 | 0.036 | -0.104 | -0.002 |
| fronto-parietal : right-accumbens | 0.877 | -0.058 | 0.026 | -2.001 | 0.045 | -0.103 | 0.000 |
| ventral-attention : left-amygdala | 0.855 | 0.057 | 0.025 | 2.118 | 0.034 | 0.004 | 0.103 |
| retrosplenial temporal : left-amygdala | 0.853 | 0.057 | 0.025 | 2.190 | 0.029 | 0.004 | 0.100 |
| default : right-caudate | 0.840 | -0.056 | 0.028 | -1.888 | 0.059 | -0.108 | -0.001 |
| ventral-attention : right-accumbens | 0.825 | 0.055 | 0.026 | 1.957 | 0.050 | -0.003 | 0.100 |
| sensorimotor hand : left-putamen | 0.820 | -0.054 | 0.025 | -2.012 | 0.044 | -0.101 | -0.001 |
| retrosplenial temporal : right-accumbens | 0.812 | -0.054 | 0.026 | -1.923 | 0.055 | -0.101 | 0.000 |
| auditory : left-caudate | 0.799 | 0.053 | 0.026 | 1.899 | 0.058 | 0.000 | 0.099 |
| auditory : brain-stem | 0.796 | 0.053 | 0.025 | 1.962 | 0.050 | -0.003 | 0.095 |
| sensorimotor mouth : right-putamen | 0.777 | 0.051 | 0.025 | 1.952 | 0.051 | -0.001 | 0.100 |
| default : right-hippocampus | 0.765 | -0.051 | 0.027 | -1.782 | 0.075 | -0.102 | 0.006 |
| cingulo-parietal : right-amygdala | 0.760 | 0.050 | 0.024 | 1.893 | 0.058 | 0.000 | 0.094 |
| visual : left-caudate | 0.752 | -0.050 | 0.026 | -1.847 | 0.065 | -0.098 | 0.006 |
| fronto-parietal : left-ventraldc | 0.749 | -0.050 | 0.026 | -1.755 | 0.079 | -0.093 | 0.008 |
| visual : right-accumbens | 0.746 | -0.049 | 0.027 | -1.690 | 0.091 | -0.097 | 0.010 |
| retrosplenial temporal : right-hippocampus | 0.746 | 0.049 | 0.027 | 1.647 | 0.100 | -0.011 | 0.095 |
| cingulo-opercular : right-putamen | 0.733 | -0.049 | 0.026 | -1.716 | 0.086 | -0.095 | 0.011 |
| dorsal attention : left-putamen | 0.727 | 0.048 | 0.025 | 1.821 | 0.069 | -0.002 | 0.096 |
| cingulo-parietal : right-accumbens | 0.713 | 0.047 | 0.026 | 1.650 | 0.099 | -0.009 | 0.095 |
| default : left-caudate | 0.702 | 0.046 | 0.026 | 1.570 | 0.117 | -0.009 | 0.092 |
| dorsal attention : left-pallidum | 0.701 | -0.046 | 0.026 | -1.702 | 0.089 | -0.097 | 0.003 |
| auditory : right-cerebellum | 0.685 | 0.045 | 0.024 | 1.818 | 0.069 | -0.006 | 0.090 |
| sensorimotor mouth : right-pallidum | 0.672 | 0.045 | 0.025 | 1.636 | 0.102 | -0.010 | 0.092 |
| cingulo-opercular : left-hippocampus | 0.649 | -0.043 | 0.027 | -1.526 | 0.127 | -0.096 | 0.012 |
| auditory : left-pallidum | 0.648 | -0.043 | 0.027 | -1.529 | 0.126 | -0.092 | 0.010 |
| visual : left-thalamus | 0.640 | -0.042 | 0.027 | -1.433 | 0.152 | -0.090 | 0.013 |

| | | | | | | | |
|---|---|---|---|---|---|---|---|
| default : right-thalamus | 0.613 | 0.041 | 0.025 | 1.514 | 0.130 | -0.011 | 0.090 |
| ventral-attention : right-caudate | 0.613 | -0.041 | 0.025 | -1.507 | 0.132 | -0.082 | 0.013 |
| salience : right-caudate | 0.608 | -0.040 | 0.026 | -1.430 | 0.153 | -0.088 | 0.018 |
| auditory : left-thalamus | 0.608 | -0.040 | 0.024 | -1.511 | 0.131 | -0.084 | 0.011 |
| salience : left-hippocampus | 0.589 | 0.039 | 0.026 | 1.372 | 0.170 | -0.017 | 0.088 |
| cingulo-opercular : right-cerebellum | 0.587 | 0.039 | 0.025 | 1.480 | 0.139 | -0.014 | 0.088 |
| ventral-attention : right-thalamus | 0.584 | 0.039 | 0.024 | 1.511 | 0.131 | -0.010 | 0.082 |
| retrosplenial temporal : right-putamen | 0.581 | 0.039 | 0.025 | 1.373 | 0.170 | -0.022 | 0.078 |
| retrosplenial temporal : left-thalamus | 0.576 | -0.038 | 0.027 | -1.370 | 0.171 | -0.089 | 0.017 |
| visual : left-cerebellum | 0.565 | -0.037 | 0.028 | -1.225 | 0.221 | -0.090 | 0.019 |
| auditory : right-caudate | 0.561 | -0.037 | 0.025 | -1.433 | 0.152 | -0.081 | 0.013 |
| cingulo-opercular : left-cerebellum | 0.548 | -0.036 | 0.026 | -1.327 | 0.185 | -0.083 | 0.018 |
| sensorimotor mouth : right-amygdala | 0.544 | 0.036 | 0.026 | 1.333 | 0.183 | -0.015 | 0.085 |
| cingulo-parietal : left-amygdala | 0.543 | -0.036 | 0.026 | -1.340 | 0.180 | -0.082 | 0.012 |
| sensorimotor hand : right-pallidum | 0.537 | -0.036 | 0.026 | -1.292 | 0.196 | -0.083 | 0.014 |
| visual : right-putamen | 0.530 | -0.035 | 0.024 | -1.321 | 0.186 | -0.080 | 0.015 |
| visual : right-ventraldc | 0.527 | -0.035 | 0.025 | -1.312 | 0.190 | -0.081 | 0.016 |
| cingulo-parietal : right-ventraldc | 0.524 | 0.035 | 0.026 | 1.275 | 0.203 | -0.017 | 0.084 |
| dorsal attention : right-amygdala | 0.508 | 0.034 | 0.025 | 1.265 | 0.206 | -0.016 | 0.079 |
| fronto-parietal : left-cerebellum | 0.507 | 0.034 | 0.025 | 1.242 | 0.214 | -0.018 | 0.079 |
| auditory : right-hippocampus | 0.505 | 0.033 | 0.025 | 1.247 | 0.213 | -0.020 | 0.079 |
| retrosplenial temporal : left-hippocampus | 0.503 | -0.033 | 0.026 | -1.172 | 0.241 | -0.079 | 0.022 |
| dorsal attention : left-thalamus | 0.494 | -0.033 | 0.028 | -1.168 | 0.243 | -0.086 | 0.022 |
| fronto-parietal : brain-stem | 0.493 | -0.033 | 0.027 | -1.170 | 0.242 | -0.085 | 0.020 |
| dorsal attention : left-caudate | 0.491 | -0.033 | 0.028 | -1.109 | 0.268 | -0.085 | 0.025 |
| retrosplenial temporal : left-accumbens | 0.488 | 0.032 | 0.027 | 1.160 | 0.246 | -0.024 | 0.083 |
| default : brain-stem | 0.486 | 0.032 | 0.026 | 1.177 | 0.239 | -0.018 | 0.083 |
| fronto-parietal : left-caudate | 0.478 | -0.032 | 0.026 | -1.146 | 0.252 | -0.076 | 0.024 |
| visual : right-thalamus | 0.478 | 0.032 | 0.024 | 1.216 | 0.224 | -0.020 | 0.074 |

| | | | | | | | |
|---|---|---|---|---|---|---|---|
| dorsal attention : left-amygdala | 0.477 | 0.032 | 0.027 | 1.082 | 0.279 | -0.024 | 0.084 |
| cingulo-opercular : right-thalamus | 0.474 | -0.031 | 0.027 | -1.102 | 0.271 | -0.083 | 0.022 |
| fronto-parietal : right-ventraldc | 0.467 | -0.031 | 0.024 | -1.210 | 0.227 | -0.076 | 0.020 |
| cingulo-opercular : right-accumbens | 0.459 | -0.030 | 0.026 | -1.114 | 0.265 | -0.076 | 0.022 |
| fronto-parietal : left-accumbens | 0.452 | 0.030 | 0.025 | 1.143 | 0.253 | -0.021 | 0.073 |
| sensorimotor mouth : left-pallidum | 0.452 | -0.030 | 0.026 | -1.113 | 0.266 | -0.078 | 0.021 |
| dorsal attention : left-ventraldc | 0.439 | -0.029 | 0.027 | -1.024 | 0.306 | -0.079 | 0.027 |
| ventral-attention : left-pallidum | 0.432 | 0.029 | 0.026 | 1.030 | 0.303 | -0.023 | 0.076 |
| cingulo-parietal : brain-stem | 0.429 | -0.028 | 0.026 | -1.013 | 0.311 | -0.079 | 0.023 |
| visual : left-pallidum | 0.427 | -0.028 | 0.026 | -1.024 | 0.306 | -0.077 | 0.024 |
| sensorimotor mouth : right-ventraldc | 0.416 | 0.028 | 0.025 | 0.962 | 0.336 | -0.027 | 0.074 |
| retrosplenial temporal : left-ventraldc | 0.413 | -0.027 | 0.024 | -1.119 | 0.263 | -0.074 | 0.020 |
| sensorimotor hand : left-accumbens | 0.407 | -0.027 | 0.026 | -0.949 | 0.343 | -0.077 | 0.026 |
| ventral-attention : right-cerebellum | 0.405 | -0.027 | 0.026 | -0.963 | 0.335 | -0.073 | 0.025 |
| salience : right-cerebellum | 0.403 | 0.027 | 0.026 | 0.964 | 0.335 | -0.025 | 0.076 |
| salience : left-pallidum | 0.391 | 0.026 | 0.027 | 0.842 | 0.400 | -0.030 | 0.078 |
| visual : left-ventraldc | 0.390 | 0.026 | 0.026 | 0.932 | 0.351 | -0.028 | 0.076 |
| cingulo-opercular : right-caudate | 0.390 | 0.026 | 0.026 | 0.950 | 0.342 | -0.028 | 0.076 |
| sensorimotor mouth : left-caudate | 0.372 | -0.025 | 0.027 | -0.857 | 0.391 | -0.076 | 0.028 |
| cingulo-opercular : left-pallidum | 0.353 | 0.023 | 0.026 | 0.919 | 0.358 | -0.026 | 0.072 |
| retrosplenial temporal : right-caudate | 0.347 | -0.023 | 0.027 | -0.803 | 0.422 | -0.076 | 0.032 |
| default : right-putamen | 0.346 | 0.023 | 0.027 | 0.776 | 0.438 | -0.033 | 0.075 |
| sensorimotor hand : right-cerebellum | 0.345 | -0.023 | 0.027 | -0.745 | 0.456 | -0.072 | 0.036 |
| salience : right-ventraldc | 0.329 | 0.022 | 0.025 | 0.817 | 0.414 | -0.029 | 0.069 |
| visual : right-amygdala | 0.321 | 0.021 | 0.027 | 0.717 | 0.473 | -0.036 | 0.074 |
| salience : brain-stem | 0.319 | -0.021 | 0.026 | -0.754 | 0.451 | -0.068 | 0.032 |
| sensorimotor mouth : right-hippocampus | 0.312 | -0.021 | 0.026 | -0.747 | 0.455 | -0.070 | 0.031 |
| retrosplenial temporal : left-putamen | 0.309 | 0.020 | 0.025 | 0.802 | 0.423 | -0.028 | 0.069 |
| sensorimotor mouth : left-thalamus | 0.309 | 0.020 | 0.025 | 0.741 | 0.459 | -0.030 | 0.068 |

| | | | | | | | |
|---|---|---|---|---|---|---|---|
| default : right-pallidum | 0.303 | -0.020 | 0.027 | -0.705 | 0.481 | -0.070 | 0.034 |
| cingulo-parietal : left-pallidum | 0.295 | 0.020 | 0.025 | 0.716 | 0.474 | -0.034 | 0.063 |
| fronto-parietal : right-putamen | 0.295 | 0.020 | 0.026 | 0.678 | 0.498 | -0.034 | 0.071 |
| auditory : left-amygdala | 0.294 | 0.019 | 0.026 | 0.736 | 0.462 | -0.029 | 0.071 |
| sensorimotor hand : right-thalamus | 0.292 | 0.019 | 0.024 | 0.731 | 0.465 | -0.030 | 0.066 |
| default : right-cerebellum | 0.278 | -0.018 | 0.026 | -0.722 | 0.470 | -0.072 | 0.030 |
| ventral-attention : right-hippocampus | 0.272 | -0.018 | 0.025 | -0.639 | 0.523 | -0.065 | 0.030 |
| fronto-parietal : right-caudate | 0.265 | -0.018 | 0.025 | -0.631 | 0.528 | -0.064 | 0.033 |
| salience : right-accumbens | 0.262 | 0.017 | 0.025 | 0.622 | 0.534 | -0.036 | 0.063 |
| cingulo-parietal : right-thalamus | 0.256 | -0.017 | 0.025 | -0.616 | 0.538 | -0.064 | 0.033 |
| ventral-attention : right-amygdala | 0.255 | 0.017 | 0.025 | 0.611 | 0.541 | -0.032 | 0.068 |
| ventral-attention : left-accumbens | 0.246 | 0.016 | 0.026 | 0.547 | 0.584 | -0.040 | 0.063 |
| salience : right-amygdala | 0.233 | 0.015 | 0.025 | 0.612 | 0.541 | -0.031 | 0.068 |
| retrosplenial temporal : left-pallidum | 0.229 | 0.015 | 0.027 | 0.505 | 0.614 | -0.037 | 0.066 |
| auditory : right-amygdala | 0.217 | 0.014 | 0.026 | 0.518 | 0.605 | -0.036 | 0.063 |
| default : left-amygdala | 0.215 | -0.014 | 0.026 | -0.570 | 0.569 | -0.063 | 0.032 |
| salience : left-caudate | 0.199 | 0.013 | 0.025 | 0.502 | 0.615 | -0.036 | 0.061 |
| auditory : right-thalamus | 0.197 | 0.013 | 0.026 | 0.506 | 0.613 | -0.037 | 0.064 |
| cingulo-parietal : left-cerebellum | 0.193 | -0.013 | 0.027 | -0.536 | 0.592 | -0.069 | 0.039 |
| fronto-parietal : left-thalamus | 0.193 | 0.013 | 0.024 | 0.459 | 0.646 | -0.037 | 0.058 |
| salience : right-thalamus | 0.191 | -0.013 | 0.026 | -0.418 | 0.676 | -0.058 | 0.040 |
| salience : right-pallidum | 0.184 | -0.012 | 0.025 | -0.505 | 0.614 | -0.058 | 0.037 |
| retrosplenial temporal : right-amygdala | 0.184 | 0.012 | 0.024 | 0.462 | 0.644 | -0.035 | 0.057 |
| dorsal attention : left-cerebellum | 0.178 | -0.012 | 0.026 | -0.392 | 0.695 | -0.059 | 0.045 |
| ventral-attention : right-pallidum | 0.162 | -0.011 | 0.027 | -0.336 | 0.737 | -0.064 | 0.044 |
| sensorimotor mouth : left-cerebellum | 0.160 | 0.011 | 0.026 | 0.404 | 0.686 | -0.039 | 0.059 |
| sensorimotor mouth : left-putamen | 0.159 | 0.011 | 0.027 | 0.370 | 0.711 | -0.044 | 0.065 |
| dorsal attention : right-thalamus | 0.126 | -0.008 | 0.027 | -0.333 | 0.739 | -0.061 | 0.045 |
| visual : brain-stem | 0.123 | 0.008 | 0.026 | 0.216 | 0.829 | -0.046 | 0.056 |

| | | | | | | | |
|---|---|---|---|---|---|---|---|
| ventral-attention : right-putamen | 0.118 | 0.008 | 0.026 | 0.266 | 0.790 | -0.042 | 0.056 |
| ventral-attention : left-ventraldc | 0.106 | 0.007 | 0.025 | 0.262 | 0.793 | -0.041 | 0.057 |
| cingulo-parietal : left-accumbens | 0.105 | -0.007 | 0.025 | -0.310 | 0.757 | -0.055 | 0.044 |
| default : right-ventraldc | 0.102 | 0.007 | 0.026 | 0.239 | 0.811 | -0.047 | 0.053 |
| sensorimotor mouth : right-accumbens | 0.101 | -0.007 | 0.027 | -0.271 | 0.786 | -0.062 | 0.044 |
| fronto-parietal : right-thalamus | 0.099 | -0.007 | 0.027 | -0.221 | 0.825 | -0.060 | 0.045 |
| dorsal attention : right-caudate | 0.088 | 0.006 | 0.027 | 0.241 | 0.809 | -0.044 | 0.061 |
| salience : right-hippocampus | 0.084 | 0.006 | 0.026 | 0.237 | 0.813 | -0.045 | 0.057 |
| cingulo-parietal : left-putamen | 0.081 | 0.005 | 0.027 | 0.220 | 0.826 | -0.044 | 0.059 |
| fronto-parietal : left-hippocampus | 0.079 | 0.005 | 0.025 | 0.175 | 0.861 | -0.045 | 0.054 |
| sensorimotor mouth : left-accumbens | 0.078 | -0.005 | 0.025 | -0.190 | 0.850 | -0.055 | 0.044 |
| cingulo-parietal : right-putamen | 0.078 | 0.005 | 0.028 | 0.142 | 0.887 | -0.047 | 0.057 |
| sensorimotor mouth : right-cerebellum | 0.072 | 0.005 | 0.028 | 0.117 | 0.907 | -0.048 | 0.058 |
| sensorimotor mouth : brain-stem | 0.071 | -0.005 | 0.025 | -0.211 | 0.833 | -0.055 | 0.044 |
| visual : left-accumbens | 0.069 | -0.005 | 0.025 | -0.183 | 0.855 | -0.057 | 0.043 |
| fronto-parietal : left-amygdala | 0.068 | 0.005 | 0.026 | 0.208 | 0.835 | -0.042 | 0.055 |
| visual : right-caudate | 0.066 | -0.004 | 0.027 | -0.149 | 0.882 | -0.055 | 0.050 |
| default : left-thalamus | 0.063 | -0.004 | 0.028 | -0.171 | 0.864 | -0.061 | 0.049 |
| sensorimotor hand : left-ventraldc | 0.059 | 0.004 | 0.026 | 0.159 | 0.873 | -0.052 | 0.050 |
| dorsal attention : left-accumbens | 0.057 | -0.004 | 0.027 | -0.104 | 0.917 | -0.051 | 0.055 |
| sensorimotor hand : left-thalamus | 0.051 | -0.003 | 0.027 | -0.124 | 0.901 | -0.055 | 0.049 |
| visual : left-putamen | 0.050 | -0.003 | 0.027 | -0.114 | 0.909 | -0.056 | 0.048 |
| dorsal attention : right-cerebellum | 0.050 | -0.003 | 0.027 | -0.101 | 0.920 | -0.061 | 0.050 |
| dorsal attention : right-hippocampus | 0.047 | 0.003 | 0.027 | 0.124 | 0.902 | -0.048 | 0.058 |
| fronto-parietal : right-cerebellum | 0.031 | 0.002 | 0.026 | 0.068 | 0.946 | -0.046 | 0.055 |
| cingulo-parietal : left-hippocampus | 0.031 | 0.002 | 0.027 | 0.080 | 0.936 | -0.050 | 0.057 |
| cingulo-opercular : right-pallidum | 0.029 | 0.002 | 0.024 | 0.017 | 0.987 | -0.045 | 0.047 |
| sensorimotor hand : right-amygdala | 0.028 | -0.002 | 0.027 | -0.152 | 0.879 | -0.059 | 0.048 |
| fronto-parietal : left-pallidum | 0.021 | 0.001 | 0.027 | 0.048 | 0.962 | -0.052 | 0.055 |

| | | | | | | | |
|---|---|---|---|---|---|---|---|
| salience : right-putamen | 0.007 | 0.000 | 0.027 | 0.027 | 0.978 | -0.055 | 0.051 |
| dorsal attention : right-pallidum | 0.007 | 0.000 | 0.029 | -0.019 | 0.985 | -0.055 | 0.054 |
| dorsal attention : right-putamen | 0.001 | 0.000 | 0.026 | 0.045 | 0.964 | -0.048 | 0.058 |

*Notes*. Variable importance in projection (VIP) was used to assess the relative importance of each item in the PLS model. Bold indicates VIP scores considered most influential in terms of their explanatory power (>1). Bootstrapping with 1,000 resampling iterations was used to obtain standard errors, t-values confidence intervals, and p-values.

**Table S33.** Mediation effect of the PLS cortico-limbic regional connectivity component on the association between the adversity component and mental health phenotypes

| *Direct Effects* | | | | **Est** | **Std Err** | **z** | **Lower 95%** | **Upper 95%** | **p** |
|---|---|---|---|---|---|---|---|---|---|
| adversity component | → | | phenoADHD | -0.106 | 0.048 | -2.197 | -0.201 | -0.011 | 0.028 |
| adversity component | → | | phenoAnxiety | -0.082 | 0.043 | -1.923 | -0.166 | 0.002 | 0.054 |
| adversity component | → | | phenoDepression | -0.061 | 0.034 | -1.780 | -0.128 | 0.006 | 0.075 |
| adversity component | → | | phenoPsychosis | -0.519 | 0.159 | -3.262 | -0.831 | -0.207 | 0.001 |
| *Indirect Effects* | | | | **Est** | **Std Err** | **z** | **Lower 95%** | **Upper 95%** | **p** |
| adversity component | → | cortico-limbic component | → phenoADHD | 0.003 | 0.004 | 0.765 | -0.004 | 0.010 | 0.444 |
| adversity component | → | cortico-limbic component | → phenoAnxiety | 0.002 | 0.003 | 0.755 | -0.004 | 0.009 | 0.450 |
| adversity component | → | cortico-limbic component | → phenoDepression | 0.002 | 0.003 | 0.932 | -0.003 | 0.008 | 0.351 |
| adversity component | → | cortico-limbic component | → phenoPsychosis | -0.042 | 0.015 | -2.728 | -0.072 | -0.012 | 0.006 |

*Notes*. Components derived from the adversity-only cortico-limbic regional connectivity PLS model. Controlling for age, sex and 6 PCs.

**Table S34.** Significance test for canonical variates for the entire sample with population substructure CVs included as covariates

| CV | statistic | approx | df1 | df2 | canonical coefficient | P value |
|---|---|---|---|---|---|---|
| 1 | 0.976 | 9.847 | 16 | 19935 | 0.130 | 0.000 |
| 2 | 0.993 | 5.045 | 9 | 15883 | 0.080 | 0.000 |
| 3 | 1.000 | 0.744 | 4 | 13054 | 0.018 | 0.562 |
| 4 | 1.000 | 0.869 | 1 | 6528 | 0.012 | 0.351 |

*Notes*. Wilk's lambda using F=approximation (Rao's F). CV= Canonical variate.

**Table S35.** Scaled canonical coefficients for the entire sample with population substructure CVs included as covariates

| | CV1 | CV2 |
|---|---|---|
| prsADHD | -0.896 | 0.605 |
| prsAnxiety | 0.115 | -0.476 |
| prsDepression | -0.415 | -0.851 |
| prsPsychosis | -0.169 | -0.009 |
| phenoADHD | -0.739 | 0.861 |
| phenoAnxiety | 0.089 | -0.543 |
| phenoDepression | -0.121 | -0.701 |
| phenoPsychosis | -0.546 | -0.258 |

**Table S36.** Canonical loadings for the entire sample with population substructure CVs included as covariates

| | | | Pearson's r | Lower 95% CI | Upper 95% CI | p | |
|---|---|---|---|---|---|---|---|
| *Canonical Variate 1* | | | | | | | |
| CV1prs | - | prsADHD | -0.859 | -0.865 | -0.853 | 0.000 | *** |
| CV1prs | - | prsAnxiety | -0.226 | -0.249 | -0.203 | 0.000 | *** |
| CV1prs | - | prsDepression | -0.481 | -0.500 | -0.462 | 0.000 | *** |
| CV1prs | - | prsPsychosis | -0.337 | -0.358 | -0.315 | 0.000 | *** |
| CV1prs | - | phenoADHD | -0.109 | -0.133 | -0.085 | 0.000 | *** |
| CV1prs | - | phenoAnxiety | -0.048 | -0.073 | -0.024 | 0.000 | *** |
| CV1prs | - | phenoDepression | -0.065 | -0.089 | -0.041 | 0.000 | *** |
| CV1prs | - | phenoPsychosis | -0.085 | -0.109 | -0.060 | 0.000 | *** |
| CV1pheno | - | prsADHD | -0.112 | -0.136 | -0.088 | 0.000 | *** |
| CV1pheno | - | prsAnxiety | -0.029 | -0.054 | -0.005 | 0.017 | |
| CV1pheno | - | prsDepression | -0.063 | -0.087 | -0.038 | 0.000 | *** |
| CV1pheno | - | prsPsychosis | -0.044 | -0.068 | -0.020 | 0.000 | *** |
| CV1pheno | - | phenoADHD | -0.837 | -0.844 | -0.829 | 0.000 | *** |
| CV1pheno | - | phenoAnxiety | -0.372 | -0.393 | -0.351 | 0.000 | *** |
| CV1pheno | - | phenoDepression | -0.501 | -0.519 | -0.482 | 0.000 | *** |
| CV1pheno | - | phenoPsychosis | -0.649 | -0.663 | -0.635 | 0.000 | *** |
| *Canonical Variate 2* | | | | | | | |
| CV2prs | - | prsADHD | 0.294 | 0.272 | 0.316 | 0.000 | *** |
| CV2prs | - | prsAnxiety | -0.507 | -0.525 | -0.489 | 0.000 | *** |
| CV2prs | - | prsDepression | -0.680 | -0.692 | -0.666 | 0.000 | *** |
| CV2prs | - | prsPsychosis | -0.237 | -0.260 | -0.214 | 0.000 | *** |
| CV2prs | - | phenoADHD | 0.016 | -0.008 | 0.041 | 0.184 | |
| CV2prs | - | phenoAnxiety | -0.049 | -0.073 | -0.025 | 0.000 | *** |
| CV2prs | - | phenoDepression | -0.049 | -0.074 | -0.025 | 0.000 | *** |

| | | | | | | | |
|---|---|---|---|---|---|---|---|
| CV2prs | - | phenoPsychosis | -0.019 | -0.044 | 0.005 | 0.119 | |
| CV2pheno | - | prsADHD | 0.024 | -0.001 | 0.048 | 0.056 | |
| CV2pheno | - | prsAnxiety | -0.041 | -0.065 | -0.017 | 0.001 | ** |
| CV2pheno | - | prsDepression | -0.055 | -0.079 | -0.030 | 0.000 | *** |
| CV2pheno | - | prsPsychosis | -0.019 | -0.043 | 0.005 | 0.123 | |
| CV2pheno | - | phenoADHD | 0.204 | 0.181 | 0.227 | 0.000 | *** |
| CV2pheno | - | phenoAnxiety | -0.611 | -0.626 | -0.596 | 0.000 | *** |
| CV2pheno | - | phenoDepression | -0.614 | -0.629 | -0.599 | 0.000 | *** |
| CV2pheno | - | phenoPsychosis | -0.240 | -0.263 | -0.217 | 0.000 | *** |

*Notes*. CV1= First canonical variate. CV2=Second canonical variate. CV#prs= Canonical component for PRS scores. CV#pheno= Canonical component for phenotypes. * p < .05, ** p < .01, *** p < .001.

**Table S37.** Significance test for canonical variates for European ancestry participants with population substructure CVs included as covariates

| **CV** | **statistic** | **approx** | **df1** | **df2** | **canonical coefficient** | **P value** |
|---|---|---|---|---|---|---|
| 1 | 0.974 | 7.035 | 16 | 13043 | 0.127 | 0.000 |
| 2 | 0.990 | 4.702 | 9 | 10392 | 0.097 | 0.000 |
| 3 | 1.000 | 0.443 | 4 | 8542 | 0.020 | 0.778 |
| 4 | 1.000 | 0.004 | 1 | 4272 | 0.001 | 0.950 |

*Notes*. Wilk's lambda using F=approximation (Rao's F). CV= Canonical variate.

**Table S38.** Scaled canonical coefficients for European ancestry participants with population substructure CVs included as covariates

| | CV1 | CV2 |
|---|---|---|
| prsADHD | -0.896 | 0.605 |
| prsAnxiety | 0.115 | -0.476 |
| prsDepression | -0.415 | -0.851 |
| prsPsychosis | -0.169 | -0.009 |
| phenoADHD | -0.739 | 0.861 |
| phenoAnxiety | 0.089 | -0.543 |
| phenoDepression | -0.121 | -0.701 |
| phenoPsychosis | -0.546 | -0.258 |

**Table S39.** Canonical loadings for European ancestry participants with population substructure CVs included as covariates

| | | | Pearson's r | Lower 95% CI | Upper 95% CI | p | |
|---|---|---|---|---|---|---|---|
| *Canonical Variate 1* | | | | | | | |
| CV1prs | - | prsADHD | -0.891 | -0.897 | -0.885 | 0.000 | *** |
| CV1prs | - | prsAnxiety | -0.189 | -0.218 | -0.160 | 0.000 | *** |
| CV1prs | - | prsDepression | -0.532 | -0.553 | -0.510 | 0.000 | *** |
| CV1prs | - | prsPsychosis | -0.323 | -0.349 | -0.295 | 0.000 | *** |
| CV1prs | - | phenoADHD | -0.107 | -0.136 | -0.077 | 0.000 | *** |
| CV1prs | - | phenoAnxiety | -0.031 | -0.061 | -0.001 | 0.043 | * |
| CV1prs | - | phenoDepression | -0.066 | -0.096 | -0.036 | 0.000 | *** |
| CV1prs | - | phenoPsychosis | -0.077 | -0.107 | -0.047 | 0.000 | *** |
| CV1pheno | - | prsADHD | -0.114 | -0.143 | -0.084 | 0.000 | *** |
| CV1pheno | - | prsAnxiety | -0.024 | -0.054 | 0.006 | 0.115 | |

| | | | | | | | |
|---|---|---|---|---|---|---|---|
| CV1pheno | - | prsDepression | -0.068 | -0.097 | -0.038 | 0.000 | *** |
| CV1pheno | - | prsPsychosis | -0.041 | -0.071 | -0.011 | 0.007 | ** |
| CV1pheno | - | phenoADHD | -0.838 | -0.847 | -0.829 | 0.000 | *** |
| CV1pheno | - | phenoAnxiety | -0.243 | -0.271 | -0.215 | 0.000 | *** |
| CV1pheno | - | phenoDepression | -0.521 | -0.543 | -0.499 | 0.000 | *** |
| CV1pheno | - | phenoPsychosis | -0.603 | -0.622 | -0.584 | 0.000 | *** |
| *Canonical Variate 2* | | | | | | | |
| CV2prs | - | prsADHD | 0.264 | 0.236 | 0.292 | 0.000 | *** |
| CV2prs | - | prsAnxiety | -0.554 | -0.574 | -0.533 | 0.000 | *** |
| CV2prs | - | prsDepression | -0.783 | -0.794 | -0.771 | 0.000 | *** |
| CV2prs | - | prsPsychosis | -0.105 | -0.135 | -0.076 | 0.000 | *** |
| CV2prs | - | phenoADHD | 0.020 | -0.010 | 0.050 | 0.195 | |
| CV2prs | - | phenoAnxiety | -0.062 | -0.092 | -0.032 | 0.000 | *** |
| CV2prs | - | phenoDepression | -0.054 | -0.083 | -0.024 | 0.000 | *** |
| CV2prs | - | phenoPsychosis | -0.037 | -0.067 | -0.007 | 0.015 | |
| CV2pheno | - | prsADHD | 0.026 | -0.004 | 0.056 | 0.093 | |
| CV2pheno | - | prsAnxiety | -0.054 | -0.084 | -0.024 | 0.000 | *** |
| CV2pheno | - | prsDepression | -0.076 | -0.106 | -0.046 | 0.000 | *** |
| CV2pheno | - | prsPsychosis | -0.010 | -0.040 | 0.020 | 0.503 | |
| CV2pheno | - | phenoADHD | 0.204 | 0.175 | 0.232 | 0.000 | *** |
| CV2pheno | - | phenoAnxiety | -0.641 | -0.659 | -0.623 | 0.000 | *** |
| CV2pheno | - | phenoDepression | -0.552 | -0.573 | -0.531 | 0.000 | *** |
| CV2pheno | - | phenoPsychosis | -0.384 | -0.410 | -0.358 | 0.000 | *** |

*Notes*. CV1= First canonical variate. CV2=Second canonical variate. CV#prs= Canonical component for PRS scores. CV#pheno= Canonical component for phenotypes. * $p < .05$, ** $p < .01$, *** $p < .001$.

**Table 40.** Descriptive statistics

| | Mean | SD | min | max |
|---|---|---|---|---|
| Cumulative adversity | 1.44 | 1.87 | 0 | 15 |
| *Phenotypes* | | | | |
| ADHD | 2.66 | 3.00 | 0 | 14 |
| Anxiety | 2.16 | 2.48 | 0 | 17 |
| Depression | 1.32 | 2.03 | 0 | 19 |
| Psychosis | 5.79 | 10.24 | 0 | 104 |
| *PRS scores* | | | | |
| ADHD | -47.32 | 3.45 | -61.36 | -34.97 |
| Anxiety | -37.46 | 3.12 | -48.51 | -26.16 |
| Depression | -31.86 | 2.41 | -41.10 | -23.78 |
| Psychosis | -22.87 | 2.28 | -31.99 | -14.34 |

*Notes*. Measures taken at baseline when participants were aged 9-10. The cumulative adversity score was obtained by summing all adversity items.
